# NON-AVIAN DINOSAUR EGGSHELL CALCITE CONTAINS ANCIENT, ENDOGENOUS AMINO ACIDS

**DOI:** 10.1101/2020.06.02.129999

**Authors:** Evan T. Saitta, Jakob Vinther, Molly K. Crisp, Geoffrey D. Abbott, Thomas G. Kaye, Michael Pittman, Ian Bull, Ian Fletcher, Xinqi Chen, Matthew J. Collins, Jorune Sakalauskaite, Meaghan Mackie, Federica Dal Bello, Marc R. Dickinson, Mark A. Stevenson, Paul Donohoe, Philipp R. Heck, Beatrice Demarchi, Kirsty E. H. Penkman

## Abstract

Rates of peptide bond hydrolysis and other diagenetic reactions are not favourable for Mesozoic protein survival. Proteins hydrolyse into peptide fragments and free amino acids that, in open systems such as bone, can leach from the specimen and be further degraded. However, closed systems are more likely to retain degradation products derived from endogenous proteins. Amino acid racemisation data in experimental and subfossil material suggests that mollusc shell and avian eggshell calcite crystals can demonstrate closed system behaviour, retaining endogenous amino acids. Here, high-performance liquid chromatography reveals that the intra-crystalline fraction of Late Cretaceous (estimated ~80 Ma) titanosaur sauropod eggshell is enriched in some of the most stable amino acids (Glx, Gly, Ala, and possibly Val) and those that racemise are fully racemic, despite being some of the slowest racemising amino acids. These results are consistent with degradation trends deduced from modern, thermally matured, sub-fossil, and ~3.8 Ma avian eggshell, as well as ~30 Ma calcitic mollusc opercula. Selective preservation of certain fully racemic amino acids, which do not racemise in-chain, along with similar concentrations of free versus total hydrolysable amino acids, likely suggests complete hydrolysis of original peptides. Liquid chromatography-tandem mass spectrometry supports this hypothesis by failing to detect any non-contamination peptide sequences from the Mesozoic eggshell. Pyrolysis-gas chromatography-mass spectrometry reveals pyrolysates consistent with amino acids as well as aliphatic hydrocarbon homologues that are not present in modern eggshell, suggestive of kerogen formation deriving from eggshell lipids. Raman spectroscopy yields bands consistent with various organic molecules, possibly including N-bearing molecules or geopolymers. These closed-system amino acids are possibly the most thoroughly supported non-avian dinosaur endogenous protein-derived constituents, at least those that have not undergone oxidative condensation with other classes of biomolecules. Biocrystal matrices can help preserve mobile organic molecules by trapping them (perhaps with the assistance of resistant organic polymers), but trapped organics are nevertheless prone to diagenetic degradation even if such reactions might be slowed in exceptional circumstances. The evidence for complete hydrolysis and degradation of most amino acids in the eggshell raises concern about the validity of reported polypeptide sequences from open-system non-avian dinosaur bone and other Mesozoic fossils.

## Introduction

Some biomolecules are highly stable and can survive deep into the geologic record with minimal alteration (Briggs & Summons 2014). Steroids (Melendez *et al.* 2013) and pigments, such as porphyrins (Greenwalt *et al.* 2013) and melanin (Glass *et al.* 2012), are good examples of diagenetically recalcitrant biomolecules. Simple hydrocarbon backbones forming aliphatic (e.g., kerogen) or ring structures preserve deep into the geologic record, but more complex molecular structures with heteroatomic backbones are less stable (Eglinton & Logan 1991). Biomacromolecules that form from the organised condensation of monomers into ordered polymers based upon the genetic code are particularly unstable, as they can irreversibly hydrolyse to their constituent monomers. However, these relatively unstable biomacromolecules (e.g., nucleic acids and proteins) are of the highest biological interest since they serve critical, complex functions in organisms and changes in their sequence and structure can provide insight into evolution, physiology, and ecology (e.g., Leonard *et al.* 2002).

Ancient DNA has been recovered from a 560–780 Ka horse bone retrieved from permafrost, nearing the expected upper limit of DNA survival in nature (Orlando *et al.* 2013). In contrast, claims of preserved older DNA, including Mesozoic DNA, have been strongly refuted (Allard *et al.* 1995; Hedges *et al.* 1995; Zischler *et al.* 1995; Poinar & Cooper 2000). Some of the oldest partially intact proteins from bone come from ~3.4 Ma bone from the high arctic (Rybczynski *et al.* 2013), with their preservation likely due to the exceptionally cold burial environment; kinetically, such peptides have very young thermal ages (Demarchi *et al.* 2016). However, controversial claims of preserved protein in bone as old as the Early Jurassic have been published (e.g., Reisz *et al.* 2013), and their low latitude and extreme geologic age (taking diagenetic geothermal heating into consideration) would place their thermal age orders of magnitude older than the reports from arctic sites (Demarchi *et al.* 2016).

One difficulty in searching for ancient proteins comes from environmental and laboratory contamination (Buckley *et al.* 2008, 2017; Bern *et al.* 2009). There is mixed evidence that amber might act to trap some ancient amino acids, but the amino acid profiles and racemization patterns are not what would be expected from a closed system of ancient amino acids (McCoy *et al.* 2019; Barthel *et al.* 2020). Examining proteins deposited within biomineralised calcium carbonate (intra-crystalline) helps to mitigate this issue. In contrast to open-system bone (Bada *et al.* 1999; Reznikov *et al.* 2018; Saitta *et al.* 2019), calcium carbonate (e.g., mollusc shells [Penkman *et al.* 2008, 2013] and avian eggshells [Brooks *et al.* 1990; Crisp *et al.* 2013]) can in some instances act as a closed system for amino acids (Towe & Thompson 1972; Towe 1980; Collins & Riley 2000). Note that this system is closed at the molecular scale so that eggshell respiratory pores, which are many orders of magnitude larger, do not influence this property, in the same manner that eggshell thickness or curvature are irrelevant to amino acid retainment. Calcite is thermodynamically more stable than aragonite, which tends to recrystallise as calcite during fossilisation (Benton 2001), potentially opening the system (Wehmiller *et al.* 1976; Harmon *et al.* 1983; Hearty & Aharon 1988; Hoang & Hearty 1989; Penkman *et al.* 2007, 2010).

The oldest well-supported closed system amino acids (i.e., not necessarily within a peptide chain) detected from the more thermostable calcite form of CaCO_3_ are from ~30 Ma mollusc opercula (Penkman *et al.* 2013). Reports of extremely ancient fossil amino acids exist, particularly in early literature, with the thermally stable amino acids Glu, Ala, and Val reported from a ~360 Ma trilobite (Abelson 1954), which have calcite in their cuticle (Dalingwater 1973) and eye lenses *in vivo* (Towe 1973; although see counter by Lindgren *et al.* 2019). However, trilobite protein diagenesis patterns and calcite system behaviour are uncharacterised; in contrast, eggshell and mollusc opercula have recent fossil records and modern tissues for thermal maturation experiments. The report also appears alongside that of a similar amino acid profile in open-system Jurassic *Stegosaurus* bone (Abelson 1954) whose amino acids might be exogenous (Saitta *et al.* 2019; Liang *et al.* 2020), urging re-examination of that study.

Although calcite can act as a closed system for peptides and amino acids, degradation of the trapped organics still proceeds. For example, in a survey of calcitic brachiopod shell, immunochemical signatures of modern shell peptides disappear by about 2 Ma (Curry *et al.* 1991; Walton 1998; Collins *et al.* 2003). Peptide fragmentation, amino acid profiles, and racemisation patterns have been thoroughly studied in modern, sub-fossil, and 3.8 Ma avian eggshell and compared to experimentally matured avian eggshell (Crisp 2013; Crisp *et al.* 2013; Demarchi *et al.* 2016). As eggshell peptides degrade over time and under higher environmental/experimental temperatures, D/L values along with relative concentrations of Glx, Gly, and Ala increase, while concentrations of Asx and Ser decrease. Among a consistent pattern of peptide degradation observed through a suite of eggshell samples, the oldest independently authenticated peptide fragment is an otherwise unstable, short, acidic region of the struthiocalcin protein preserved in ~3.8 Ma low-latitude ratite calcite eggshell (Demarchi *et al.* 2016). Despite its warm burial history, the high binding energy of this region of the peptide to calcite results in a unique ‘molecular refrigeration’ mechanism that drops the effective temperature around the peptide by ~30 K, reducing rates of hydrolysis (thermal age equivalent to ~16 Ma at 10 °C).

Non-avian dinosaur eggshell also consisted of calcite, with a similar structural organisation to avian eggshell, and can be found in large quantities at certain nesting sites, such as the Late Cretaceous (Campanian, ~83.5–79.5 Ma) titanosaur eggshell site from the Anacleto Member of the Río Colorado Formation in Auca Mahuevo, Argentina (Chiappe *et al.* 1998, 2000, 2003, 2005; Dingus *et al.* 2000; Grellet-Tinner *et al.* 2004; Garrido 2010). Sedimentological descriptions noted that those eggs contacted a sandstone layer below them while entombed by mudstone, indicating that they were laid on the surface of sandy depressions and subsequently buried by flooding (Chiappe *et al.* 2003, 2005). Non-avian dinosaur eggshell is known to contain endogenous biomolecules such as stable porphyrin pigments (Wiemann *et al.* 2017). More extreme claims of biomolecular preservation have been proposed in Auca Mahuevo eggshells, where immunochemistry was used as evidence for intact protein or protein-derived organics across the eggshell cross-section, including inter-crystalline regions considered to be outside of the closed system calcite crystals (Schweitzer *et al.* 2005). However, using immunochemistry to detect proteins in fossils has been suspected to yield false positives (Saitta & Vinther 2019). Therefore, using a variety of analytical techniques, this study aims to test the potential for preservation of original amino acids (and ultimately peptide sequence information) from Mesozoic calcite eggshell.

## Materials and Methods

To explore the potential for preservation of peptide sequences from dinosaur eggshell, we have analysed two independently obtained Late Cretaceous Argentine titanosaur eggshells (referred to as A and B) alongside modern chicken (*Gallus gallus domesticus*) and ostrich (*Struthio camelus*) eggshells as comparisons. In order to gather a range of evidence (i.e., triangulation), we used complementary analytical techniques to investigate the potential for amino acid and peptide sequence preservation (Table 1). We used light microscopy, laser stimulated fluorescence (LSF) imaging, reversed-phase high performance liquid chromatography (RP-HPLC), liquid chromatography coupled to tandem mass spectrometry (LC-MS/MS), pyrolysis-gas chromatography-mass spectrometry (Py-GC-MS), and Raman spectroscopy (along with attempts at time-of-flight secondary ion mass spectrometry [TOF-SIMS] [supplemental material]). Samples were prepared (e.g., cracked, powdered, resin-embedded thin sectioned, or polished) as needed, including a bleach treatment that allows for isolation of intra-crystalline amino acids. Modern, thermally matured (300 °C, 120 hr), and ≤ 151 Ka ratite eggshell data from Crisp (2013), run on the same RP-HPLC equipment and in the same laboratory as the samples described here, was used for further comparison. See the supplemental material for details of these fossil/modern samples and methods.

**Table 1.** Methodological triangulation employed in this study.

| Technique | Signal detected | Insight into protein preservation | Example reference |
| --- | --- | --- | --- |
| Light microscopy | Plane or crossed polarized, transmitted or reflected light | Integrity of calcite crystal structure (i.e., system dynamics); localization of dark organic material | Hirsch & Quinn 1990 |
| LSF | Fluoresced light | Localization of non-fluorescing organic material | Kaye *et al.* 2015 |
| RP-HPLC | 13 primary amino acids in their relevant chiral forms | Amino acid concentration, composition, racemization extent, and hydrolysis extent; endogenicity of amino acids and any preserved peptides | Crisp *et al.* 2013 |
| LC-MS/MS | Peptide sequences | Endogenicity of any recovered peptides; if endogenous, evolutionary information | Demarchi *et al.* 2016 |
| Py-GC-MS | Pyrolysis decomposition products of molecules | Presence of molecules consistent with amino acids, proteins, or organic geopolymers | Saitta *et al.* 2017 |
| Raman spectroscopy | Raman-active molecular vibrations | Presence and localization of molecules consistent with amino acids, proteins, or organic geopolymers | Wiemann *et al.* 2018 |
| TOF-SIMS | Secondary ions from fragmented molecules | Presence and localization of molecules consistent with amino acids, proteins, or organic geopolymers | Orlando *et al.* 2013 |

## Results

One of the very first observations was that, upon powdering and polishing, the titanosaur eggshells A and B released a fairly strong odour reminiscent of a mixture of burnt hair and petrol.

### Light microscopy and LSF imaging

The titanosaur eggshell A fragment has a lightly coloured interior and exterior surface, and the exterior surface is covered in small, round ornamentation (Fig. 1A, C). The interior cross-section of the eggshell shows large regions of black calcite whose structure has been lost, however, there is a band of lightly coloured calcite deep in the interior of the eggshell cross-section (Fig. 1B, E). The black, astructural calcite does not fluoresce, while the lightly coloured calcite fluoresces pale white/yellow and the infilling material, likely from the sediment matrix, between ornaments and calcite crystal units within the pore spaces fluoresces blue (Fig. 1 D, F). About half of the calcite in the eggshell appears to be black and astructural, lacking their characteristic morphology (as in Chiappe *et al.* 1998, 2000, 2003, 2005; Grellet-Tinner *et al.* 2004).

**Figure 1.**
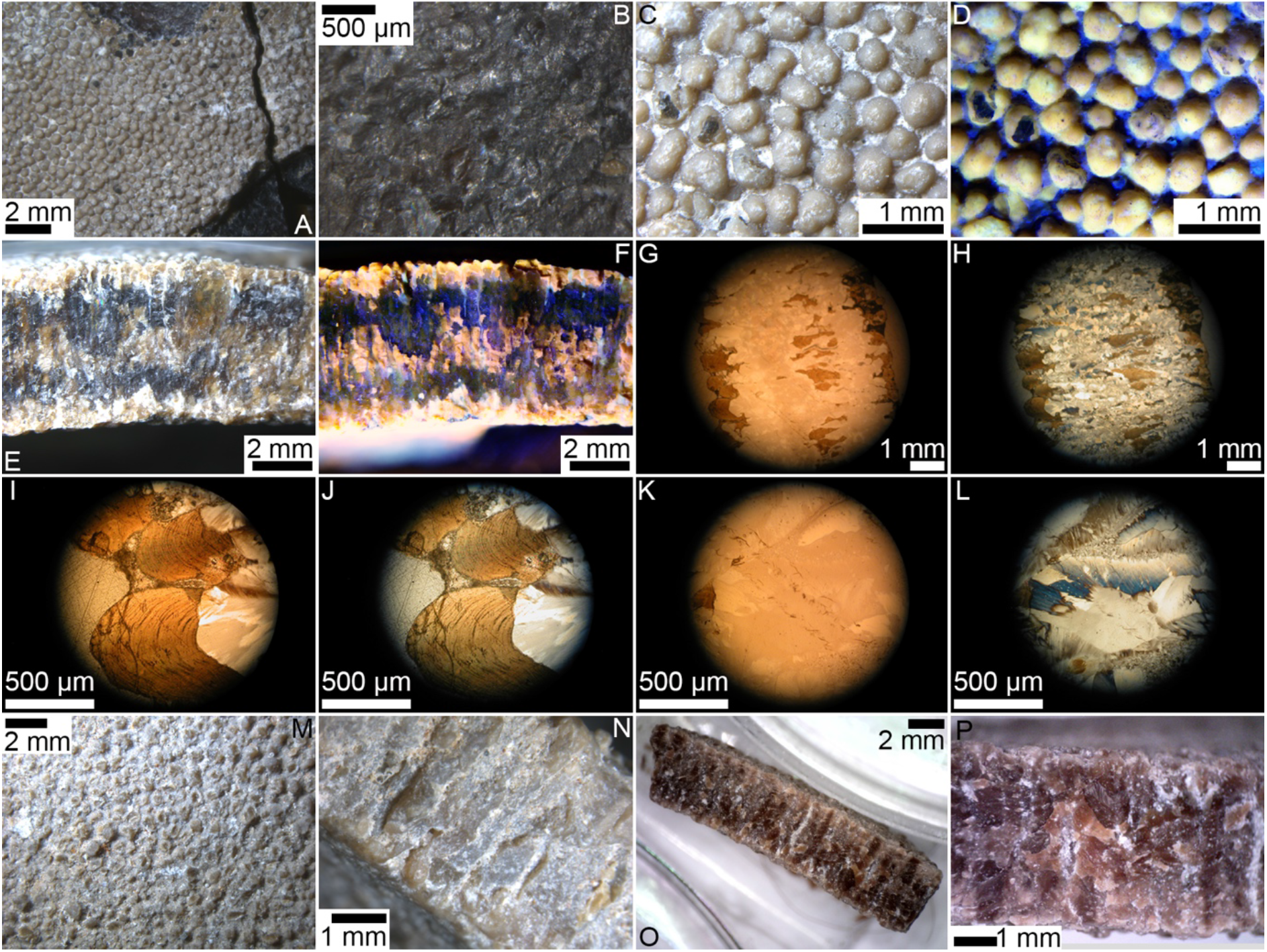
Titanosaur eggshell analysed in this study under light microscopy. A–L, titanosaur eggshell A. A, large fragment of titanosaur eggshell A viewed from the exterior surface showing ornamentation as well as some underlying black, amorphous calcite revealed when surface layers flaked off during splitting with a pestle. B, amorphous, black calcite viewed from exterior that was exposed. Exterior surface ornamentation under white light, C, and LSF, D. Cross section through the entire eggshell with the exterior surface to the top of the panel under white light, E, and LSF, F. Thin section of entire eggshell cross-section with exterior surface to the left of the panel under plane, G, and crossed, H, polarised light. Thin section of eggshell exterior ornamentation with the exterior surface to the left of the panel under plane, I, and crossed, J, polarised light. Thin section of recrystallised interior calcite under plane, K, and crossed, L, polarised light. M–P, titanosaur eggshell B. M, titanosaur eggshell B view from the exterior surface showing ornamentation. N, a weathered edge of the eggshell revealing palisade/column crystals. O–P, freshly broken edge of the eggshell showing brown staining of the calcite crystals with the exterior surface to the top of the panel.

Thin sections reveal highly organised, light brown calcite with some original prismatic external layer and ornamentation, palisade/column layer, or mammillary cone layer morphology (as in Chiappe *et al.* 1998, 2000, 2003, 2005; Grellet-Tinner *et al.* 2004) when observed under plane and crossed polarised light, correlating to the lightly coloured regions observed in the non-thin-sectioned fragment (Fig. 1G–J). Much of the palisade/column layer structure has been lost, more so than the other layers. The dark regions in the non-thin-sectioned fragment are clear under plane polarised light and have a disorganised white and blue refraction pattern under crossed polarised light without any original morphology (Fig. 1G–H, K–L) and are recrystallized. Sediment infilling between adjacent external ornamentation is apparent in the thin sections.

Titanosaur eggshell B shows a similar external and internal structure to titanosaur eggshell A, such as the presence of ornamentation on the exterior surface (Fig. 1M). The internal palisade column crystals appear to be more recognizable in titanosaur eggshell B than in titanosaur eggshell A, and the titanosaur eggshell B be shows a less stratified pattern of dark staining (Fig. 1N–P).

### RP-HPLC amino acid analysis

The titanosaur eggshell samples had a consistent total hydrolysable amino acid (THAA) compositional profile that matches those from old and/or thermally mature eggshell (Fig. 2A–B) and Eocene mollusc opercula (Penkman *et al.* 2013), being enriched in stable Glx, Gly, and Ala while being depleted in other amino acids, particularly unstable Asx and Ser. Only Glx, Gly, Ala, and Val consistently appear in appreciable concentrations among the variously treated replicates of titanosaur eggshell. All titanosaur samples yielded similar profiles (supplemental material). The elevated baseline signal post-58 minutes in some of the chromatograms (commonly seen in very degraded organic samples [Crisp 2013]) means that data obtained after this time (e.g., on Val, Phe, Ile, and sometimes D-Tyr) are reduced in accuracy and should be interpreted cautiously.

**Figure 2.**
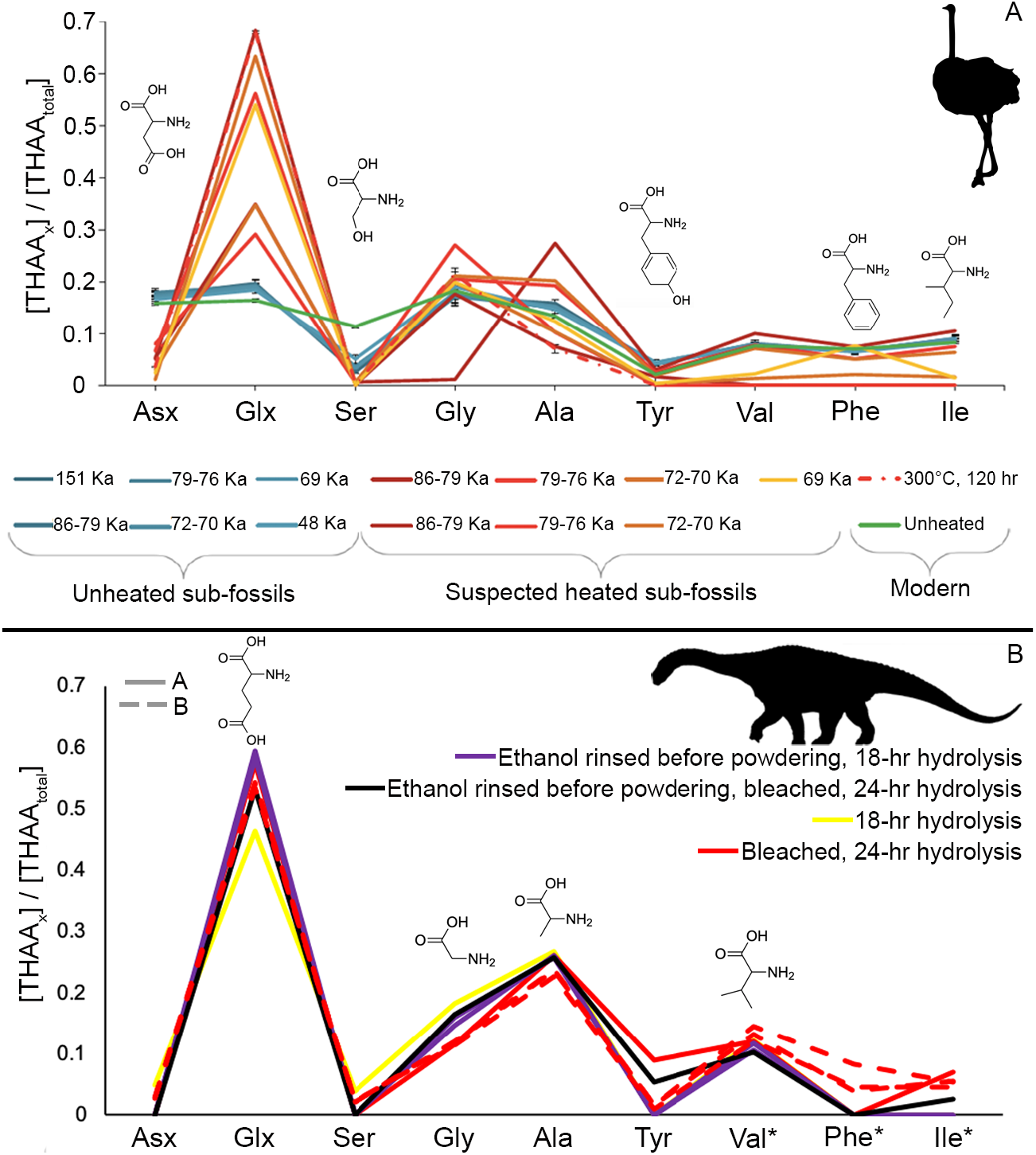
THAA compositional profiles of modern, experimental, and ancient eggshell. A, modern, thermally matured (300 °C, 120 hr), and ≤ 151 Ka ratite eggshell from Crisp (2013). All error bars (black) represent two standard deviations about the mean and are very narrow. The 86-79 Ka, suspected heated, sub-fossil eggshell sample with low Gly content is potentially as a result of inaccurate peak quantification. Panel, A, reproduced and modified from Figure 6.19 of Crisp (2013). B, Titanosaur eggshell A and B. *Data in titanosaur eggshell B from elution time > 58 min (e.g., Val, Phe, Ile) is of low accuracy due to elevated baseline values. Chemical structures are shown above each peak (only the deamidated forms of Asx and Glx are shown).

All amino acids present in the titanosaur eggshell capable of racemization show strong evidence of being fully racemic (i.e., high D/L values), despite low detected concentrations that can make calculating some of the D/L values challenging (Table 2; supplemental material). THAA and FAA yield similar D/L values and amino acid concentrations (e.g., Gly, Ala, Val concentrations, although errors can be large and lactam formation from the cyclisation of free Glx results in an underestimation in free Glx in this RP-HPLC method) (supplemental material). D/L values > 1 in Val result from statistical error as a result of low amino acid concentration rather than co-elution with another molecule; similar Val D/L values have also been reported in ancient ratite eggshell (Demarchi *et al.* 2016). S-allo-Ile co-elutes with some other molecule, evidenced by poorly resolved chromatography peaks for D-allo-Ile using RP-HPLC (Powell *et al.* 2013), and calculated Ile racemisation values are suspect (supplemental material). Regardless, Ile presence in the titanosaur eggshells is not strongly supported.

**Table 2.** Bleached titanosaur eggshell A and B amino acid racemisation and [Ser]/[Ala] values. NA indicates that amino acid concentration was below detection limit. *Data from elution time > 58 min is of low accuracy due to elevated baseline values.

| Treatment | Sample | Glx D/L | Ala D/L | Val D/L* | Asx D/L | Ser D/L | [Ser]/[Ala] |
| --- | --- | --- | --- | --- | --- | --- | --- |
| Bleached FAA | A | 1.03 | 0.93 | 1.22 | 0 | NA | 0 |
|  | B | 0.99 | 0.92 | 1.09 | 0.9 | 0.32 | 0.02 |
|  | B | 1.04 | 0.92 | 1.19 | 0.98 | 1.75 | 0.01 |
|  | B | 1.02 | 0.94 | 1.34 | 1.03 | 0.79 | 0 |
|  | B | 1.01 | 0.94 | 1.4 | 1.01 | 0.96 | 0.01 |
| Ethanol rinsed before powdering, bleached FAA | A | 1.05 | 0.97 | 1.11 | NA | NA | 0 |
| Bleached, 24-hr hydrolysis THAA | A | 1.04 | 0.96 | 1.29 | NA | NA | 0 |
|  | B | 0.99 | 0.67 | 1.47 | 0.23 | 0.1 | 0.1 |
|  | B | 0.99 | 0.7 | 1.16 | 0.23 | 0 | 0.09 |
|  | B | 0.99 | 0.7 | 0.93 | 0.23 | 0.03 | 0.09 |
| Ethanol rinsed before powdering, bleached, 24-hr hydrolysis THAA | A | 1.02 | 0.93 | 1.14 | NA | NA | 0 |

The [Ser]/[Ala] values in the titanosaur eggshell are very low, consistent with Ser degradation and Ala enrichment.

### LC-MS/MS

Seven peptides were detected by LC-MS/MS in the sample prepared in Turin (Table 3). Of these, three could be matched by PEAKS to protein sequences contained in the Aves_Reptilia database (namely, to histone H4 from *Gallus gallus* [supplemental material]). Of note, the peptide DNIQGITK matched to *Gallus gallus* histone H4 contains two potential deamidation sites, both of which were found to be unmodified, indicative of its modernity. The four peptide sequences not identified by PEAKS were further searched against UniProtKB_SwissProt using BLASTp and yielded matches to: human isoform 2 of Histone H2B type 2-F (sequence AMGIMNSFVNDIFER, 100% identity); KC19, human keratin (sequence SRSGGGGGGGLGSGGSIRSSY, 100% identity; also identified by PEAKS in the Copenhagen replicate); K2C4, human keratin (sequence LALDIEIATYR, 100% identity); human POTE ankyrin domain family member I (sequence AGFAGDDAPR, 100% identity).

**Table 3.**
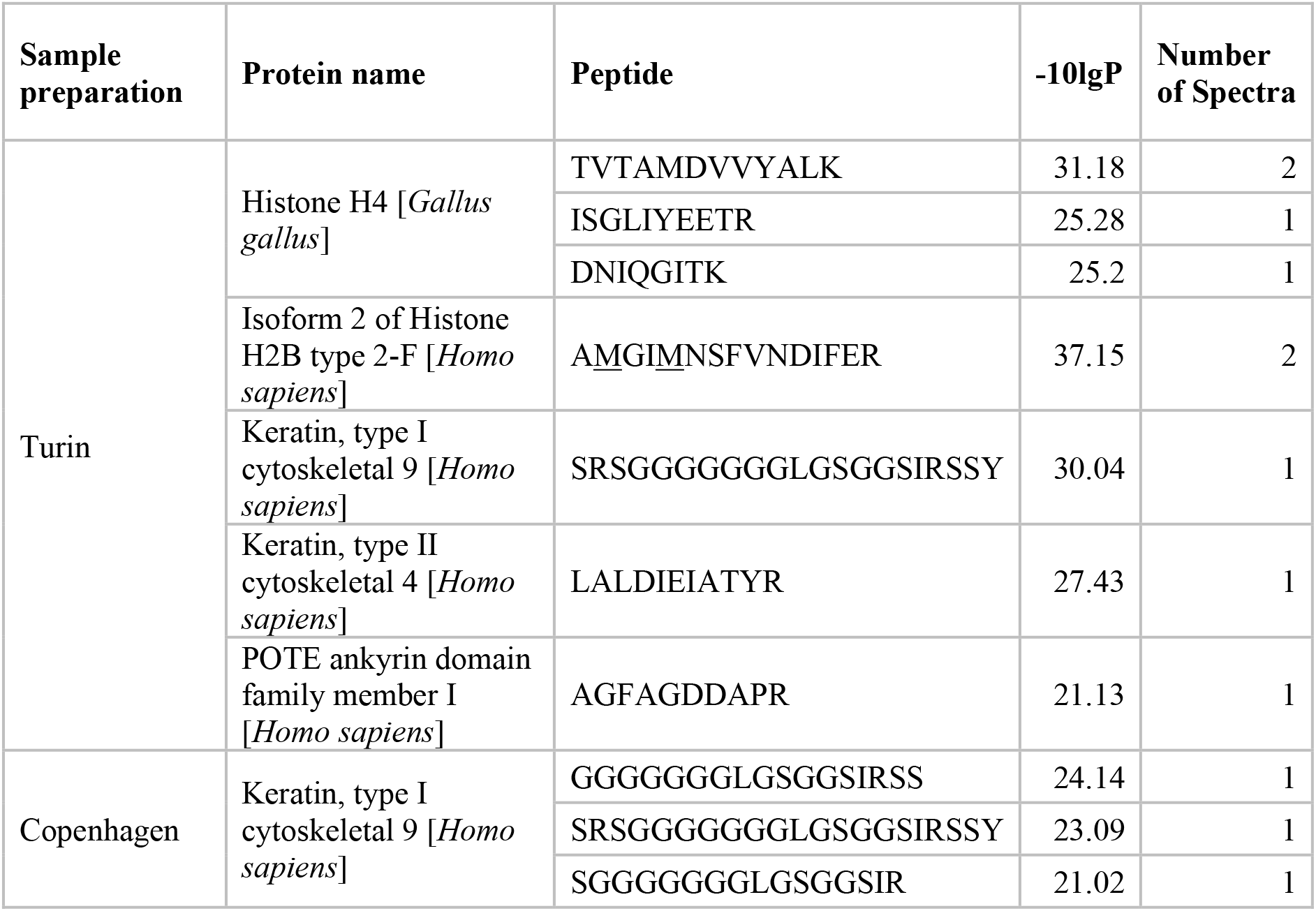
Peptides detected in the bleached titanosaur eggshell A (Turin and Copenhagen replicates) and their significant matches to known proteins. Note that the asparagine and glutamine are undeamidated in peptide DNIQGITK (Histone 4), supporting its modern origins. Underlined methionines are oxidised.

All *de novo* peptides (i.e., unmatched sequences reconstructed by PEAKS, with peptide scores −10lgP < 20; Table S.5), were also searched by BLASTp against UniProtKB_SwissProt and only two (ESYSVYVYK and LAAAARFMAW) yielded significant matches (to macaque Histone H2B and an uncharacterized protein from an Ascomycete, respectively).

The procedural blank (prepared in the same Turin laboratory as the dinosaur eggshell) contained four peptides of human albumin, two tubulin peptides, one highly conserved fragment and one potential histone peptide (DNLQGITK, also found in the eggshell sample; Table S.6); the “wash” water blank analysed before the eggshell sample contained a range of sequences, including four histone peptides (also AMGIMNSFVNDIFER found in the eggshell sample).

The sample prepared in the aDNA facilities in Copenhagen was cleaner than that prepared in Turin: it yielded three peptide sequences, all identified as human keratin (Table 3), and three *de novo* sequences which did not yield any matches to known proteins (Appendix, Table S.7). The Copenhagen procedural blank contained two peptides identified as human albumin and no peptides were found in the wash blank preceding the sample.

### Py-GC-MS

Examining the total ion chromatograms from Py-GC-MS of modern chicken and titanosaur eggshell A reveals how different decontamination methods can greatly affect results (supplemental material). This is particularly apparent in the titanosaur eggshell A samples, where more intensive decontamination decreased the organic content, evidenced by the more prominent column bleed at the end of the run and reduction of the intensity of some of the relatively later eluting peaks. Overall, it appears that organic content in the titanosaur eggshell A samples are lower than that in the modern chicken eggshell samples, evidenced by the prominence of the column bleed observed in the titanosaur eggshell A samples that was not observed in the modern chicken eggshell samples. However, minor variation in the mass of eggshell powder analysed could also influence this pattern, at least in part.

Comparing pyrolysates from the samples that had been DCM rinsed and Soxhlet extracted seems to be the most appropriate approach, given that these have been thoroughly decontaminated in a similar manner and were analysed on the same Py-GC-MS unit in close temporal proximity, making comparisons of retention times easier (Fig. 3A–B). With respect to lipids, titanosaur eggshell A pyrolysates contain *n*-alkanes/*n*-alkenes typical of kerogen (supplemental material), and these are also observable in the bleached (but not DCM rinsed and Soxhlet extracted) titanosaur eggshell A (supplemental material), while these are absent in modern chicken eggshell. Both titanosaur eggshell A and chicken eggshell contain simple pyrolysates with ring structures like toluene and phenols. The modern chicken eggshell contains several prominent nitrogen-bearing peaks such as nitriles, indoles, pyrrole, and pyridine, unlike the titanosaur eggshell A.

**Figure 3.**
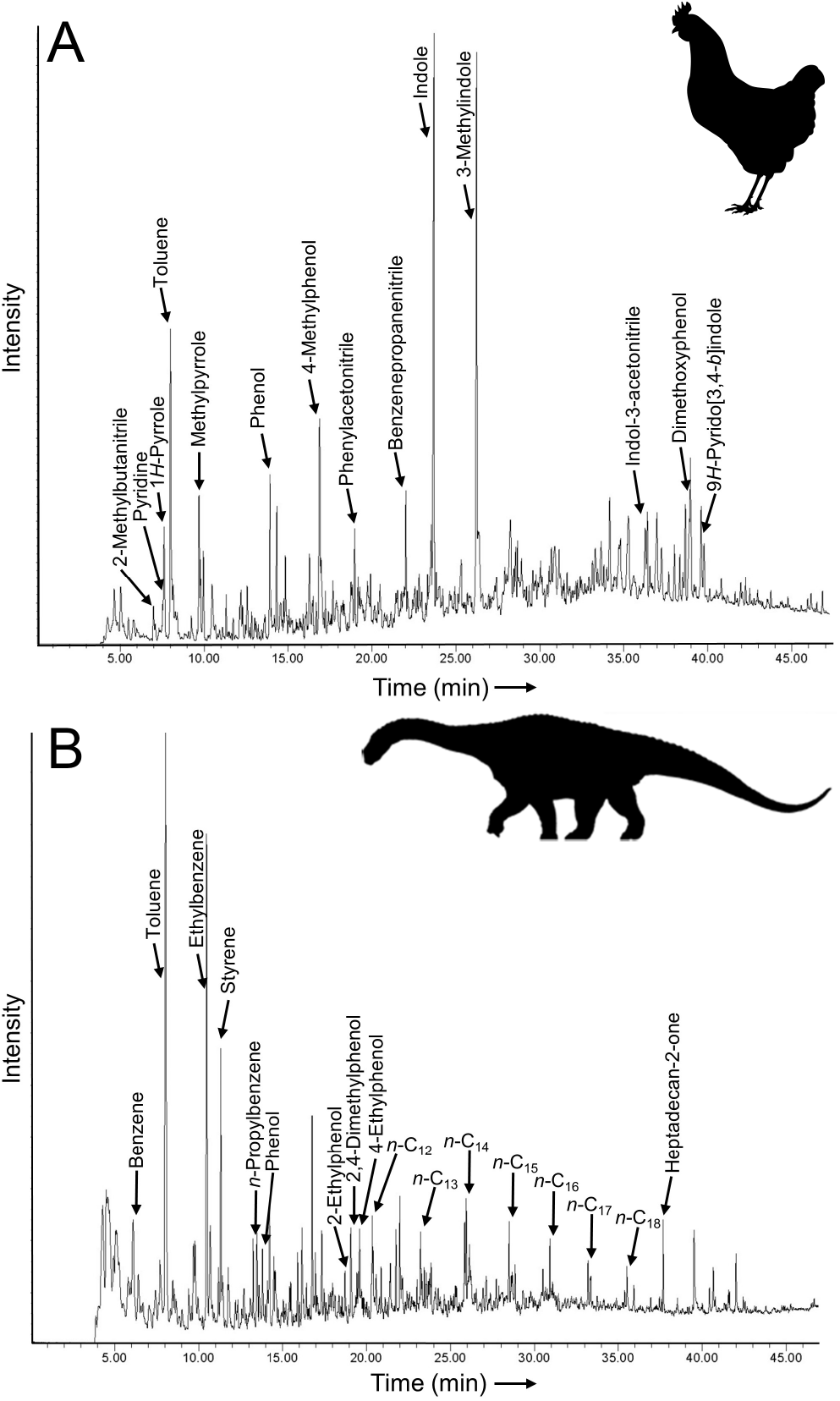
Comparison of identified pyrolysis products in, A, modern chicken (ethanol rinsed before powdering) and, B, the titanosaur eggshell A (not ethanol rinsed before powdering) eggshell after DCM rinsing and Soxhlet extraction.

### Raman Spectroscopy

The titanosaur eggshell A has two distinct chemical phases as revealed by Raman mapping (Fig. 4A–B; supplemental material). These phases correspond to 1) the light/non-recrystallized regions at the outer and inner surfaces, as well as the center of the eggshell’s cross section, and 2) the dark/recrystallized regions between these light regions. The light regions showed much greater fluorescence than the dark regions during Raman spectroscopy; this resulted in more noise and therefore the need to lower the excitation laser power relative to the analyses of the dark regions, making quantitative comparisons of spectral data between the two phases extremely difficult. However, both phases showed various peaks consistent with reference vibrations from calcite, quartz (likely from infilling sediment), and organic molecules, including potential non-cyclic, cyclic, and aromatic organic compounds (Fig. 4C). Some peaks are consistent with hydrocarbon vibrations with either absent or unspecified heteroatoms, while others are consistent with molecules containing O, N, S, or halogens.

**Figure 4.**
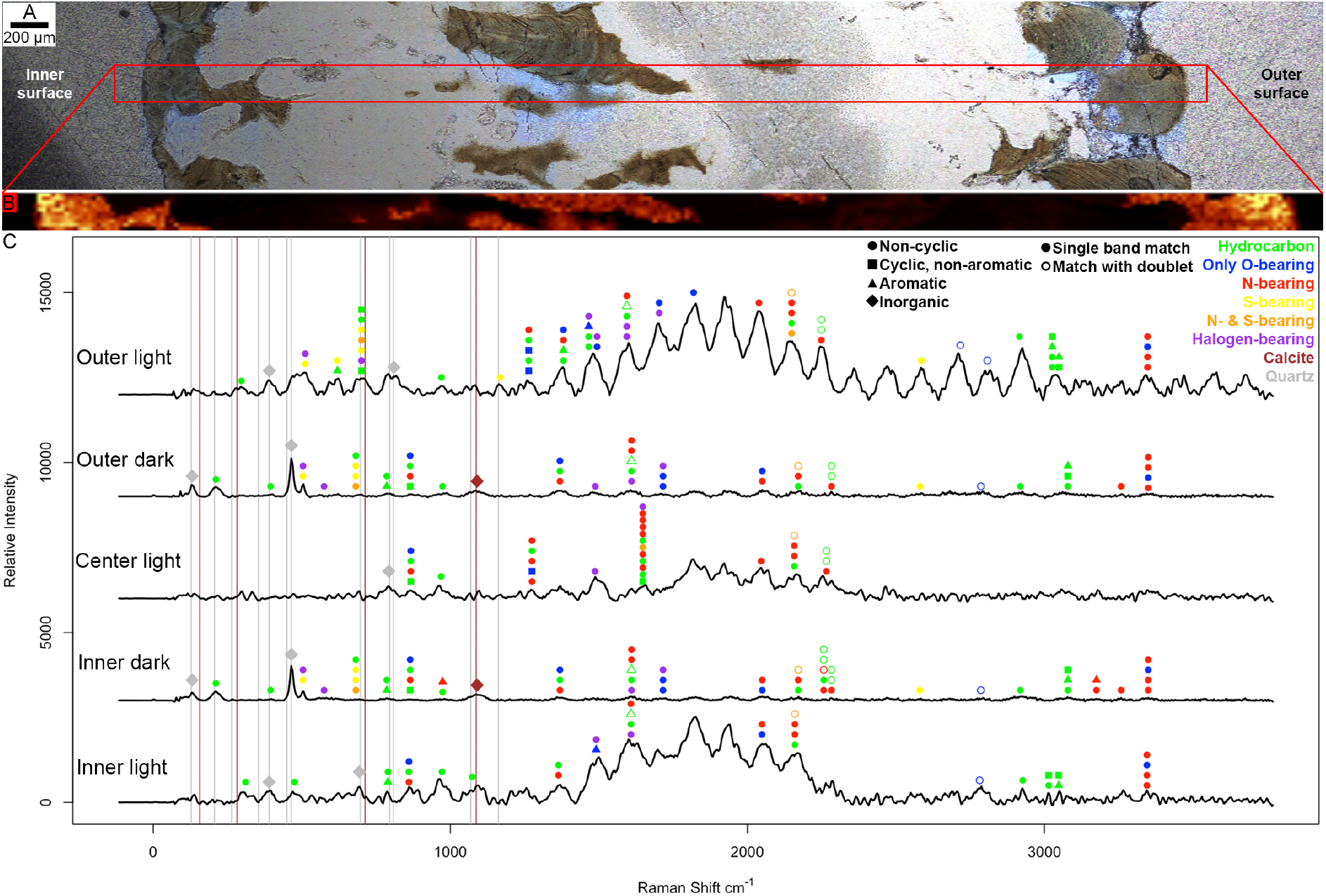
Raman spectroscopy of titanosaur eggshell A resin-embedded thin section. A, transmitted light micrograph with area mapped in red. Dark regions appear transparent, while light regions appear brown. B, Whole-spectrum map (i.e., all wavenumbers) under ~100 μW laser power. C, Spectra from the dark/recrystallized (20 mW laser power) and light/non-recrystallized (~100 μW laser power) regions. Fitted/deconvoluted peaks consistent with organic vibrations (Lin-Vien *et al.* 1991) indicated. Matches to reference vibrations that always have more than one band indicated by open symbols; in some cases, these potential matches might have lower support. Inorganic reference peak positions (Handbook of Raman Spectra for Geology, Laboratoire de Géologie de Lyon, Université de Lyon) shown with vertical lines, and closely matching peaks are coded as inorganic, overriding any potential organic matches. However, some potential organic peaks lie just beside these inorganic vibrations and might instead represent inorganic vibrations.

Modern ostrich eggshell showed calcite and organic peaks (with less noise than the titanosaur eggshell), including potential non-cyclic, cyclic, aromatic compounds, as well as hydrocarbons, O-, N-, S-, or even halogen-bearing molecules (supplemental material). The Raman spectrum outer (prismatic external) layer of the ostrich eggshell was noisier than those of the center column/palisade and inner mammillary cone layers.

## Discussion

Studies concluding protein preservation in fossils must take into account several aspects of this claim. Fossil proteins or protein-derived organics are those that have an appropriate *chemical signature*, *endogenicity* (McLoughlin 2011), and *antiquity*.

1. The composition of the organics must A) be consistent with protein or their degradation products generally (*chemical signature*) and B) should specifically be consistent with the composition expected from the *in vivo* proteins of the tissue or their degradation products (*chemical signature, endogenicity*).
2. The organics should be analysed for their degree of preservation (*antiquity*). Typically, older fossils would be expected to have greater degradation and alteration. Mechanisms explaining the observed degree of preservation must be supported (e.g., ‘molecular cooling’ of ~3.8 Ma eggshell peptide fragments; Demarchi *et al.* 2016).
3. The organics must localise in a manner that would be expected from endogenous protein sources as opposed to exogenous sources (*endogenicity*). The openness of the matrix any organics are fossilised in will dictate what patterns of organic influx or outflux are observed. Closed systems, as eggshell calcite can be, make interpreting these patterns far easier.

### 1) Composition of protein-derived material

A) The chemical signature of the titanosaur eggshell matches with that of organic, protein-derived material. There is a non-fluorescing (in bulk cross-section under LSF), black/brown colouration typical of organic material, as well as a release of organic volatiles upon powdering (as evidenced by the strong, peculiar odour); characterisation of similar volatile organic compounds by GC-MS supported the existence of a closed system in ~3.8 Ma ratite eggshell (Demarchi *et al.* 2016). The titanosaur eggshell A also yields organic pyrolysis products that are consistent with the presence of protein-derived material, such as toluene, benzenes, and phenols. Furthermore, Raman spectroscopy bands consistent with various organic molecules, including N-bearing molecules, are present throughout the titanosaur eggshell A cross section. RP-HPLC shows that amino acids are present within the titanosaur eggshell calcite.

B) As for the more precise nature of this organic signature consistent with protein-derived material, the THAA compositional profiles match those expected from old, thermally mature avian eggshell (i.e., Glx, Gly, and Ala enriched, but Asx and Ser depleted), unsurprising given that birds are dinosaurs and non-avian dinosaurs also produced calcitic eggs that could possibly have utilised similar mineralising proteins. Ala and Gly are decomposition products of Ser. In heating experiments (Vallentyne 1964) and fossils (Walton 1998), Ala, Val, and Glu had the longest half-lives, Glu being further stabilised by condensation to form pyroglutamic acid. Beyond thermal stability, acidic amino acids potentially play a role in eggshell mineralisation through involvement in Ca^2+^ binding (Marin *et al.* 2007) so it is perhaps unsurprising that Glx is found in high concentration in the titanosaur eggshell relative to other thermally stable amino acids.

However, the only significant matches of detected peptides in bleached titanosaur eggshell A all derive from likely contaminants. Keratins are expected to be common contaminating proteins in laboratory environments and can be introduced during sample handling, preparation, and/or analysis (Keller *et al.* 2008). Likely contaminating histone peptides have been identified in Mesozoic fossil bones in other studies; indeed, peptides TVTAMDVVYALK and ISGLIYEETR found in the titanosaur eggshell (Turin replicate) have also been reported by Schweitzer et al. (2013) and Cleland et al. (2015). The conserved nature of the histone protein sequence, the lack of Asn and Gln deamidation, the presence of histone sequences in the water blank analysed immediately before the eggshell sample and in the procedural blank, as well as their absence in the eggshell replicate prepared in a clean (ancient DNA) laboratory setting in Copenhagen, strongly support the exogenous origin of histone peptides (Table S.6). The *de novo* peptides reconstructed by PEAKS can also be considered insignificant, as they were derived from single spectra with low scores. Additional, broader BLAST searches yielded matches of these *de novo* peptides to *Macaca* and fungal sequences (Table S.5) phylogenetically distant to dinosaurs. Therefore, although there is evidence for original amino acids within the titanosaur eggshell, it is not possible to retrieve peptide sequence data.

### 2) Preservation of protein-derived material

As only four amino acids (Glx, Gly, Ala, and possibly Val) show clear, consistent evidence of survival in all of the variously treated titanosaur eggshell THAA and FAA replicates (supplemental material), consistent with known half-lives and decomposition products (Vallentyne 1964), this is strongly suggestive of significant peptide bond hydrolysis and subsequent degradation of less stable amino acids. These amino acids tend to be thermally resistant/stable over deep time in avian eggshell (Crisp 2013; Crisp *et al.* 2013) and simple in structure (e.g., Gly, Ala, Val). They are the only amino acids unequivocally present in the titanosaur eggshells and are in low concentrations (supplemental material), indicative of long-term diagenesis. Ala and Val have hydrophobic side chains, and insolubility might further enhance their preservation. Ser does not appear to be present in the titanosaur eggshell, and this amino acid is one of the least thermally stable, with the degradation of Ser contributing to Ala enrichment (Vallentyne 1964) in ancient or thermally mature eggshell. The amino acid compositional profiles from ~30 Ma mollusc shell (Penkman *et al.* 2013) show similarities to those detected in the titanosaur eggshell, despite presumably different profiles of the original proteins, suggesting that amino acid thermal stability is key to preservation. Given such a decrease in the amino acid types, long phylogenetically informative peptides are not expected – analogous to taking a novel and selectively removing all but five letters. Paragraphs, sentences, and words would be lost in the process. Furthermore, relatively little comparative literary criticism would be expected merely by comparing novels by their relative frequency of these remaining five letters.

The amino acids in the titanosaur eggshell are all fully racemic, suggesting that they are very ancient. Furthermore, the amino acids detected in the titanosaur eggshell are among the slowest racemising and most stable amino acids (Smith & Evans 1980; Crisp *et al.* 2013). Since relative racemisation rates between different amino acids are consistent over a range of temperatures (Crisp *et al.* 2013), any endogenous amino acids are likely fully racemic regardless of the titanosaur eggshells’ burial temperatures. Most amino acids can only racemise as free amino acids or N-terminally bound in-chain (Mitterer & Kriausakul 1984), with the exception of Ser (Demarchi *et al.* 2013a) and Asx (Stephenson & Clarke 1989) that can racemise internally bound in-chain; neither Ser or Asx are likely present in the titanosaur eggshell. The fully racemic mixtures observed in the titanosaur eggshell suggest that the amino acids derive largely from free amino acids or dipeptides in the form of cyclic dipeptides (e.g., diketopiperazines formed under thermal polymerisation from even racemized amino acid reactants [Hartmann *et al.* 1981]), abiotically condensed dipeptides (i.e., secondarily synthesized from previously free amino acids [Cleaves *et al.* 2009]), or the final remnants of the original peptide chain. However, abiotic dipeptide synthesis would require significant geothermal heat (Cleaves *et al.* 2009) and predicted rates of peptide hydrolysis are not supportive of original Mesozoic polypeptide survival (Nielsen-Marsh 2002). Gly, Ala, and Val in the titanosaur eggshell replicates show some consistency in having somewhat similar THAA and FAA concentrations, which would suggest high levels of peptide bond hydrolysis. Furthermore, similar D/L values retrieved from FAA and THAA suggests that very few peptide-bound L-amino acids persist. This similarity in THAA and FAA D/L values in the titanosaur eggshell is in contrast to younger proteinaceous samples whose FAA D/L values are greater than their THAA D/L values (Hare 1971; Smith & Evans 1980; Liardon & Lederman 1986), likely because most amino acids cannot readily racemise within a peptide chain (Hare 1971; Smith & Evans 1980; Liardon & Lederman 1986; Crisp *et al.* 2013). At low temperatures, such as would be expected during early taphonomic processes prior to any significant geothermal heating during diagenesis, hydrolysis is favoured over racemisation for many amino acids (Crisp *et al.* 2013; Demarchi *et al.* 2013b; Tomiak *et al.* 2013), meaning that the fully racemic amino acids detected here are likely indications of heavily hydrolysed proteins.

Detected Glx is predicted to be largely comprised of Glu since irreversible deamidation is a rapid degradation reaction, especially in acidic conditions (Hill 1965; Geiger & Clarke 1987; Wilson *et al.* 2012). Recrystallisation observed in the titanosaur eggshell A is consistent with past acidic conditions (Plummer *et al.* 1978). Additionally, given their role in eggshell mineralisation, one might also expect many acidic amino acids to be present prior to diagenetic alteration (Marin *et al.* 2007).

The apparently effectively complete hydrolytic cleavage of amino acids in the titanosaur eggshell A, compounded by the loss of most of the unstable amino acids, is further supported by the failure of LC-MS/MS to detect any significant, non-contaminant peptides. No homologous sequence to the highly stable region of struthiocalcin, as detected in ~3.8 Ma ratite eggshell (Demarchi *et al.* 2016), was detected. Of course, one would not necessarily expect a titanosaur to have a homolog to ratite struthiocalcin, given the vast evolutionary distance between them. However, struthiocalcin and related proteins are involved in eggshell mineralisation (Mann & Siedler 2004; Sánchez-Puig 2012; Ruiz-Arellano & Moreno 2014; Ruiz-Arellano *et al.* 2015) and make up ~20 % of the total organics in modern eggshell (Nys *et al.* 1999, 2004; Mann & Siedler 2004; Woodman 2012). If any endogenous peptides were to occur in the titanosaur, a similar negatively charged, Asp-rich sequence that binds tightly to calcite and has high preservation potential (Marin *et al.* 2007; Demarchi *et al.* 2016) might be a prime candidate. Importantly, most of the detected peptides in LC-MS/MS contain the labile amino acid Ser, as well as amide-bearing Asn and Gln residues. Since Asn and Gln are expected to undergo fairly rapid deamidation, even in-chain (Hill 1965; Geiger & Clarke 1987; Wilson *et al.* 2012), if such peptides were indeed Mesozoic, one might predict them to be fully converted into Asp and Glu.

Furthermore, while modern eggshell yields several prominent nitrogen-bearing pyrolysis products, the same is not true for the titanosaur eggshell A. This likely indicates a far higher proteinaceous concentration in modern eggshell and, conversely, high amounts of degradation of original proteins in the titanosaur eggshell, confirmed by the lower amino acid concentrations evident in the RP-HPLC data. Modern ostrich eggshell appeared to yield Raman vibrations with greater signal/noise ratio than the titanosaur eggshell A (i.e., cleaner spectra), even under the same excitation laser power (i.e., 20 mW). This greater noise is potentially consistent with relatively lower concentrations of organic molecules in the titanosaur eggshell A than in the ostrich eggshell, although fluorescence in the fossil sample during Raman spectroscopy can make such quantitative comparisons unreliable. On a related note, the presence of various Raman bands in the titanosaur eggshell A potentially consistent with halogen-bearing organic molecules possibly indicates bonding of exogenous halogens to endogenous organic geopolymers during diagenesis (Schöler & Keppler 2003). Significant diagenetic alteration of organics might also be supported by Raman bands in the titanosaur eggshell A consistent with S-bearing organic molecules, if indeed the S is exogenous and incorporated via sulfurization/vulcanization rather than deriving from the decomposition of endogenous S-bearing amino acids (which is also plausible).

Given the above evidence of significant protein degradation and diagenetic alteration of organic molecules it seems likely that the amino acids detected in the titanosaur eggshell have been fully hydrolysed, with further degradation through racemisation and loss of less stable amino acids.

### 3) Localisation of protein-derived material

It is apparent that a strong chemical signature for degraded, protein-derived organics is present in the titanosaur eggshell. However, the potential localisation patterns of these signatures was also investigated. Although there is no sedimentary matrix associated with the titanosaur eggshell specimens that can be analysed separately as a control (other than minor amounts of infilling in eggshell pores), due to the closed system behaviour of eggshells and other biocalcites, the oxidative bleach decontamination allows us to conclude that these amino acids are indeed intra-crystalline and therefore likely endogenous. However, the titanosaur eggshell calcite occurs in distinct layers with unique structure, and the potential for organic localisation within certain layers was also examined.

Modern avian eggshell has few organics in the outer crystal layer (Heredia *et al.* 2005), which could be consistent with the light coloration of the exterior of the titanosaur eggshells (although other regions were similarly light in color). Proteins are relatively abundant in the underlying palisade/column and mammillary cone layers of modern eggshells (Hincke *et al.* 1995; also see the THAA data within different eggshell layers in Demarchi *et al.* 2016). Thus, one might expect the titanosaur amino acids to be present in these more internal layers. The dark black/brown staining of the titanosaur eggshells is indeed most prominent in the central regions of the eggshell cross-sections.

Calcite’s birefringent, anisotropic optical properties allow for easy determination under crossed polarised light as to what portions of the titanosaur eggshell A cross-section have been recrystallised, altering their orientation and leading to a loss of original eggshell morphology. One might hypothesize that such recrystallisation could open the system, leading to a loss of endogenous amino acids. The recrystallised regions of the titanosaur eggshell A are those that also have black colouration – indicative of the presence of organics. It has been experimentally demonstrated in ostrich eggshell that calcite can maintain closed system behaviour with respect to their intra-crystalline proteins between pH 5 and pH 9, at least, without affecting protein degradation and amino acid racemisation (Crisp *et al.* 2013). Recrystallisation, if induced by pH fluctuations, might have occurred to a degree that resulted in a loss of original eggshell structure but maintained the closed system behaviour of intra-crystalline proteins without completely dissolving the calcite or inducing acid hydrolysis of any organic geopolymers possibly contributing to closed system behaviour (see following section).

Exogenous environmental amino acids might have been subsequently trapped in the recrystallised calcite. The amino acids are very ancient, so such re-entrapment would have to have occurred long ago. Given that recrystallisation could have occurred under significant diagenetic influence, the immediate burial environment might have been low in exogenous amino acids. Hypothetically, if exogenous amino acids were trapped late in diagenesis, the environmental THAA profile might be enriched in diagenetically stable amino acids. However, the THAA compositional profile of the titanosaur eggshell matches that predicted from ancient, thermally mature eggshell (i.e., ratios of Glx to Gly, Ala, and Val). The relatively high Glx concentration compared to moderate Gly and Ala concentrations in the titanosaur eggshell is better explained by eggshell protein precursors than diagenetic biases. Gly is the simplest amino acid and might be expected to occur in the highest concentration if amino acid compositional profiles contained solely a diagenetic signal. Open-system, Late Cretaceous dinosaur bone supporting an active microbiome can be heavily Gly dominated (Saitta *et al.* 2019) (although note that bone and eggshell amino acid composition differ *in vivo*, with high Gly content in bone). Furthermore, depending on the precise mechanism by which biocalcite crystals act as a closed system, re-entrapment of exogenous amino acids might be unlikely (see following section).

Raman spectroscopy revealed that both light and dark phases of the titanosaur eggshell A contained Raman vibrations consistent with various organic molecules, including N-bearing molecules. This further mitigates the concern that all of the amino acids are hypothetically deriving from exogenous amino acids trapped in the recrystallized regions of the eggshell. However, given differences in fluorescence between the two phases under Raman spectroscopy and the associated noise in the spectra, quantitative comparisons of the concentrations of organics between the two phases is ill-advised. As such, the hypothesis that the majority of the amino acids are associated with the dark regions of the eggshell remains open.

### Non-protein organics in eggshell through fossilisation

Modern eggshells contain organics other than proteins. In avian eggshells and other biocalcite, their closed system behaviour may be purely a result of the calcite crystals themselves or a combination of calcite and recalcitrant organics within the biomineral pores (Crisp *et al.* 2013).

Modern eggshells contain endogenous phospholipids (Simkiss & Tyler 1958). Kerogen-like aliphatic compounds can form taphonomically via *in situ* polymerisation of labile lipids (e.g., fatty acids from hydrolysed phospholipids) during decay and diagenesis (Stankiewicz *et al.* 2000; Gupta *et al.* 2006a, 2006b, 2007a, 2007b, 2008, 2009). Kerogen signatures were detected in the titanosaur eggshell A using Py-GC-MS under full scan mode, and these could have derived from endogenous phospholipids. Further analysis of the fossil eggshell kerogen using selected ion monitoring (SIM) scanning mode would allow for a useful comparison of carbon number between modern eggshell phospholipid fatty acid tails and the alkanes/alkenes detected in the fossil in order to estimate the extent of *in situ* polymerisation. Raman vibrations possibly from aliphatic organic compounds (e.g., hydrocarbons) were also detected in the titanosaur eggshell A, consistent with alkane/alkene geopolymers.

Furthermore, protein breakdown products can react with oxidised lipids through Maillard-like reactions to condense into stable, browning compounds referred to as N-heterocyclic polymers (Hidalgo *et al.* 1999; Wiemann *et al.* 2018). Raman bands in the titanosaur eggshell A consistent with cyclic N-bearing organic molecules could support the presence of such heterocyclic polymers. Therefore, kerogen and/or N-heterocyclic polymers contribute to the dark, organic colouration observed in the titanosaur eggshells. The possibility that these lipid-derived organic fossils help to trap endogenous amino acids should be investigated.

Polysaccharides are also present in the palisade/column and mammillary cone layers in modern avian eggshells (Baker & Balch 1962). Additionally, acid-mucopolysaccharide and protein complexes are present in eggshells (Simkiss & Tyler 1957). Melanoidins, condensation products formed from protein and polysaccharide degradation via Maillard reactions, can be present in fossils (Collins *et al.* 1992; Stankiewicz *et al.* 1997). Low molecular weight, aromatic structures comprise a significant portion of humic acids, formed through similar Maillard-like reactions (Hatcher *et al.* 1981; Hedges *et al.* 2000; Sutton & Sposito 2005). Therefore, the small, aromatic pyrolysis products detected in the titanosaur eggshell A may be evidence of melanoidins. These melanoidin or humic acid-like organics might also contribute to the black colouration in the titanosaur eggshells. Raman bands in the titanosaur eggshell A possibly from aromatic and/or N-bearing organic compounds are consistent with melanoidins. Melanoidins can be bleach resistant, although they can be degraded using acid hydrolysis (Hoering 1980; Namiki 1988; Wang *et al.* 2011). Therefore, the potential presence of melanoidins might help to protect amino acids in the titanosaur eggshell, shielding the so-called ‘intra-crystalline’ amino acid fraction from bleach oxidation but subsequently releasing them upon acid hydrolysis in the laboratory.

Kerogen can form early on in taphonomy during decay (Gupta *et al.* 2009) and humic acids can form in surface soils (Sutton & Sposito 2005). However, it is also possible that the dark-staining, non-protein organics in the titanosaur eggshell formed after long periods of time and through diagenesis during deep burial, possibly consistent with their localisation to the recrystallized calcite in titanosaur eggshell A (as evidenced by the dark colouration). Given rates of protein hydrolysis in eggshells (Crisp *et al.* 2013), it is reasonable to hypothesise that protein hydrolysis would typically occur before and contribute reactants to N-heterocyclic condensation products between amino acids and either sugars (i.e., producing melanoidins) or oxidised lipids. If recalcitrant organics like N-heterocyclic polymers or kerogen contribute to the retention of surviving endogenous amino acids, such a process might occur relatively early or late during the taphonomic process (i.e., at different points along the decomposition of proteins).

Based on the correlation between the black colouration and recrystallisation in the titanosaur eggshell A, one might wonder whether calcite dissolution promotes kerogen or N-heterocyclic polymer formation, freeing trapped reactants and allowing for them to mix more easily to ultimately condense into these resistant organic geopolymers. Experimental production of melanoidin can be done using Gly as a reactant, but subsequent acid hydrolysis of the melanoidin product yields <1 % Gly, suggesting that Gly is ultimately modified and becomes irretrievable upon incorporation into the polymer (Benzing-Purdie & Ripmeester 1983). This implies that the ancient amino acids detected in the titanosaur eggshell are indeed free and not secondarily released from covalent bonds from within a recalcitrant organic polymer. Therefore, the formation of N-heterocyclic polymers can lead to a loss of the endogenous amino acids that it incorporates, but can their recalcitrant nature (along with that of kerogen) then trap the remaining thermally stable amino acids? Such a protective capability might offset the likelihood of opening the system as a result of calcite dissolution and recrystallisation.

The extreme degree of organic degradation in the titanosaur eggshells demonstrated by the possible presence of kerogen or N-heterocyclic polymers, the degradation of the amino acids themselves, and other possible diagenetic signatures (e.g., calcite recrystallization or potential halogen-/S-bearing Raman vibrations) further testifies to the antiquity of the fossils.

### The future of analysing Mesozoic protein-derived material

Given observed and theoretical rates of hydrolysis, it seems highly unlikely for peptides to persist from the Mesozoic to the present without exceptional preservation mechanisms. With regard to hydrolysis, decreasing temperature is a key way by which to reduce the rate of thermodynamically favourable (i.e., inevitable) hydrolytic cleavage of peptides. However, the current polar ice caps have only existed on Earth for relatively limited periods of time and were not present during the Mesozoic (Holz 2015) and no fossilisation process or depositional environment has yet been reported that is anhydrous throughout the entirety of the taphonomic process.

If fully hydrolysed free amino acids (likely a subset of the original amino acid composition of the starting proteins) are the only proteinaceous remnants in Mesozoic fossils not subsequently condensed into a highly altered geopolymer, it limits their utility to provide peptide sequence information. However, the capacity of eggshell calcite to maintain a closed system deep into the fossil record, as suggested by the results here, indicates that a broader sampling in both number, locality, and age of Mesozoic eggshells will likely provide clearer insight into patterns of ancient amino acid preservation in this system.

Sub-fossil and fossil eggshell will help to calibrate experimental studies of organic degradation in closed systems. Short, intense thermal maturation experiments may sometimes be inappropriate to compare to specimens that have spent longer periods of time at relatively lower temperatures (Tomiak *et al.* 2013). For example, protein three-dimensional structure might affect rates of hydrolysis and racemisation (Collins *et al.* 1999) and denaturation can occur under elevated temperature more typical of experimental maturation than natural early taphonomic settings. The closed system experienced by intra-crystalline amino acids avoids confounding effects due to leaching of amino acids, pH changes, contamination, and microbial decay (Child *et al.* 1993; Walton 1998; Crisp *et al.* 2013), so a deep fossil record of eggshells allows for studying long-term protein degradation in completely natural closed system environments. However, for Mesozoic eggshell, it is reasonable to assume that some degree of diagenesis or even catagenesis will have taken place. For example, such geothermal gradients can expose the eggshell to or above temperatures of protein denaturation, i.e., 50–80 °C (Roos 1995). Therefore, thermal maturation remains a useful experimental tool for studying organic degradation in fossils of appreciable age and thermal maturity.

Very ancient amino acids might yield insights into palaeobiology in addition to organic geochemistry, potentially preserving taxonomic signatures in their amino acid profiles, as seen in calcium carbonate tissue such as brachiopod shells, modern to ~3 Ma mollusc shells, foraminifera tests up to 18 Ma, and modern to recently extinct avian eggshell (Miller *et al.* 2000, Hincke *et al.* 1995; Mann & Siedler 1999; Lakshminarayanan *et al.* 2002, 2003; Crisp *et al.* 2013; Jope 1967; Andrews *et al.* 1984; Kaufman *et al.* 1992; Demarchi *et al.* 2014; King & Hare 1972; Haugen *et al.* 1989). Such potential insight depends on the presence of sufficient variation in the original concentrations of stable amino acids of non-avian and avian dinosaur eggshells so as to be able to detect differences in original protein content after significant diagenesis and degradation. At the very least, endogenous, ancient amino acids and other fossil organics are good candidates for stable isotope analysis (e.g., C, O, or N) without the likelihood for incorporated environmental isotopes altering the observed ratios.

As far as pushing the upper age limit for well-supported amino acids, calcified eggshell represents a fairly limited fossil record. Examining the fossils of other calcite biominerals such as mollusc or brachiopod shells, and particularly trilobite cuticles and eye lenses, might provide opportunities to detect demonstrably ancient, endogenous amino acids throughout the Palaeozoic.

## Conclusions

The titanosaur eggshells show strong chemical evidence for the presence of highly stable ancient, endogenous amino acids. Although eggshell calcite is known to act as an extremely efficient closed system, these results are still about an order of magnitude older than the oldest reported eggshell amino acids and an estimated ~50 million years older than the oldest reported amino acids in biocalcite fossils for which there is good evidence, potentially making these titanosaur amino acids the oldest highly supported amino acids from a dinosaur. These results add further excitement for the potential for eggshell calcite to aid in the study of ancient organic degradation. As for their level of preservation, the amino acids appear to be predominantly hydrolysed; this has negative implications for the likelihood of highly preserved Mesozoic peptides and proteins, especially from open systems like bone or integument. The closed system phenomenon of eggshell calcite also highlights that there are two general aspects of molecular preservation in fossils: stability of the original molecule (e.g., against microbial/autolytic decay or diagenesis) and mobility of the molecule and its degradation products (e.g., solubility or the degree of openness of the matrix). However, the results here, although exciting, deserve to be replicated in other Mesozoic eggshell samples (and surrounding sediment matrix controls) alongside the addition of analyses (e.g., principal component) of large amino acid datasets in order to better characterise diagenetic patterns in these ancient eggshells.

## Supporting information

Raw Data

## Acknowledgments

We thank Jennika Greer (University of Chicago) for assistance in aseptic polishing of the eggshells that were not embedded in a matrix, Sheila Taylor (University of York) for assistance in preparing the samples for RP-HPLC, Erzsebet Thornberry (University of Bristol) for assistance in Py-GC-MS data analysis, Claudia Hildebrandt (University of Bristol) for assistance in polarized light microscopy, and Prof. Jesper Velgaard Olsen (Novo Nordisk Center for Protein Research, University of Copenhagen) for providing MS access and resources. Work supported by the University of Bristol Bob Savage Memorial Fund. *Gallus gallus domesticus* (Public Domain Dedication 1.0, https://creativecommons.org/publicdomain/zero/1.0/legalcode), *Struthio camelus* (credit: Lukasiniho, Creative Commons Attribution-NonCommercial-ShareAlike 3.0 Unported, https://creativecommons.org/licenses/by-nc-sa/3.0/legalcode, CC BY-NC-SA 3.0), and titanosaur (credit: T. Tischler, Creative Commons Attribution-ShareAlike 3.0 Unported, https://creativecommons.org/licenses/by-sa/3.0/legalcode, CC BY-SA 3.0) silhouettes were obtained from phylopic.org.

