## Appendix for "NON-AVIAN DINOSAUR EGGSHELL CALCITE CONTAINS ANCIENT, ENDOGENOUS AMINO ACIDS"

### SUPPORTING METHODS AND DATA: SAITTA *ET AL.* 2020

**MATERIALS:** A small piece of titanosaur eggshell had previously been purchased from a gem and mineral shop in the USA and was recently donated for research to the University of Bristol. The specimen is easily verifiable as a fossil non-avian dinosaur eggshell by its colour, structure, and texture, as well as exterior ornamentation and inner calcite structure that are consistent with descriptions of titanosaur eggshells such as those from Auca Mahuevo (Chiappe *et al.* 1998, 2000, 2003, 2005; Grellet-Tinner *et al.* 2004). Here, we simply refer to the specimen as titanosaur eggshell A.

A small second piece of titanosaur eggshell, referred to here as titanosaur eggshell B, was previously purchased from the public gift shop at the Natural History Museum of Denmark in Copenhagen (their sale has since ceased) and donated for research to the University of Bristol. This specimen is similarly consistent with titanosaur eggshell (e.g., from Auca Mahuevo) in colour, texture, and structure.

Both pieces were purchased roughly in the late 1990's to early 2000's, at the height of the international sale of Auca Mahuevo eggshell. Therefore, we are fairly confident that these titanosaur eggshell pieces are from Auca Mahuevo and highly confident that they are from the Late Cretaceous of Argentina. Such fragments have been, and continue to be, commonly sold throughout the world (often inappropriately labelled as "*Saltasaurus*" eggshell) from Patagonia, Argentina, despite regulations prohibiting their export. As such, the remaining fragments after destructive analyses of these two specimens will be repatriated to the Museo Carmen Funes in Argentina (Rodolpho Coria, pers. comm.). The fragments' compromised collection histories make them prime candidates for highly destructive exploratory analyses.

For comparison, chicken (*Gallus gallus domesticus*) eggs were purchased at a grocery store in Bristol, UK. Additionally, modern ostrich (*Struthio camelus*) eggshell was obtained from Oslinc ostrich farm in Lincolnshire, UK.

Modern, thermally matured (300 ºC, 120 hr), and ≤ 151 Ka ratite eggshell data from Crisp (2013), run on the same RP-HPLC equipment and in the same laboratory as the samples described here, was used for further comparison. Those samples were bleached according to the protocol used in this study (72 hr). Five samples with the least racemisation and producing no smell from each horizon of the study locality (Pinnacle Point, South Africa) were designated as unheated sub-fossils to produce error bars. Three samples of modern eggshell were used to produce error bars.

**RESIN-EMBEDDED THIN SECTIONING:** Petrolab Ltd. produced thin sections of titanosaur A and ostrich eggshell. Samples were embedded in epoxy resin. Polishing used diamond grit in water with the final polishing step using colloidal silica (~5 % ethylene glycol).

**LIGHT MICROSCOPY AND LSF IMAGING:** Light microscopy and LSF imaging were run at the University of Bristol. LSF images were collected using a modified version of the protocol of Kaye *et al.* (2015). Both titanosaur eggshell samples A and B were split into smaller fragments in order to expose fresh cross-sectional surfaces with a strong strike using a mortar and pestle that had been cleaned with 70 % ethanol.

Titanosaur eggshell A fragments were illuminated with a 405 nm, 500 mw violet laser diode and imaged with a Nikon D810 digital camera under magnification from a Leica M205 C stereomicroscope. An appropriate long pass blocking filter was fitted to the objective of the stereoscope to prevent image saturation by the laser. The laser diode was defocused to project a beam cone that evenly lit the specimen during the photo's time exposure in a dark room. The images were post-processed in Photoshop CC 2016 for sharpness, colour balance, and saturation. White light images were taken using the same microscope and camera setup for comparison.

Titanosaur eggshell B fragments were examined under white light using a Leica M205 C stereomicroscope as well as a Dino-Lite Edge AM73915MZTL (10X~140X; 5.0MP; USB 3.0) Digital Microscope.

Plane and crossed polarised light images of the resin-embedded thin sections were obtained using a Nikon Coolpix 5000 mounted on a Nikon Optiphot polarising microscope.

**RP-HPLC AMINO ACID ANALYSIS:** Reversed-phase HPLC was run at the University of York. The titanosaur eggshell A was initially split into two subsamples at the University of York: one was rinsed with 70 % ethanol prior to powdering, one was not rinsed prior to powdering. The eggshell subsamples were powdered in a cleaned mortar and pestle (70 % ethanol rinsed) and split further into two subsamples each: one for the analysis of the whole-shell protein, one for the analysis of the intra-crystalline fraction, following the methods of Crisp *et al.* (2013). For isolation of the intra-crystalline fraction, the powdered eggshell was weighed accurately into a sterile 2 mL microcentrifuge tube and 50 μL 12 % (w/v) NaOCl (VWR) was added. Samples were re-agitated every 24 hr, and the bleach removed after 72 hr. Samples were then rinsed five times in milliQ water, with a final rinse in HPLC-grade methanol (VWR) and then air-dried under a fume hood. Flakes from the outer eggshell of titanosaur eggshell A that separated naturally during the sampling process were also separately analyzed.

The powdered whole-shell and intra-crystalline fractions were then prepared for reversed-phase HPLC analysis of free amino acids (FAA) (~2–5 mg of powder) and THAA (~1–5 mg of powder) following the methods of Penkman *et al.* (2008). Unbleached samples underwent 18-hr rather than 24-hr hydrolysis due to the logistics of laboratory preparation at the time those samples were run.

Titanosaur eggshell B, as well as outer flakes from titanosaur eggshell A that separated during early splitting, were first powdered at the University of Bristol and then prepared at the University of York using slightly modified procedures of Penkman *et al*. (2008) to isolate the intra-crystalline protein by bleaching; a 1-week bleaching time was used. Two subsamples were then taken; one fraction was directly demineralised and the FAA analysed, and the second was treated to release the peptide-bound amino acids, thus yielding the THAA. Samples were analysed in duplicate by RP-HPLC, with standards and blanks run alongside samples. During preparative hydrolysis, both asparagine and glutamine undergo rapid irreversible deamination to aspartic acid and glutamic acid respectively (Hill 1965). It is therefore not possible to distinguish between the acidic amino acids and their derivatives and they are reported together as Asx and Glx respectively.

The DL ratios of aspartic acid/asparagine, glutamic acid/glutamine, serine, alanine and valine (D/L Asx, Glx, Ser, Ala, Val) as well as the [Ser]/[Ala] value were then assessed to provide an overall estimate of intra-crystalline protein decomposition (IcPD). In a closed system, the amino acid ratios of the FAA and the THAA subsamples should be highly correlated, enabling the recognition of compromised samples (e.g. Preece & Penkman 2005). The D/L of an amino acid will increase with increasing time, while the [Ser]/[Ala] value will decrease. Each amino acid racemises at different rates, and therefore is a useful indicator of protein diagenesis over different timescales.

*Note:* An oxidative bleach treatment is used to remove inter-crystalline (i.e., open system) amino acids and proteins (some of which may be exogenous contamination), in order to specifically analyse closed-system, intra-crystalline amino acids and proteins from avian eggshell (Crisp 2013; Crisp *et al.* 2013). It has been demonstrated that ~99 % of the total hydrolysable amino acids (THAA) are retained in the intra-crystalline fraction of avian eggshells are resistant to leaching, even at high temperature. This, combined with the predictable degradation kinetics of THAA, demonstrates the closed system behaviour of the isolated intra-crystalline fraction in eggshell (Crisp *et al.* 2013).

| Treatment | [Asx] | [Glx] | [Ser] | [L-Thr] | [L-His] | [Gly] | [L-Arg] | [Ala] | [Tyr] | [Val]* | [Phe]* | [Leu]* | [Ile]* |
|---|---|---|---|---|---|---|---|---|---|---|---|---|---|
| Bleached FAA | 112 | 237 | 0 | 80 | 0 | 182 | 0 | 515 | 96 | 197 | 12 | 0 | 102 |
| 18-hr hydrolysis THAA | 80 | 763 | 65 | 0 | 0 | 300 | 0 | 440 | 0 | 201 | 0 | 0 | 0 |
| Bleached, 24-hr hydrolysis THAA | 0 | 585 | 0 | 50 | 0 | 117 | 0 | 265 | 92 | 123 | 0 | 0 | 71 |
| Ethanol rinsed before powdering, bleached FAA | 0 | 80 | 0 | 75 | 0 | 69 | 0 | 182 | 30 | 62 | 0 | 0 | 16 |
| Ethanol rinsed before powdering, 18-hr hydrolysis THAA | 0 | 609 | 0 | 0 | 0 | 150 | 0 | 267 | 0 | 120 | 0 | 0 | 0 |
| Ethanol rinsed before powdering, 18-hr hydrolysis THAA | 0 | 593 | 0 | 0 | 0 | 163 | 0 | 263 | 0 | 107 | 0 | 0 | 0 |
| Ethanol rinsed before powdering, bleached, 24-hr hydrolysis THAA | 0 | 162 | 0 | 14 | 0 | 50 | 0 | 79 | 16 | 31 | 0 | 0 | 8 |
| Outer flakes, bleached FAA | 3 | 59 | 3 | 24 | 0 | 44 | 1 | 164 | 27 | 64 | 37 | 48 | 44 |
| Outer flakes, bleached FAA | 3 | 58 | 3 | 25 | 0 | 47 | 1 | 165 | 15 | 78 | 30 | 22 | 45 |
| Outer flakes, bleached FAA | 2 | 44 | 3 | 17 | 0 | 50 | 3 | 133 | 4 | 50 | 21 | 25 | 21 |
| Outer flakes, bleached FAA | 2 | 43 | 2 | 16 | 0 | 48 | 0 | 133 | 8 | 43 | 31 | 39 | 28 |
| Outer flakes, bleached, 24-hr hydrolysis THAA | 7 | 143 | 12 | 13 | 0 | 73 | 5 | 168 | 15 | 78 | 50 | 29 | 61 |
| Outer flakes, bleached, 24-hr hydrolysis THAA | 6 | 121 | 12 | 12 | 0 | 70 | 5 | 159 | 9 | 67 | 36 | 17 | 58 |
| Outer flakes, bleached, 24-hr hydrolysis THAA | 7 | 103 | 10 | 10 | 0 | 56 | 5 | 154 | 15 | 49 | 27 | 47 | 74 |
| Outer flakes, bleached, 24-hr hydrolysis THAA | 6 | 101 | 10 | 11 | 0 | 47 | 4 | 152 | 10 | 83 | 27 | 49 | 75 |

*Table S.1. Late Cretaceous titanosaur eggshell A amino acid concentrations in picomoles / mg. *Data from elution time > 58 min is of low accuracy due to elevated baseline values.*

| Treatment | Asx D/L | Glx D/L | Ser D/L | Ala D/L | Tyr D/L | Val D/L* | Phe D/L* | Leu D/L* | Ile D/L* | [Ser]/[Ala] |
|---|---|---|---|---|---|---|---|---|---|---|
| Bleached FAA | 0.00 | 1.03 | NA | 0.93 | 0.43 | 1.22 | 0.00 | NA | 1.81 | 0 |
| 18-hr hydrolysis THAA | 0.00 | 0.84 | 0.00 | 0.64 | NA | 0.87 | NA | NA | NA | 0.15 |
| Bleached, 24-hr hydrolysis THAA | NA | 1.04 | NA | 0.96 | 0.22 | 1.29 | NA | NA | 2.39 | 0 |
| Ethanol rinsed before powdering, bleached FAA | NA | 1.05 | NA | 0.97 | 0.00 | 1.11 | NA | NA | 0.00 | 0 |
| Ethanol rinsed before powdering, 18-hr hydrolysis THAA | NA | 0.98 | NA | 0.89 | NA | 1.12 | NA | NA | NA | 0 |
| Ethanol rinsed before powdering, 18-hr hydrolysis THAA | NA | 0.96 | NA | 0.89 | NA | 1.11 | NA | NA | NA | 0 |
| Ethanol rinsed before powdering, bleached, 24-hr hydrolysis THAA | NA | 1.02 | NA | 0.93 | 0.00 | 1.14 | NA | NA | 0.00 | 0 |
| Outer flakes, bleached FAA | 0.45 | 1.01 | 0.00 | 0.94 | 0.85 | 1.18 | 0.46 | 0.00 | 9.59 | 0.02 |
| Outer flakes, bleached FAA | 0.56 | 1.00 | 0.10 | 0.94 | 0.24 | 1.21 | 0.22 | 0.00 | 8.16 | 0.02 |
| Outer flakes, bleached FAA | 0.62 | 0.97 | 0.00 | 0.96 | 0.00 | 1.39 | 0.31 | 0.00 | NA | 0.02 |
| Outer flakes, bleached FAA | 0.67 | 0.97 | 0.00 | 0.95 | 0.00 | 1.09 | 0.23 | 0.00 | 6.05 | 0.02 |
| Outer flakes, bleached, 24-hr hydrolysis THAA | 0.22 | 0.96 | 0.08 | 0.86 | 0.60 | 1.21 | 0.35 | 0.00 | 8.06 | 0.07 |
| Outer flakes, bleached, 24-hr hydrolysis THAA | 0.29 | 0.96 | 0.11 | 0.86 | 0.00 | 1.18 | 0.18 | 0.00 | 7.08 | 0.07 |
| Outer flakes, bleached, 24-hr hydrolysis THAA | 0.34 | 0.95 | 0.05 | 0.88 | 0.62 | 2.34 | 0.46 | 0.00 | 10.04 | 0.07 |
| Outer flakes, bleached, 24-hr hydrolysis THAA | 0.31 | 0.95 | 0.00 | 0.87 | 0.00 | 1.68 | 0.46 | 0.00 | 9.06 | 0.06 |

*Table S.2. Late Cretaceous titanosaur eggshell A D/L and [Ser}/[Ala] values. *Data from elution time > 58 min is of low accuracy due to elevated baseline values. NA indicates that amino acid concentration was below detection limit or division by 0.*

Data File C:\CHEM32\1\DATA\G840\G840 2017-04-04 14-05-32\003-0701.D
Sample Name: 11489bH*

=====================================================================

Acq. Operator : Sheila Taylor    Seq. Line : 7
Acq. Instrument : Instrument 1    Location : Vial 3
Injection Date : 04/04/2017 22:12:02    Inj : 1
Inj Volume : Inj prog
Acq. Method : C:\CHEM32\1\DATA\G840\G840 2017-04-04 14-05-32\AAR_YK8.M
Last changed : 22/05/2015 11:37:32 by Kirsty Penkman
Analysis Method : C:\CHEM32\1\METHODS\AAR_YK8.M
Last changed : 11/07/2017 14:08:59 by Kirsty Penkman
(modified after loading)
Method Info : 2ul inject; 2.2ul OPA; 2.0 ul mixing; 13 cycles; 5% at 0, 23+0.4% @ 31 min; 48.82+5% @ 95 min, 95 min cutoff; 15 min flush; gain 11; flow 0.56ml/min; 3mm id

Sample Info : 11489bH* - 12.1bH* - Saltasaurus eggshell, Argentina, L ate Cretaceous, bleached 2 days, Hyd 110C 24 hrs, rehyd 20ul only

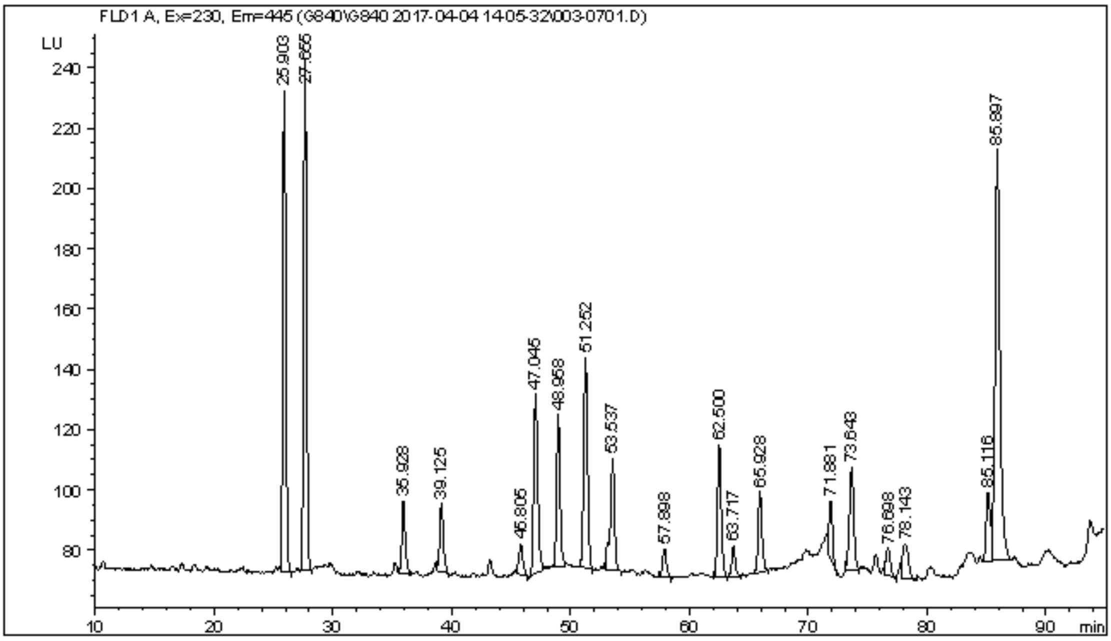


=====================================================================

Area Percent Report

=====================================================================

Sorted By : Signal
Multiplier: : 1.0000
Dilution: : 1.0000
Use Multiplier & Dilution Factor with ISTDs

Signal 1: FLD1 A, Ex=230, Em=445

| Peak # | RetTime [min] | Type | Width [min] | Area LU *s | Height [LU ] | Area % |
|---|---|---|---|---|---|---|
| 1 | 25.903 | BB | 0.2704 | 2750.70459 | 159.42297 | 13.6545 |
| 2 | 27.655 | BB | 0.2660 | 2858.24683 | 169.41339 | 14.1883 |

Instrument 1 11/07/2017 14:09:06 Kirsty Penkman    Page 1 of 2

*Figure S.1. RP-HPLC printout of the bleached, 24-hr hydrolysis Late Cretaceous titanosaur eggshell A THAA.*

```
Data File C:\CHEM32\1\DATA\G840\G840 2017-04-04 14-05-32\001-0501.D
Sample Name: 11489bF

    =====================================================================
    Acq. Operator   : Sheila Taylor                    Seq. Line :   5
    Acq. Instrument : Instrument 1                      Location : Vial 1
    Injection Date  : 04/04/2017 18:09:20                    Inj :   1
                                                     Inj Volume : Inj prog
    Acq. Method     : C:\CHEM32\1\DATA\G840\G840 2017-04-04 14-05-32\AAR_YK8.M
    Last changed    : 22/05/2015 11:37:32 by Kirsty Penkman
    Analysis Method : C:\CHEM32\1\METHODS\AAR_YK8.M
    Last changed    : 11/07/2017 14:08:30 by Kirsty Penkman
                      (modified after loading)
    Method Info     : 2ul inject; 2.2ul OPA; 2.0 ul mixing; 13 cycles; 5% at 0, 23+0.4% @
                      31 min; 48.82+5% @ 95 min, 95 min cutoff; 15 min flush; gain 11;
                      flow 0.56ml/min; 3mm id

    Sample Info     : 11489bF - 12.1bF - Saltasaurus eggshell, Argentina, Lat
                      e Cretaceous, bleached 2 days, Free, rehyd 20ul only
```

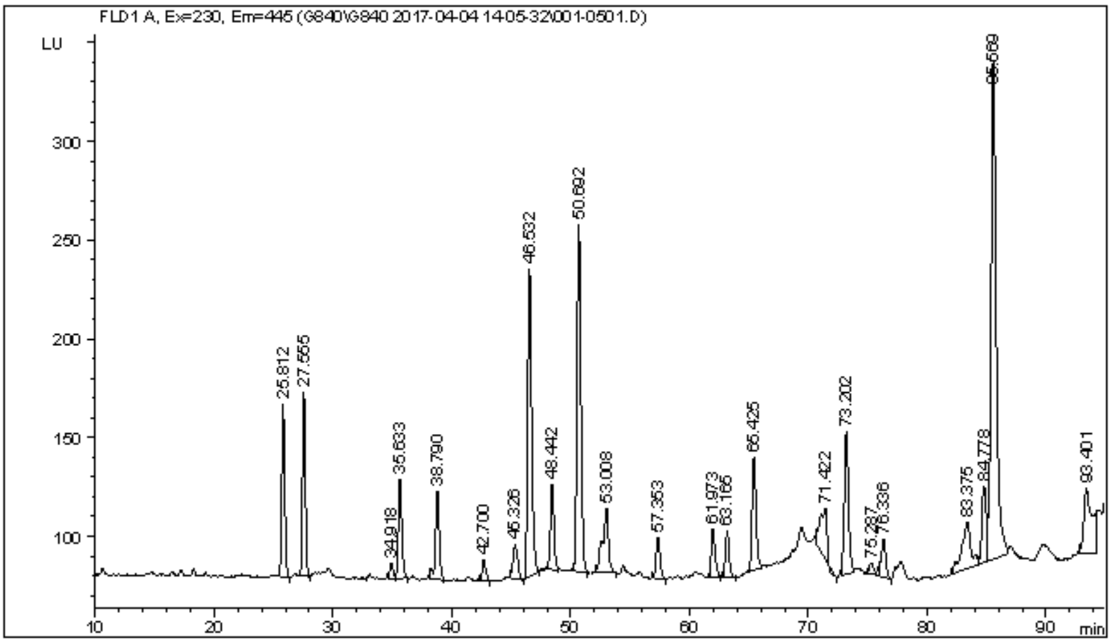


```
    =====================================================================
                           Area Percent Report
    =====================================================================

    Sorted By             :      Signal
    Multiplier:                   :      1.0000
    Dilution:                     :      1.0000
    Use Multiplier & Dilution Factor with ISTDs


    Signal 1: FLD1 A, Ex=230, Em=445

    Peak RetTime Type  Width     Area      Height     Area
      #   [min]        [min]  LU     *s   [LU    ]      %
    ----|-------|----|-------|----------|----------|--------|
       1  25.812 BB    0.2668 1488.37537   87.85777   4.4570
       2  27.555 BB    0.2524 1535.31726   93.32893   4.5975
       3  34.918 BV    0.2724  138.08318    7.60037   0.4135

Instrument 1 11/07/2017 14:08:45 Kirsty Penkman                        Page 1 of 2
```
*Figure S.2. RP-HPLC printout of the bleached Late Cretaceous titanosaur eggshell A FAA.*

| Treatment | [Asx] | [Glx] | [Ser] | [L-Thr] | [L-His] | [Gly] | [L-Arg] | [Ala] | [Tyr] | [Val]* | [Phe]* | [Leu]* | [Ile]* |
|---|---|---|---|---|---|---|---|---|---|---|---|---|---|
| Bleached FAA | 24 | 203 | 14 | 123 | 6 | 217 | 25 | 581 | 62 | 313 | 89 | 133 | 123 |
| Bleached FAA | 20 | 204 | 6 | 121 | 0 | 196 | 0 | 577 | 40 | 323 | 77 | 77 | 159 |
| Bleached FAA | 19 | 187 | 3 | 102 | 0 | 208 | 16 | 544 | 45 | 329 | 50 | 139 | 174 |
| Bleached FAA | 19 | 183 | 4 | 90 | 0 | 185 | 17 | 537 | 65 | 330 | 59 | 149 | 178 |
| Bleached, 24-hr hydrolysis THAA | 37 | 719 | 31 | 57 | 31 | 154 | 14 | 305 | 12 | 163 | 60 | 44 | 60 |
| Bleached, 24-hr hydrolysis THAA | 95 | 1835 | 70 | 121 | 34 | 406 | 32 | 804 | 53 | 491 | 279 | 167 | 180 |
| Bleached, 24-hr hydrolysis THAA | 92 | 1746 | 69 | 87 | 33 | 388 | 30 | 766 | 52 | 424 | 123 | 160 | 175 |

*Table S.3. Late Cretaceous titanosaur eggshell B amino acid concentrations in picomoles / mg. *Data from elution time > 58 min is of low accuracy due to elevated baseline values.*

| Treatment | Asx D/L | Glx D/L | Ser D/L | Ala D/L | Tyr D/L | Val D/L* | Phe D/L* | Leu D/L* | Ile D/L* | [Ser]/[Ala] |
|---|---|---|---|---|---|---|---|---|---|---|
| Bleached FAA | 0.90 | 0.99 | 0.32 | 0.92 | 0.44 | 1.09 | 0.33 | 0.00 | 4.34 | 0.02 |
| Bleached FAA | 0.98 | 1.04 | 1.75 | 0.92 | 0.00 | 1.19 | 0.22 | 0.00 | 5.83 | 0.01 |
| Bleached FAA | 1.03 | 1.02 | 0.79 | 0.94 | 0.00 | 1.34 | 0.44 | 0.00 | 6.17 | 0.00 |
| Bleached FAA | 1.01 | 1.01 | 0.96 | 0.94 | 0.31 | 1.40 | 0.46 | 0.00 | 5.46 | 0.01 |
| Bleached, 24-hr hydrolysis THAA | 0.23 | 0.99 | 0.10 | 0.67 | 0.83 | 1.47 | 0.56 | 0.00 | 7.99 | 0.10 |
| Bleached, 24-hr hydrolysis THAA | 0.23 | 0.99 | 0.00 | 0.70 | 0.30 | 1.16 | 1.59 | 0.00 | 6.74 | 0.09 |
| Bleached, 24-hr hydrolysis THAA | 0.23 | 0.99 | 0.03 | 0.70 | 0.31 | 0.93 | 0.19 | 0.00 | 6.33 | 0.09 |

*Table S.4. Late Cretaceous titanosaur eggshell B D/L and [Ser}/[Ala] values. *Data from elution time > 58 min is of low accuracy due to elevated baseline values.*

**LC-MS/MS:** The titanosaur eggshell A was first powdered in a mortar and pestle that had been sterilised with 70 % ethanol at the University of Bristol. LC-MS/MS analyses of this powder were then performed independently at the University of Turin (Italy) and University of Copenhagen (Denmark). The powdered sample was bleached in a comparable manner to the method described in the RP-HPLC section above (72 hr, 12 % NaOCl, 50 uL/mg). The bleached powders were then split and further preparation steps carried out independently at Turin and Copenhagen, following the protocol described in Demarchi *et al.* (2016). Briefly, powders were demineralised in 0.6 M HCl and the peptides concentrated directly using zip-tips (Turin) and stage-tips (Copenhagen), without performing a digestion step. Both laboratories applied the controls detailed in Hendy *et al.* (2018); however, the Copenhagen lab is equipped for ancient DNA analysis and therefore operates in a sterile environment, while the preparation in Turin was carried out under a clean laminar flow cabinet.

*Turin analytical procedure:* Dried peptides were resuspended in 0.1% formic acid and analysed using an Ultimate 3000 Dionex nanoHPLC instrument coupled with an Orbitrap Fusion (Thermo Scientific, Milan, Italy). Separation was achieved using a PepMap RSLC C18, 2 µm, 100 Å, 75 µm × 50 cm column (Thermo Scientific) and an Acclaim PepMap100 C18, 5 µm, 100 Å, 300 µm i.d. × 5 mm, preconcentration column (Thermo Scientific). The eluent used for preconcentration step was 0.05% trifluoroacetic acid in water/acetonitrile 98/2 and the flowrate was 5 µL/min. The eluents used for chromatographic separation were 0.1% formic acid in water (solvent A) and 0.1% formic acid in acetonitrile/water 8/2 (solvent B) in a program which was initially isocratic at 5:95 (A:B %) for 5 min, increased to 75:25 in 55 min, run up to 60:40 in 6 min, and to 10:90 in 5 min. Recondition time was 20 min. The injection volume was 1 µL and the flow rate 300 nL $min^{-1}$. The nanocolumn was provided with the ESI source. The mass spectrometry parameters were as follows: positive spray voltage 2,300 (V), sweep gas 1 (Arb), and ion transfer tube temperature 275°C. Full scan spectra were acquired in the range of m/z 375–1,500 (resolution 120,000 at m/z 200). MSn spectra in data-dependent analysis mode were acquired in the range between the ion trap cut-off and precursor ion m/z values. HCD collision energy was fixed at 28%, orbitrap resolution 50,000 and the isolation window was 1.6 m/z units. Blank samples were analysed before and after the titanosaur extract.

*Copenhagen method:* Prior to LC-MS/MS, the stage-tipped sample was eluted in 30 µL 40% acetonitrile (ACN) 0.1% formic acid (FA) and vacuum centrifuged until < 3 µL remained. 5 µL of 0.1% trifluoroacetic acid (TFA), 5% ACN was then added. The sample was separated on a 15 cm column (75 µm inner diameter) in-house laser pulled and packed with 1.9 µm C18 beads (Dr. Maisch, Germany) on an EASY-nLC 1200 (Thermo Fisher Scientific, Bremen, Germany) connected to a Q-Exactive HF-X (Thermo Fisher Scientific, Bremen, Germany) on a 77 min gradient. The column temperature was maintained at 40°C using an integrated column oven. Buffer A was milliQ water. 5 µL of sample was injected. The peptides were separated with increasing buffer B (80% ACN and 0.1% FA), going from 5% to 30% in 50 min, 30% to 45% in 10 min, 45% to 80% in 2 min, held at 80% for 5 min before dropping back down to 5% in 5 min and held for 5 min. Flow rate was 250 nL/min. A wash-blank method using 0.1% TFA, 5% ACN was run in between each sample to hinder cross contamination. The Q-Exactive HF-X was operated in data dependent top 10 mode. Spray voltage was 2 kV, S-lens RF level at 50, and heated capillary at 275°C. Full scan mass spectra were recorded at a resolution of 120,000 at m/z 200 over the m/z range 350–1,400 with a target value of 3e6 and a maximum injection time of 25 ms. HCD-generated product ions were recorded with a maximum ion injection time set to 108 ms and a target value set to 2e5 and recorded at a resolution of 60,000. Normalised collision energy was set at 28% and the isolation window was 1.2 m/z with the dynamic exclusion set to 20 s.

*Bioinformatics:* Product ion spectra were analysed using the software PEAKS v. 8.5 (Zhang *et al.* 2012). The results reported were obtained by SPIDER searches of all the *de novo* peptides against a database containing all known peptide sequences from Reptilia and Aves (total 3,865,676 sequences, downloaded on 09/07/2019) and common contaminants (cRAP, https://www.thegpm.org/crap/). Acceptance thresholds applied were as follows: False Discovery Rate ≤ 0.5 %, Peptide -10lgP ≥ 17.4, Protein -10lgP ≥ 20, Proteins unique peptides ≥ 2, *De novo* ALC Score ≥ 80 %. Data from both Turin and Copenhagen eggshell samples, as well as the Turin procedural and wash blank are available via ProteomeXchange Consortium (http://proteomecentral.proteomexchange.org) via the PRIDE partner repository (Perez-Riverol *et al.* 2019) with data set identifier PXD014712 (Username:; Password: 9BP6ydxK).

**University of Turin**

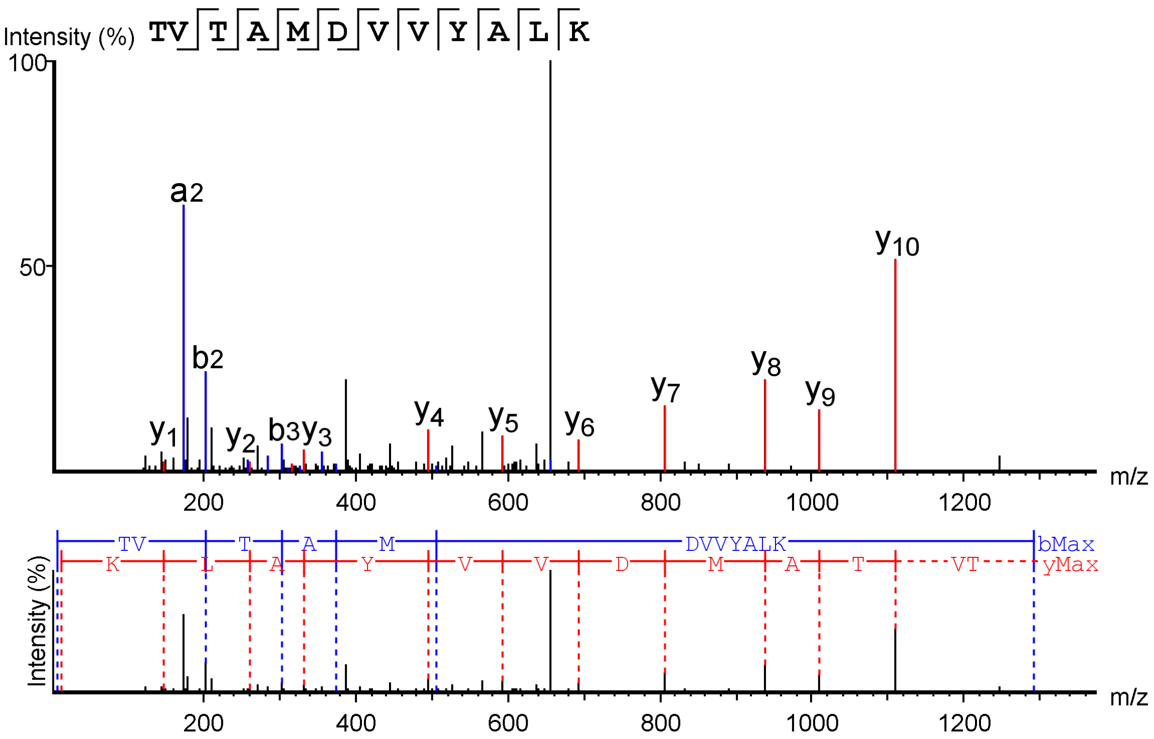


*Figure S.3. Product ion spectrum of peptide TVTAMDVVYALK from the sequence of Histone H4 [*Gallus gallus*] identified in Late Cretaceous titanosaur eggshell A.*

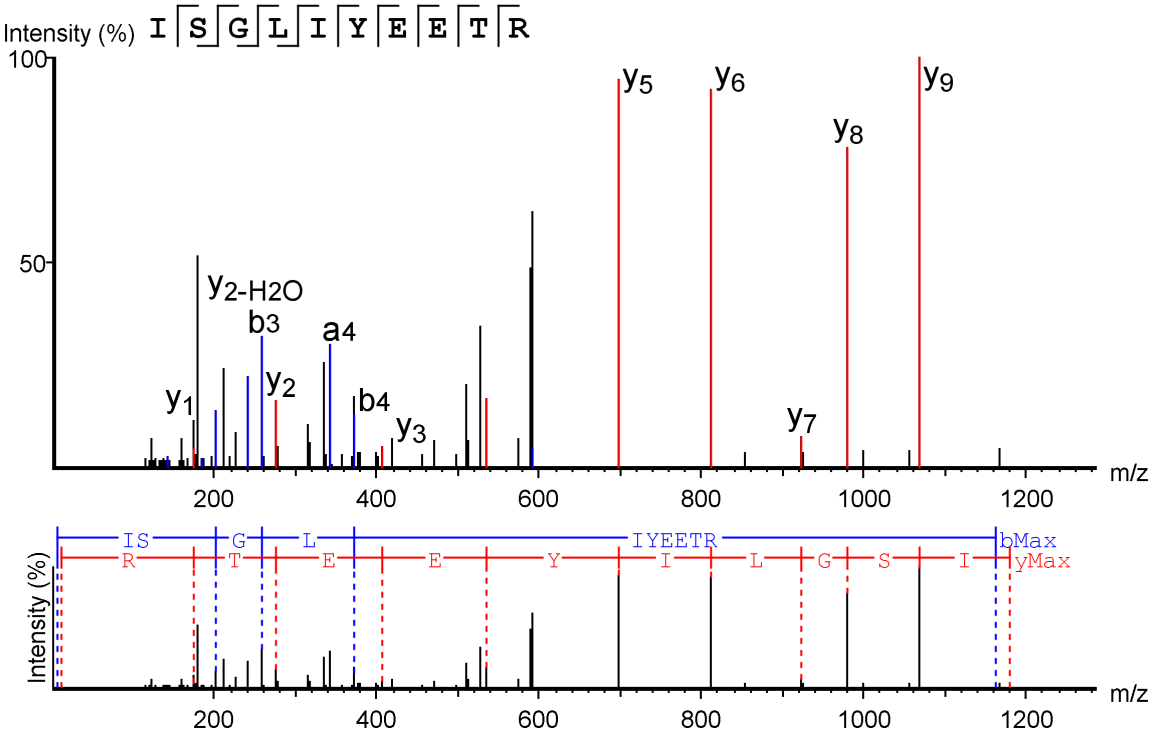


*Figure S.4. Product ion spectrum of peptide ISGLIYEETR from the sequence of Histone H4* [Gallus gallus] *identified in Late Cretaceous titanosaur eggshell A.*

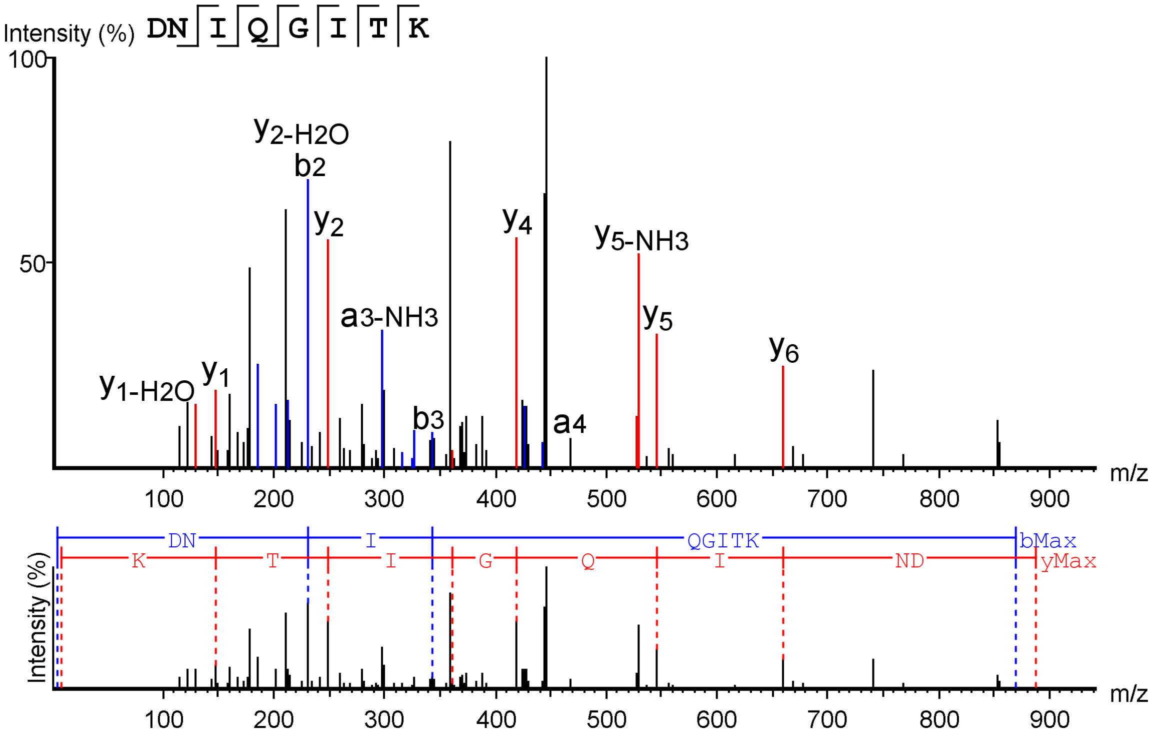


*Figure S.5. Product ion spectrum of peptide DNIQGITK from the sequence of Histone H4* [Gallus gallus] *identified in Late Cretaceous titanosaur eggshell A.*

1 MSGRGKGGKG LGKGGAKRHR KVLR**DNIQGI**

31 **TK**PAIRRLAR RGGVKR**ISGL IYEETR**GVLK

61 VFLENVIRDA VTYTEHAKRK **TVTAMDVVYA**

91 **LK**RQGRTLYG FGG

*Figure S.6. Coverage of protein histone H4* [Gallus gallus] *identified in bleached titanosaur eggshell A (Turin preparation). Note that one of the three peptides supporting the identification of the protein, DNIQGITK, contains two potential deamidation sites, which are unmodified.*

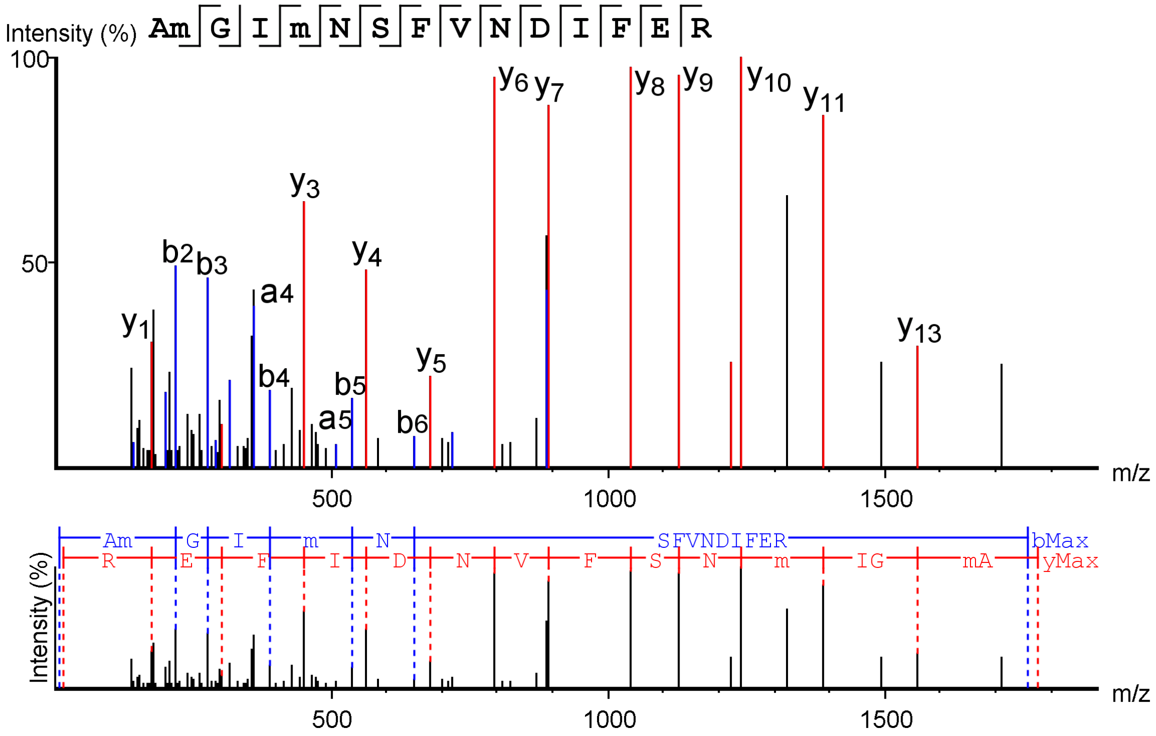


*Figure S.7. Product ion spectrum of peptide AM(+15.99)GIM(+15.99)NSFVNDIFER, identified in Late Cretaceous titanosaur eggshell A. The peptide sequence was not matched to any known protein by the software PEAKS, but a BLASTp search on UniprotKB_SwissProt yielded 100% sequence identity to a region of Isoform 2 of Histone H2B type 2-F [*Homo sapiens*].*

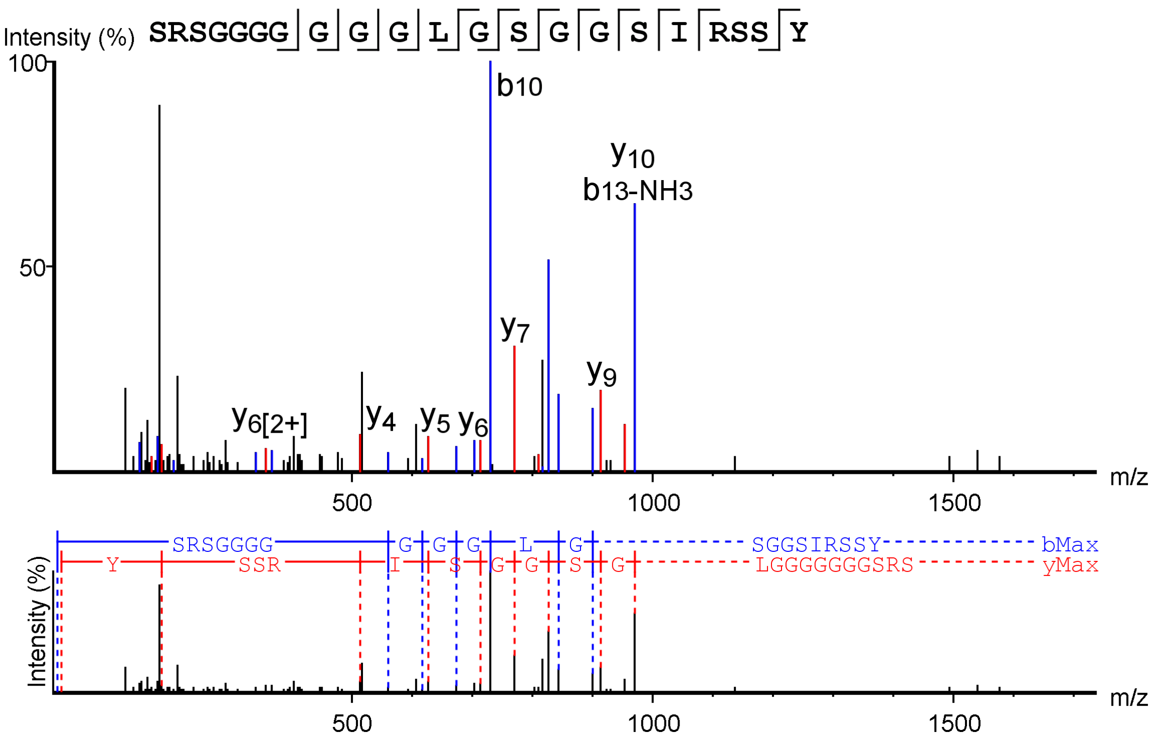


*Figure S.8. Product ion spectrum of peptide SRSGGGGGGGLGSGGSIRSSY, identified in Late Cretaceous titanosaur eggshell A. The peptide sequence was not matched to any known protein by the software program PEAKS, but a BLASTp search on UniprotKB_SwissProt yielded 100% sequence identity with Keratin, type I cytoskeletal 9 [*Homo sapiens*].*

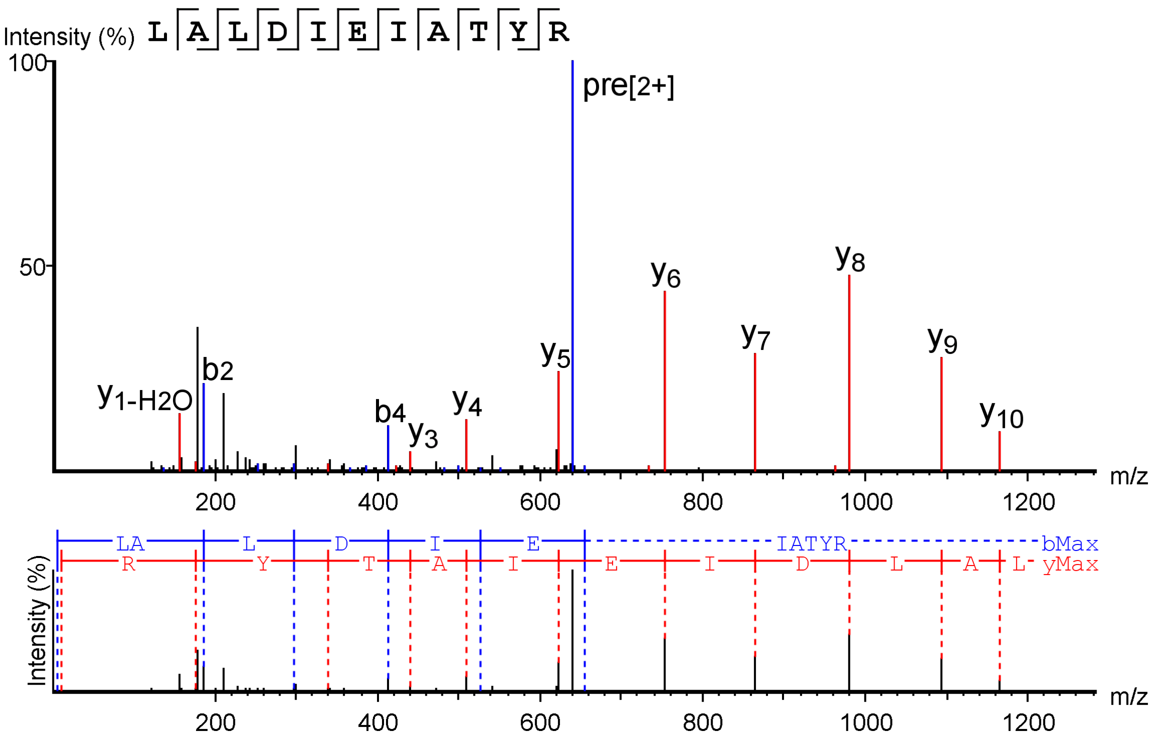


*Figure S.9. Product ion spectrum of peptide LALDIEIATYR, identified in Late Cretaceous titanosaur eggshell A. The peptide sequence was not matched to any known protein by the software program PEAKS, but a BLASTp search on UniprotKB_SwissProt yielded 100% sequence identity with Keratin, type II cytoskeletal 4 [*Homo sapiens*].*

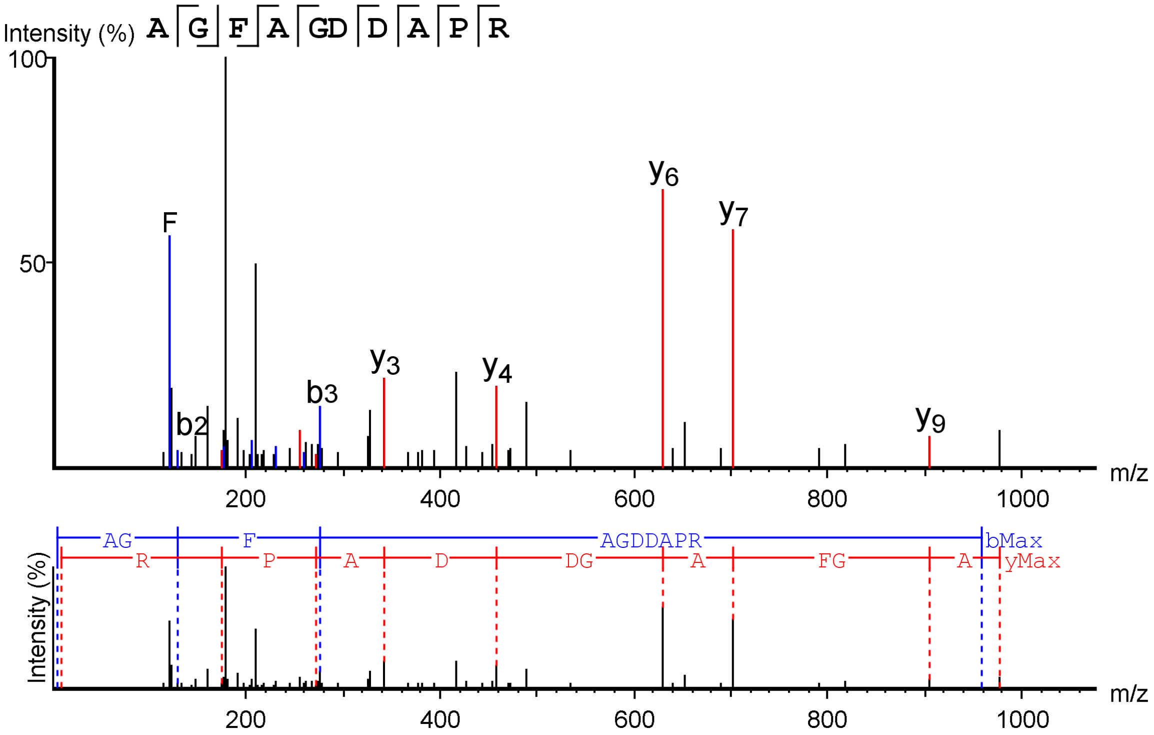


*Figure S.10. Product ion spectrum of peptide AGFAGDDAPR, identified in Late Cretaceous titanosaur eggshell A (prepared at the University of Turin). The peptide sequence was not matched to any known protein by the software program PEAKS, but a BLASTp search on UniprotKB_SwissProt yielded 100% sequence identity with POTE ankyrin domain family member I [*Homo sapiens*].*

| **Peptide** | **ALC (%)** | **m/z** | **z** | **Mass** | **ppm** | **Local confidence (%)** | **BLAST search (100% identity)** |
|---|---|---|---|---|---|---|---|
| ESYSVYVYK | 93 | 569.2813 | 2 | 1136.539 | 7.9 | 79 81 95 98 98 98 98 98 96 | Histone H2B type 1-K (*Macaca fascicularis*) |
| LLLLLLL | 93 | 405.8103 | 2 | 809.599 | 8.8 | 92 95 94 93 93 95 93 | No hits |
| PEN(+.98)VLLGTFKVLD | 88 | 723.4051 | 2 | 1444.781 | 10 | 38 77 92 94 100 100 95 98 99 98 94 88 75 | No 100% identity |
| LLLLLLL | 87 | 405.8092 | 2 | 809.599 | 6 | 90 91 90 79 82 90 90 | No hits |
| LAAAARFMAW | 85 | 554.2917 | 2 | 1106.57 | -0.7 | 86 88 99 99 97 96 94 87 60 40 | Uncharacterized protein (*Microdochium bolleyi*) |
| YTKLLLLN | 80 | 489.308 | 2 | 976.5957 | 5.8 | 50 54 86 82 92 93 95 89 | No hits |

*Table S.5.* De novo *peptide sequences reconstructed by the software program PEAKS on the basis of tandem mass spectra of titanosaur eggshell A (prepared at the University of Turin).*

| **Sample** | **Protein name** | **Peptide** | **-10lgP** | **#Spectra** |
|---|---|---|---|---|
| Procedural blank | Albumin precursor [*Homo sapiens*] | DAHKSEVAHRFKDLGEENFKALVLIAFAQ(-.98) | 61.67 | 5 |
| | | DAHKSEVAHRFKDLGEENFKALVLIAFAQ | 46.35 | 5 |
| | | DAHKSEVAHRFKDLGEENFKALVLIAFAQYLQQ(-.98) | 33.39 | 1 |
| | | FAEEGKKLVAASQAALGL | 32.12 | 7 |
| | Serum albumin precursor [*Homo sapiens*] | LVAASQAALGL | 29.09 | 1 |
| | Tubulin (highly conserved in various taxa) | GHYTEGAELVDSVLDVVR | 39.96 | 1 |
| | Conserved sequence | PGVVPFKR | 25.88 | 2 |
| | Tubulin (highly conserved in various taxa) | EIIDLVLDR | 25.65 | 7 |
| | Histone H4 [*Chelonia mydas*] | DNLQGITK | 24.28 | 1 |
| Wash blank | Chain A, histone H2a | AGLQFPVGR | 29.28 | 1 |
| | | VTIAQGGVLPNIQAVLLPK | 48.39 | 1 |
| | Histone H2B [*Gallus gallus*] | AMGIMNSFVNDIFER | 40.91 | 2 |
| | | QVHPDTGISSK | 48.98 | 1 |

*Table S.6. Peptide sequences from various proteins identified in the procedural blank and the four peptides belonging to histone sequences identified in the wash blank (prepared at the University of Turin).*

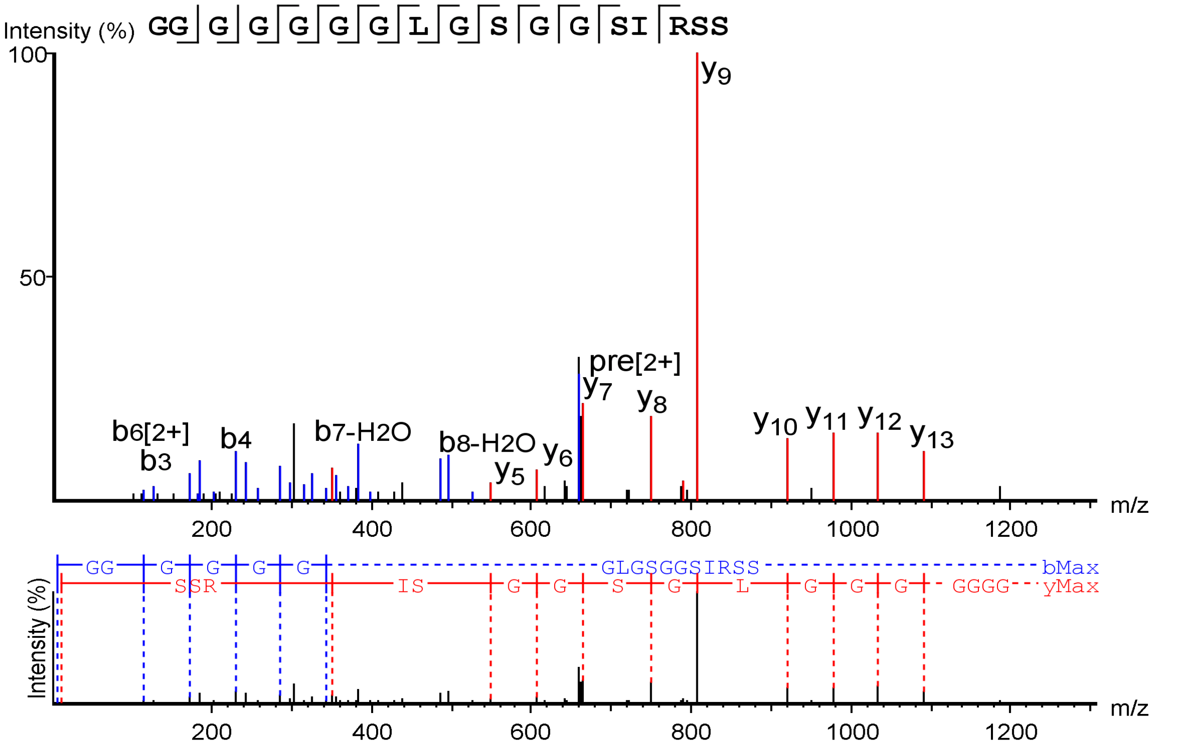


*Figure S.11. Product ion spectrum of peptide GGGGGGGLGSGGSIRSS, from the sequence of Keratin, type I cytoskeletal 9 [*Homo sapiens*], identified in Late Cretaceous titanosaur eggshell A.*

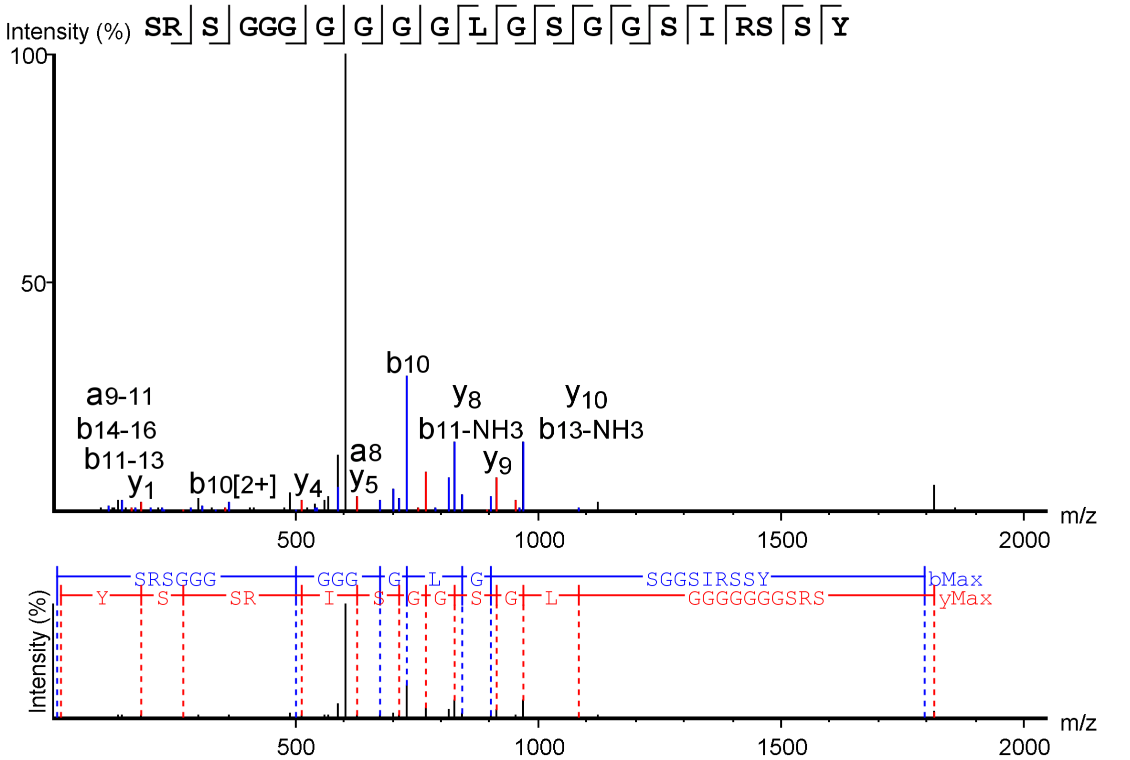


*Figure S.12. Product ion spectrum of peptide SRSGGGGGGGLGSGGSIRSSY, from the sequence of Keratin, type I cytoskeletal 9 [*Homo sapiens*], identified in Late Cretaceous titanosaur eggshell A.*

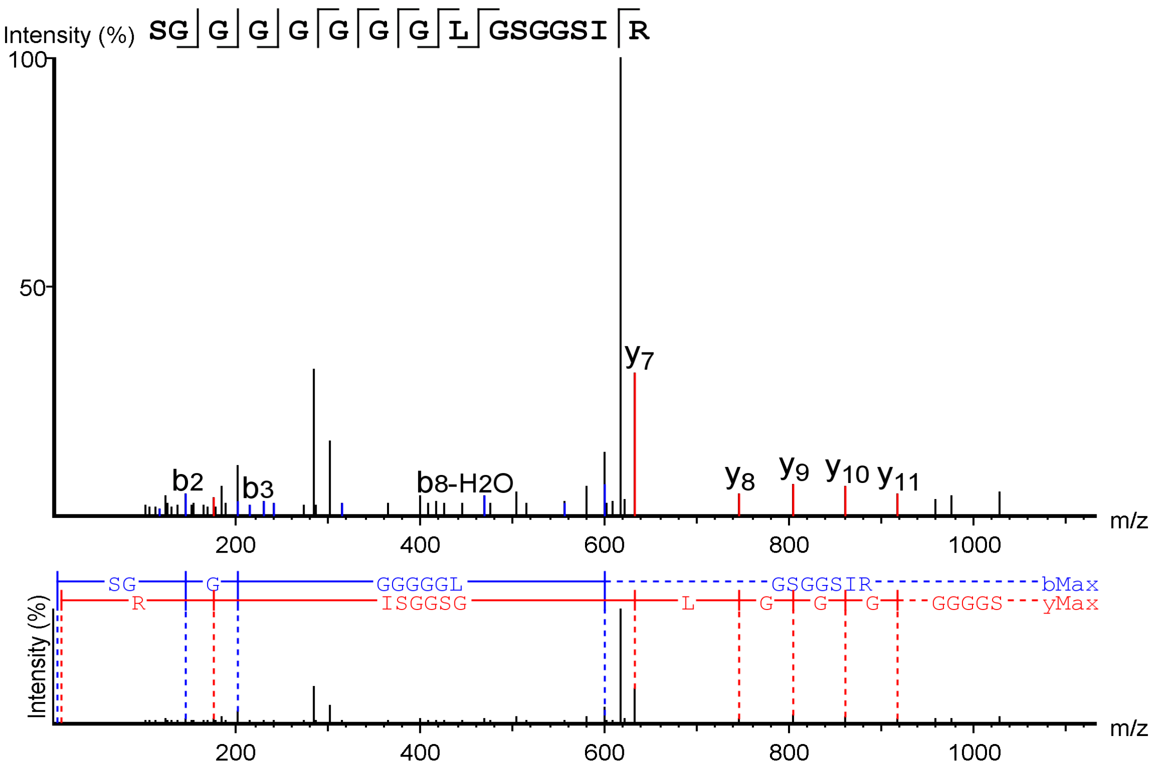


*Figure S.13. Product ion spectrum of peptide SRSGGGGGGGLGSGGSIRSSY, from the sequence of Keratin, type I cytoskeletal 9 [*Homo sapiens*], identified in Late Cretaceous titanosaur eggshell A.*

| Peptide | ALC (%) | *m/z* | z | Mass | ppm | Local confidence (%) | BLAST search (100% identity) |
|---|---|---|---|---|---|---|---|
| TN(+.98)LAHR | 87 | 356.6913 | 2 | 711.3664 | 2.3 | 79 82 94 90 90 89 | No hits |
| TVLVKKH | 86 | 412.7676 | 2 | 823.528 | -8.8 | 67 68 90 95 99 95 94 | No hits |
| ERQFSSRGS | 86 | 527.2607 | 2 | 1052.5 | 6.5 | 92 74 97 99 99 98 92 56 65 | No hits |

*Table S.7.* De novo *peptide sequences reconstructed by software PEAKS on the basis of tandem mass spectra for titanosaur eggshell A (prepared at the University of Copenhagen).*

**PY-GC-MS:** Py-GC-MS was run at the University of Bristol (School of Chemistry) and Newcastle University due to the desire for some degree of replicability, although analytical conditions slightly varied. .cdf and .raw files containing the raw chromatogram and spectrometry data are available.

Titanosaur eggshell A and the modern chicken eggshell were treated prior to analysis at the University of Bristol by rinsing the fragments with 70 % ethanol to reduce surficial contamination followed by powdering with a mortar and pestle that had been sterilised with 70 % ethanol. The interior membrane of the modern chicken eggshell was dissected from the mineralised eggshell prior to powdering using tweezers sterilised with 70 % ethanol so as not to be included in subsequent analyses.

*Bristol method:* A quartz tube was loaded with ~1 mg of the titanosaur and chicken sample powder and capped with glass wool. A pyrolysis unit (Chemical Data Systems [CDS] 5200 series pyroprobe) was coupled to a gas chromatograph (GC; Agilent 6890A; Varian CPSil-5CB fused column: 0.32 mm inner diameter, 0.45 µm film thickness, 50 m length, 100 % dimethylpolysiloxane) and a double focussing mass spectrometer (ThermoElectron MAT95, ThermoElectron, Bremen; electron ionisation mode: 310 °C GC interface, 200 °C source temperature) with a 2 mL $min^{-1}$ helium carrier gas. Samples were pyrolysed in the quartz tube (20 s, 610 °C), transferred to the GC (310 °C pyrolysis transfer line), and injected onto the GC (310 °C injector port temperature was maintained, 10:1 split ratio). The oven was programmed to heat from 50 °C (held for 4 min) to 300 °C (held for 15 min) by 4 °C $min^{-1}$. A *m/z* range of 50–650 was scanned (one scan per second). There was a 7 min delay whereby the filament was switched off for protection against any pressure increases at the start of the run. MAT95InstCtrl (v1.3.2) was used to collect data. QualBrowser (v1.3, ThermoFinnigan, Bremen) was used to view data.

*Newcastle method:* Powdered titanosaur eggshell A and modern chicken eggshell samples were also analysed at Newcastle and were treated prior to analysis using a DCM rinse followed by Soxhlet extraction to remove depositional ingress. One subsample of titanosaur eggshell A underwent the bleach treatment described in the RP-HPLC section above but did not receive DCM and Soxhlet extraction. Py-GC-MS analysis of all samples at Newcastle was performed on a CDS Pyroprobe 1000 via a CDS1500 valved interface (320 °C) to a Hewlett-Packard 6890GC split injector (320 °C) coupled to a Hewlett-Packard 5973MSD (emission current 35 uA, electron voltage 70 eV, quadrupole temperature 150 °C, source temperature 230 °C, multiplier voltage 2200 V, interface temperature 320 °C). A HP kayak XA chemstation computer (full scan mode: 50–650 amu) controlled acquisition. Approximately 1 mg of the sample was weighed into a quartz tube with glass wool end plugs. The tube was then placed into a pyroprobe platinum heating coil and sealed into the valved interface. The sample was pyrolysed at 610 °C for 10 s with the split open, while the GC temperature programme and data acquisition commenced. Separation was performed using a fused silica capillary column (0.25 mm inner diameter, 60 m length) coated with 0.25 µm of 5 % phenyl methyl silicone (HP- 5MS). Initially, the GC was held at 50 °C for 5 min. Then, temperature was programmed from 50–320 °C at 5 °C $min^{-1}$ and held at final temperature for 15 min, for a total of 65 min, with helium as the carrier gas (120 kPa initial pressure, 1 mL/min constant flow, split at 30 mL/min).

Compounds were identified with the aid of the National Institute of Standards and Technology (NIST98) database.

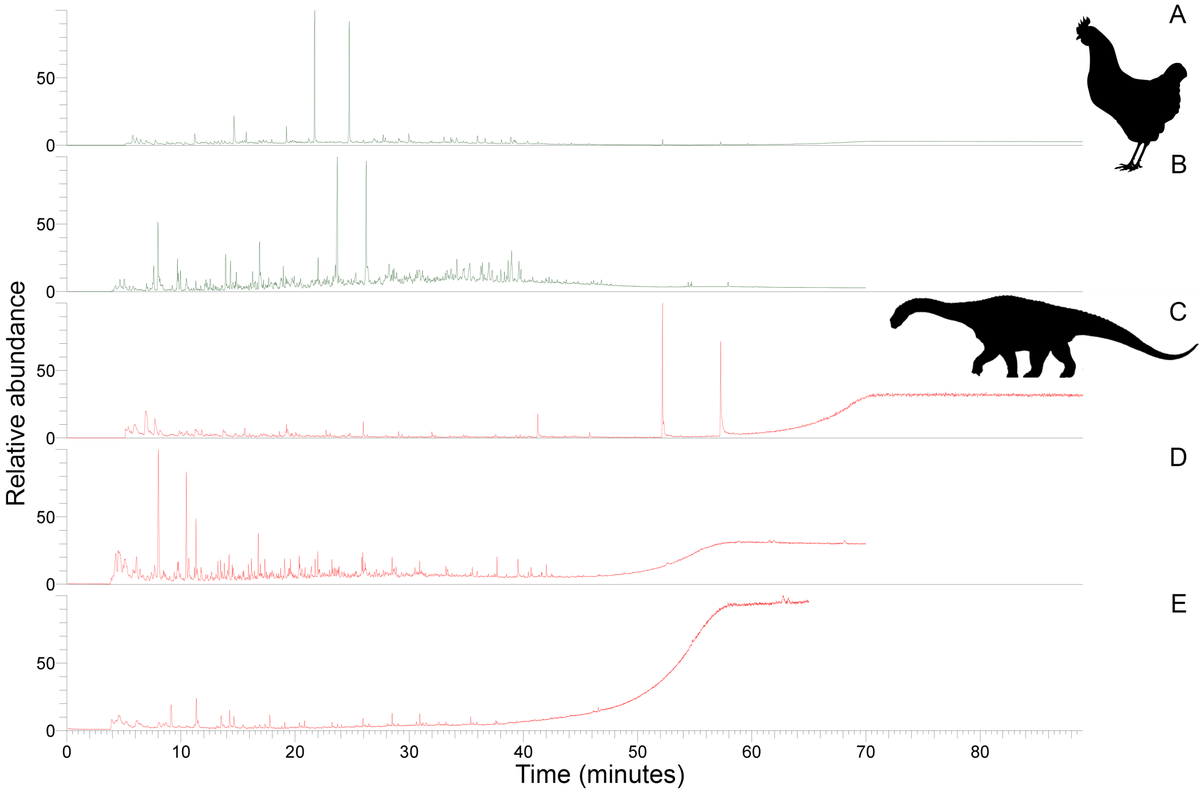


*Figure S.14. Total ion chromatograms from Py-GC-MS of, A–B, modern chicken and, C–E, titanosaur eggshell A. A, ethanol rinsed before powdering. B, ethanol rinsed before powdering, DCM rinsed, and Soxhlet extracted. C, ethanol rinsed before powdering. D, DCM rinsed and Soxhlet extracted. E, bleached. Note that A, C were obtained at Bristol while B, D–E were obtained at Newcastle with E having been run after B, D, making comparison more difficult by shifting retention times.*

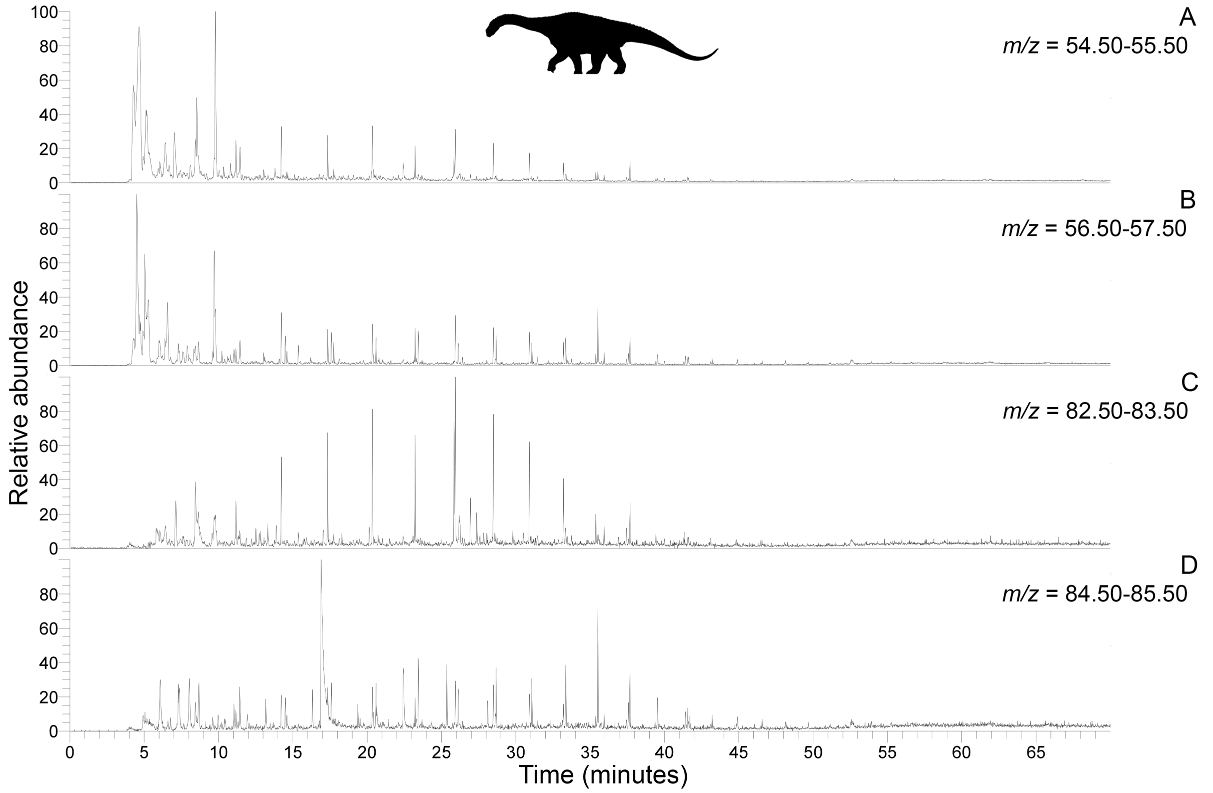


*Figure S.15. Py-GC-MS chromatograms searching for ion* m/z *ranges typical of alkanes and alkenes from kerogen in DCM rinsed and Soxhlet extracted titanosaur eggshell A. Doublets are strongly apparent. A,* m/z *= 55. B,* m/z *=57. C,* m/z *=83. D,* m/z *=85.*

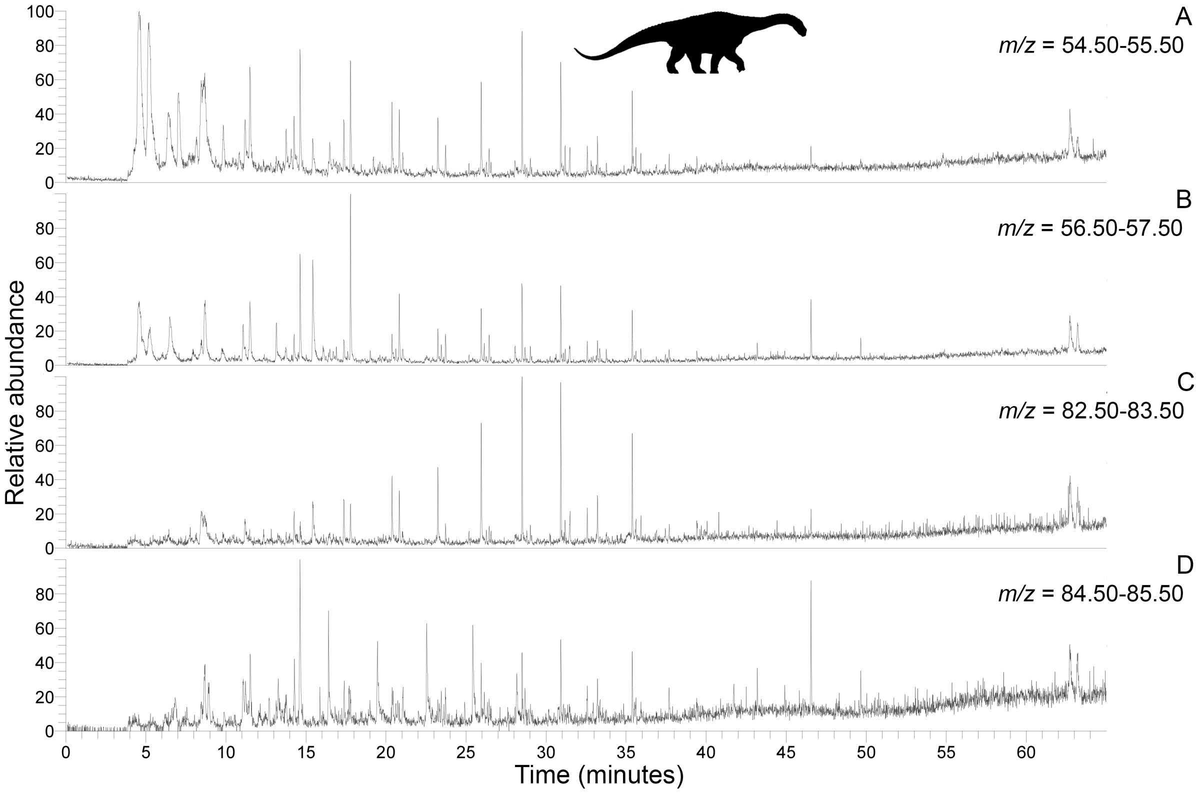


*Figure S.16. Py-GC-MS chromatograms searching for ion* m/z *ranges typical of alkanes and alkenes from kerogen in bleached titanosaur eggshell A (not DCM rinsed or Soxhlet extracted). Doublets are apparent. A,* m/z *= 55. B,* m/z *=57. C,* m/z *=83. D,* m/z *=85.*

**ASEPTIC POLISHING PROTOCOL:** Three falcon tubes were rinsed five times with milli-Q water then filled with 5 mL of ethanol, shaken, then emptied. Diamond grit powder was then added to the tubes such that each tube had a certain rating of grit: 600, 1200, or 3000. Then, 10 mL of 70 % ethanol was added to the tubes, the tubes were shaken and then centrifuged at 5000 g for 5 minutes. The supernatant was pipetted off to yield a paste. A glass polishing plate was rinsed with 10 % bleach, then wiped with a Kimwipe, then rinsed with 70 % ethanol. The 600 rated diamond paste was added to the plate using a disposable pipette and milli-Q water was added as necessary.

The eggshell fragment (titanosaur A and B, modern ostrich) was polished by hand while wearing gloves that had been rinsed with 70 % ethanol. Ethanol, followed by milli-Q water, was used to clear the grit off of the plate in between different polishing stages from 600, to 1200, to 3000, as well as between each sample (titanosaur eggshell followed by ostrich eggshell). A bleach (10 %) rinse and wipe with a Kimwipe prior to the ethanol and milli-Q water was used to clean the glass plate prior to the ostrich eggshell fragment to avoid contamination from the fossil fragments polished before it.

After polishing, each of the three eggshell fragments was stored in separate glass vials that had been rinsed with 70 % ethanol.

**TOF-SIMS:** TOF-SIMS was performed at Newcastle University and at Northwestern University.

##### Newcastle University

The resin-embedded thin sectioned samples were given three 8-minute ultrasonic washes in fresh iso-hexane in an attempt to remove any polishing residues. The samples for analysis were mounted directly onto a sample holder using clean stainless steel screws and clips.

Static SIMS analyses were carried out using an ION-TOF 'TOF-SIMS IV – 200' instrument (ION-TOF GmbH, Münster, Germany) of single-stage reflectron design (Schwieters *et al.* 1991). Positive and negative ion spectra and images were obtained using a $Bi_3^+$ focused liquid metal ion gun at 25 keV energy, incident at 45° to the surface normal and operated in 'bunched' mode for high mass resolution. This mode used ~20 ns wide ion pulses at a 10 kHz repetition rate. Charge compensation was affected by low-energy (~20 eV) electrons provided by a flood gun. The total ion dose density was $5 \times 10^{15}$ ions $m^{-2}$. The topography of the sample surface and the ion gun mode of operation limited the mass resolution in this work to approximately $m/\Delta m = 4000$. The spatial resolution was limited by the primary ion beam diameter to ~4 µm.

Positive and negative ion static SIMS spectra and images were recorded from the outermost ~1 nm of the sample surface at room temperature. Raw data containing the secondary ions recorded at each pixel was acquired with a 128 × 128 pixel raster and a field of view of 500 µm × 500 µm.

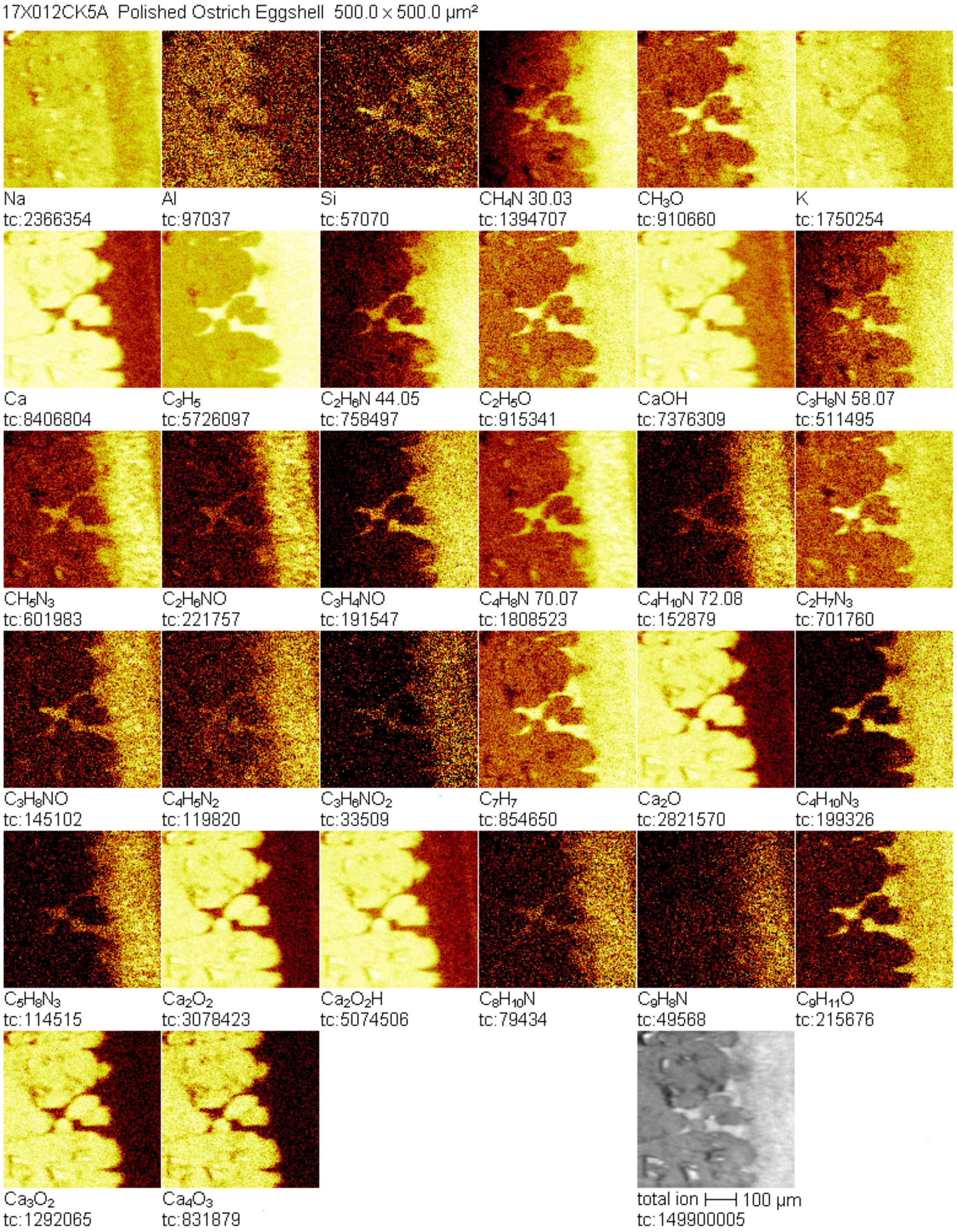


*Figure S.17. Positive TOF-SIMS mapping performed at Newcastle University on a resin-embedded ostrich eggshell thin section showing dominance of the organic signal arising from the epoxy resin even in this modern, presumably organic-rich sample.*

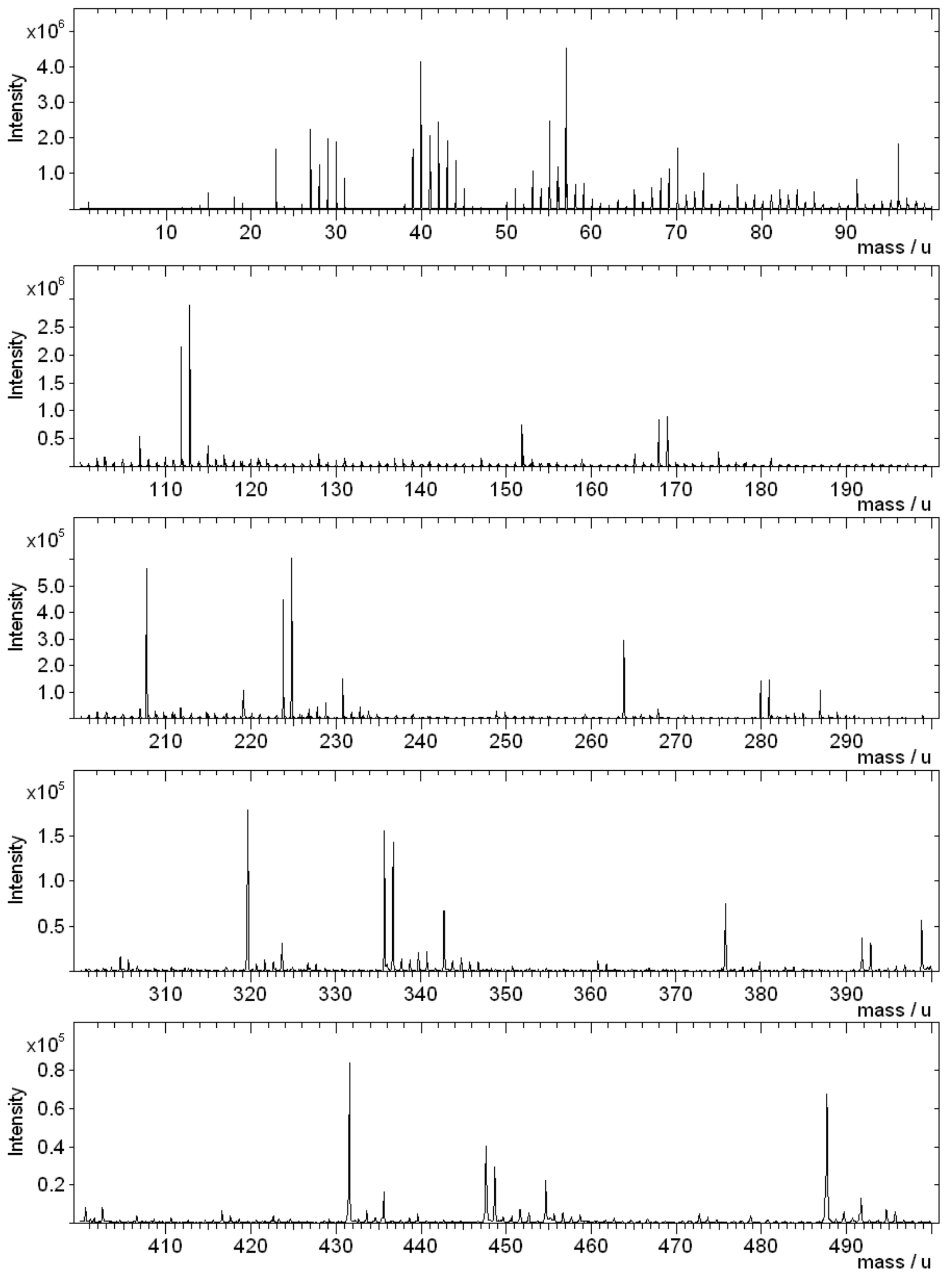


*Figure S.18. Positive TOF-SIMS spectrum performed at Newcastle University of resin-embedded ostrich eggshell thin section.*

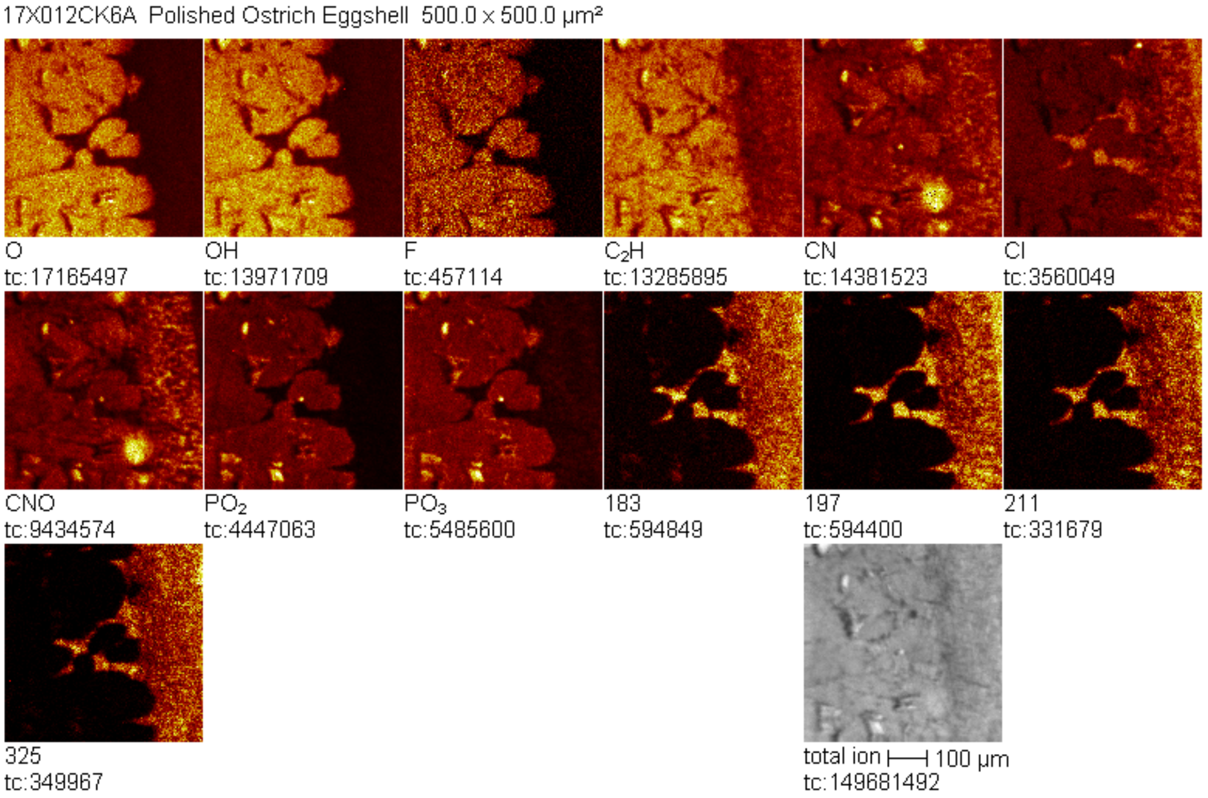


*Figure S.19. Negative TOF-SIMS mapping performed at Newcastle University on a resin-embedded ostrich eggshell thin section.*

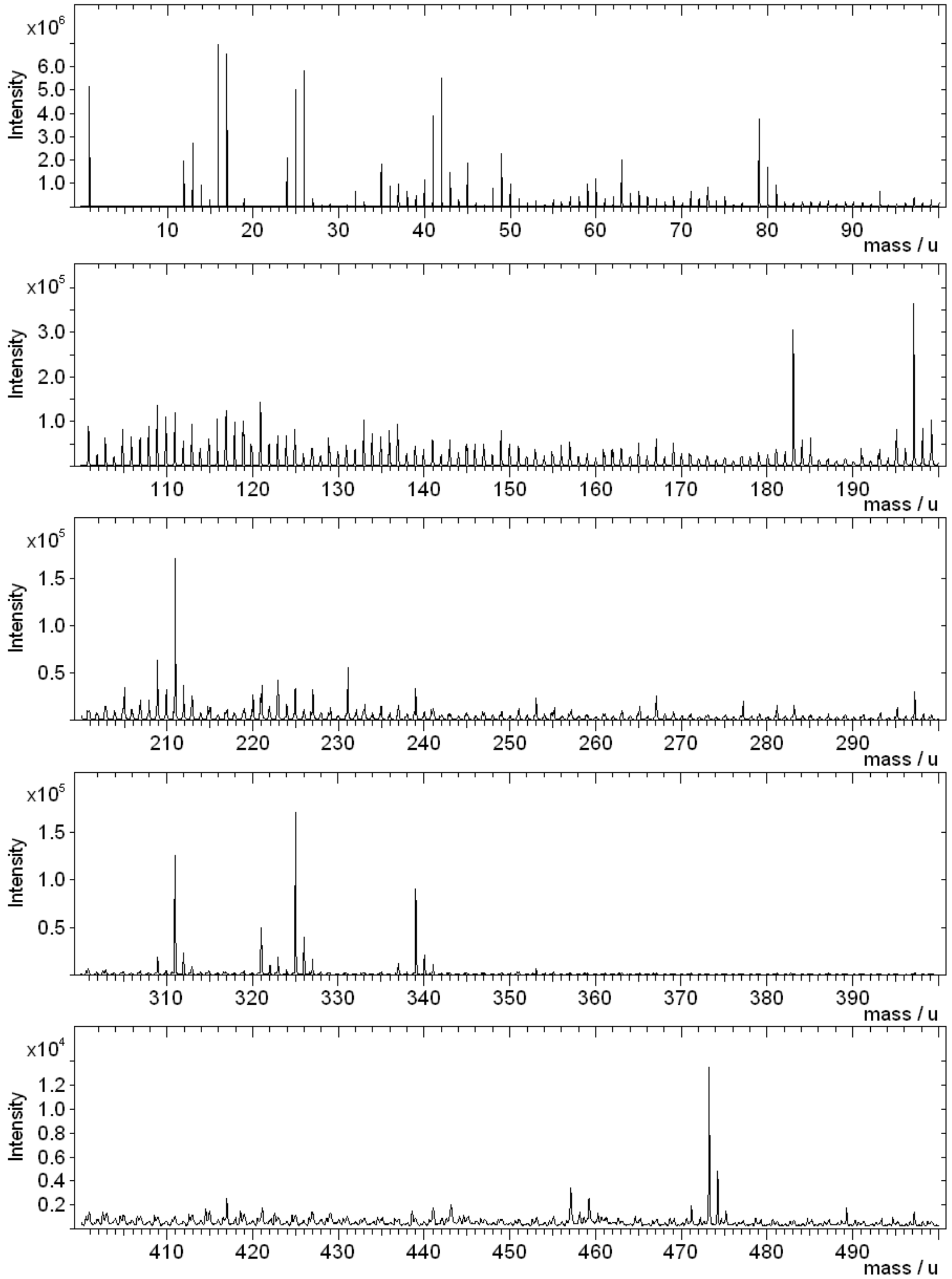


*Figure S.20. Negative TOF-SIMS spectrum performed at Newcastle University of resin-embedded ostrich eggshell thin section.*

#### Northwestern University

Aseptically polished (non-embedded) samples were examined at Northwestern University under TOF-SIMS carried out on a Physical Electronics TRIFT III spectrometer. The polished samples were mounted under a metal mesh to reduce the surface charging. The primary ion source was a gallium beam with 25KeV energy for ion mapping.

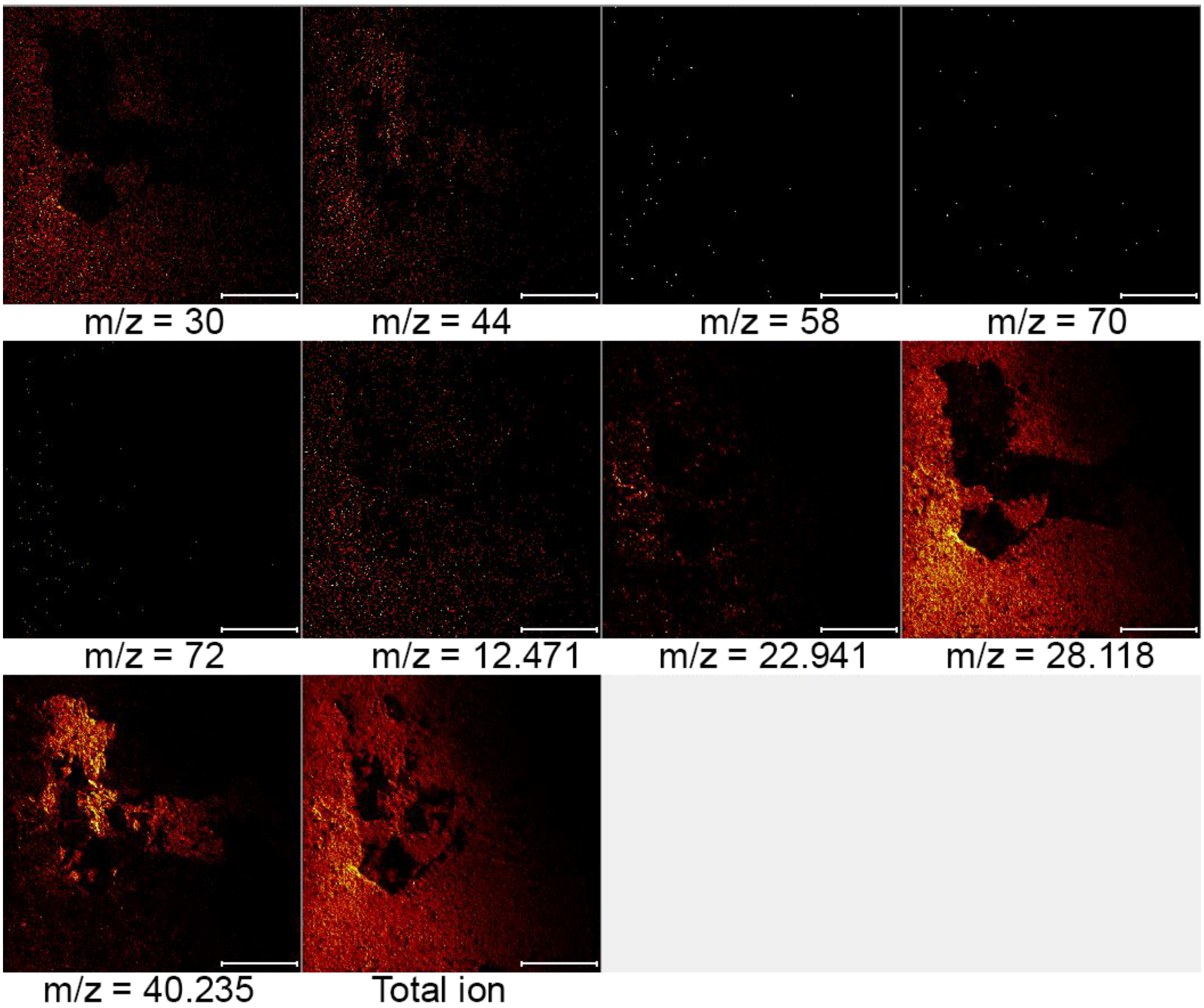


*Figure S.21. Positive TOF-SIMS mapping performed at Northwestern University on a cross section of aseptically polished fragment of titanosaur eggshell A highlighting secondary ions of potential amino acid fragments from Hedberg* et al. *2012 (*m/z = *~30 [Gly], ~44 [Ala], ~58 [Glu], ~70 [Val], ~72 [Val]), C (*m/z = *~12), Na (*m/z = *~23), Ca (*m/z = *~40), and the total secondary ion map. The signal at* m/z = *~28 could be consistent with either $CH_2N$, $C_2H_4$, or Si. Note issues with charging, potential rough topography, and the stronger total ion signal to the bottom left of mapped area, indicating a non-level polished surface relative to the detector.*

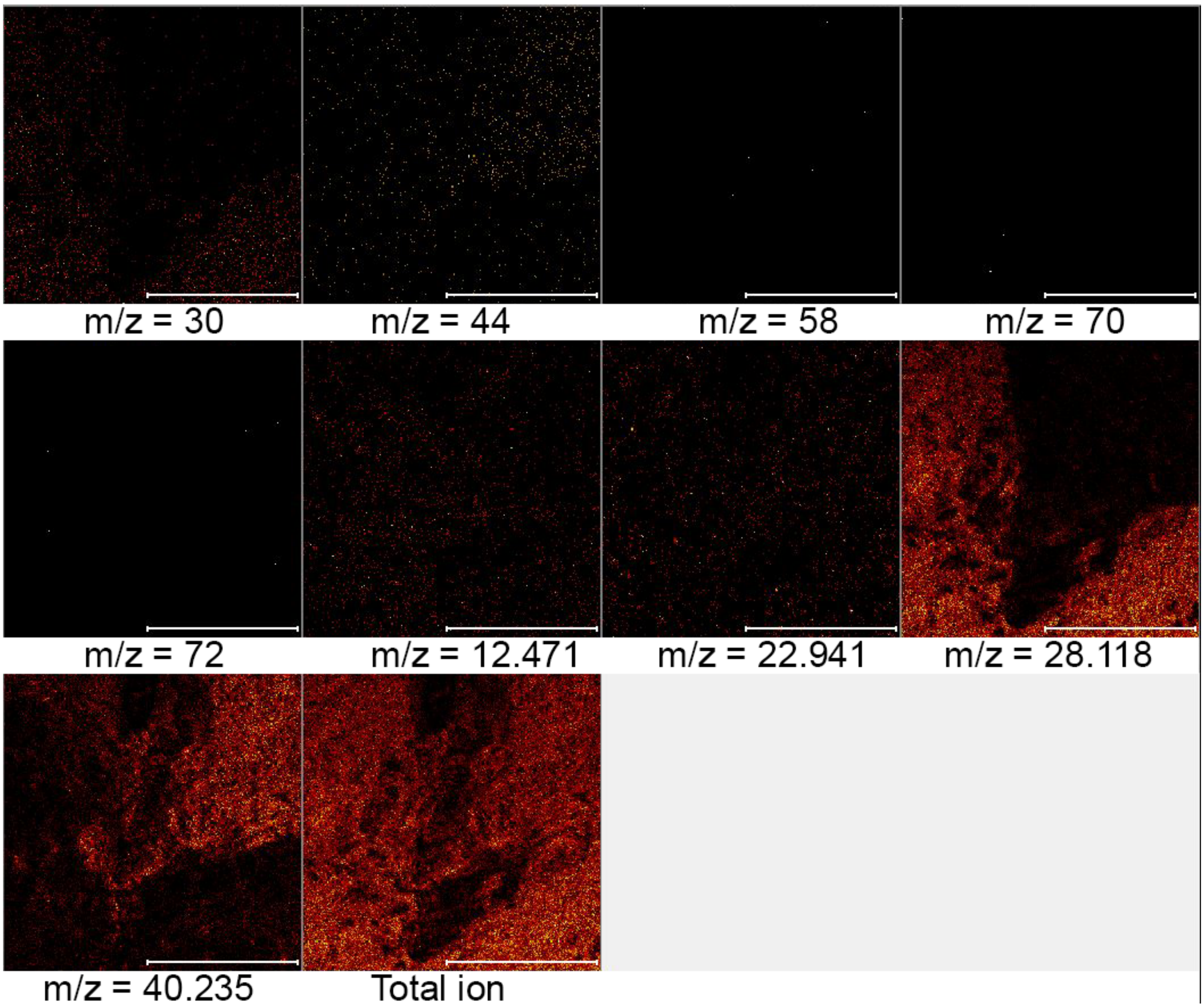


*Figure S.22. Positive TOF-SIMS mapping performed at Northwestern University on a second region of interest in the cross section of aseptically polished fragment of titanosaur eggshell A highlighting secondary ions of potential amino acid fragments from Hedberg* et al. *2012 (*m/z *= ~30 [Gly], ~44 [Ala], ~58 [Glu], ~70 [Val], ~72 [Val]), C (*m/z *= ~12), Na (*m/z *= ~23), Ca (*m/z *= ~40), and the total secondary ion map. The signal at* m/z *= ~28 could be consistent with either $CH_2N$, $C_2H_4$, or Si. Note issues with charging and potential rough topography.*

**TOF-SIMS Results Summary**

Attempts at TOF-SIMS on resin-embedded thin sections of eggshell yielded results that were dominated by a secondary ion signature of the embedding epoxy resin and were therefore not useful for examining the distribution of organic signatures. Further attempts using aseptically polished eggshell that were not embedded in a matrix were most often were dominated by inorganic secondary ions of calcite (and possibly silicate) and complicated by sample charging issues, potential topographic unevenness, and difficulty in levelling the polished surface relative to the detector. Calcite domination in the secondary ion signature may not be surprising given that TOF-SIMS typically examines the nanoscale surface of the sample, so any amino acids held within the calcite crystals might be obscured by the surrounding biomineral.

**RAMAN SPECTROSCOPY:** Raman spectroscopy was performed at the Field Museum of Natural History on a WITec Raman alpha 300R system using a 532 nm wavelength and a 600g/mm grating. Resin-embedded thin sections of ostrich and titanosaur A eggshell were examined (after light microscopy and TOF-SIMS) through point analysis and mapping.

Point spectra for the ostrich eggshell were obtained using a 20 mW laser power and 20 second integration time (Figs. S.23–24; tables S.9–11).

For the titanosaur A eggshell, one map was produced under low laser power (~100 µW, 5 second integration, 20 x 20 µm raster [Fig. 8B]) and another under slightly higher laser power (0.5 mW, 10 second integration, 20 x 20 µm raster [Fig. S.27]). For the point spectra (Figs. 8, S.25 and tables S.12–16), the darkly colored, recrystallized points of the titanosaur A eggshell were analyzed under high laser power (20 mW, 20 second integration). However, due to intense background fluorescence of the calcite, the lightly colored, non-recrystallized points of the titanosaur A eggshell were obtained directly from the low laser power map file in WITec software (~100 µW, 5 second integration, 20 x 20 µm raster), as opposed to a separate point acquisition.

Data then underwent cosmic ray correction, background correction, and smoothing in WITec software, as necessary. All spectra underwent cosmic ray removal and then background correction (shape, 150 size, noise factor 1). Titanosaur eggshell A low laser power map (~100 µW, 5 second integration, 20 x 20 µm raster) and the light phase points extracted from that map file were additionally smoothed (SG, parameters = 5,5,4,0) after background correction.

Due to the high amount of noise from calcite fluorescence, this raw/total spectral data was then analyzed in a manner that attempted to limit subjective interpretation and improve reproducibility. WITec software was first used to automatically label prominent peaks so as to avoid potentially interpreting noise as peaks (output of this shown in tables S.9–16). These chosen raw/total peak values were then verified or refined in FitYK software in order to fit more accurate peak positions and sometimes deconvolute peaks for subsequent interpretation of molecular structure (output of this shown in tables S.9–16). The automatic 'add peak' (Gaussian curve) and fitting functions in FitYK were used for this purpose (see captions of tables S.9–16 for details of how each spectrum was fitted/deconvoluted).

The fitted peak values from FitYK were then compared to reference vibrations in Lin-Vien *et al.* (1991) (table S.8) for organic molecules and the Handbook of Raman Spectra for Geology (Laboratoire de Géologie de Lyon, Université de Lyon) for calcite and quartz reference spectra. Peaks on the spectra in Figs. 8, S.23 were then labelled through color coded symbols based on whether they fell within the reported range of reference vibrations (or if they were within ~2 $cm^{-1}$ of a reference vibration listed as a single value, as opposed to a range) of organic molecules in Lin-Vien *et al.* (1991). Peaks that were close to the single values of inorganic vibrations for calcite or quartz listed in the Handbook of Raman Spectra for Geology (i.e., within ~3 $cm^{-1}$) were assumed to be inorganic, and this inorganic assignment was given priority over any potential organic matches (tables S.9–16) when labelling the spectra in Figs. 8, S.23.

Raw spectral data .txt files are available for the point analyses.

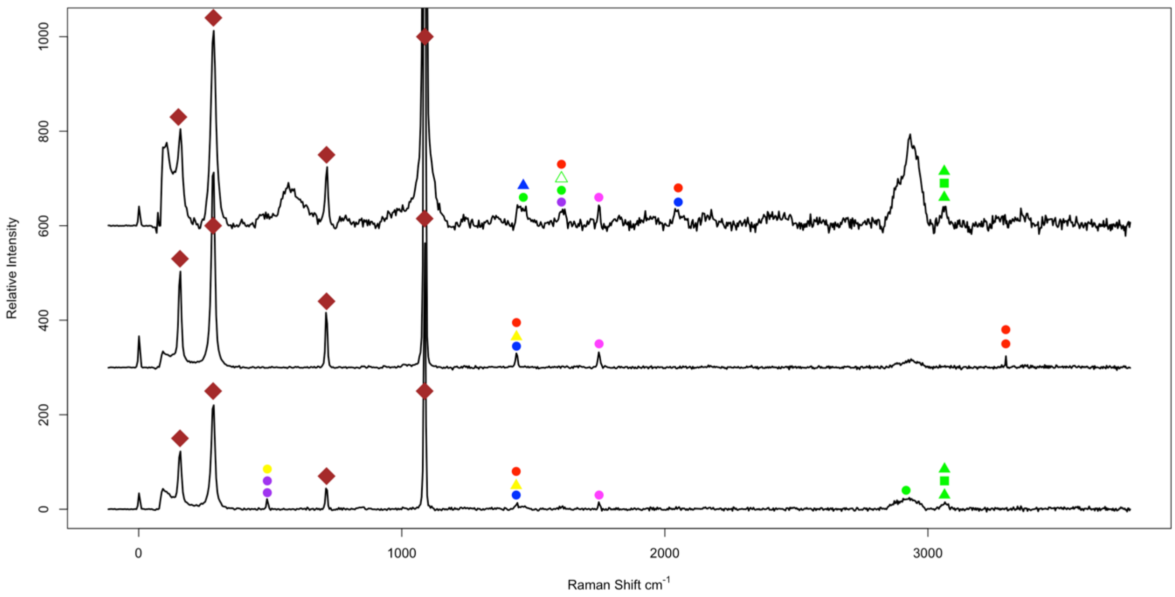


*Figure S.23. Ostrich eggshell Raman spectra taken from points in the outer prismatic external (top), center palisade/column (middle), and inner mammillary cone (bottom) layers of the resin-embedded thin section. Largest calcite peak is allowed to extend outside of the plot area in order to aid visual comparison of the smaller peaks. Peak labelling is the same as Fig. 8, but with magenta additionally indicating a match to a vibration from a compound containing both O and a halogen from Lin-Vien* et al. *(1991). Matches to halogen-bearing compounds might indicate the incorporation of environmental contaminants/pollutants from diet into the eggshell.*

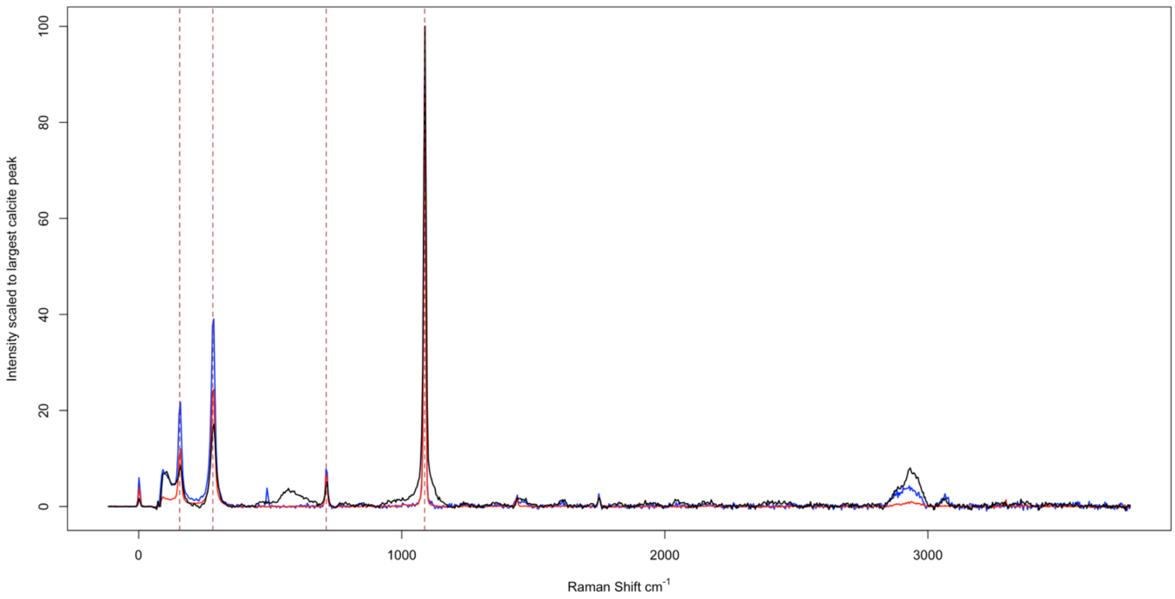


*Figure S.24. Ostrich eggshell resin-embedded thin section Raman spectra scaled to the height of the major calcite peak as a percentage. Reference calcite vibrations (from the Handbook of Raman Spectra for Geology, Laboratoire de Géologie de Lyon, Université de Lyon) are indicated by brown, dashed, vertical lines. Outer prismatic external (black), center palisade/column (red), and inner mammillary cone (blue) layers are indicated.*

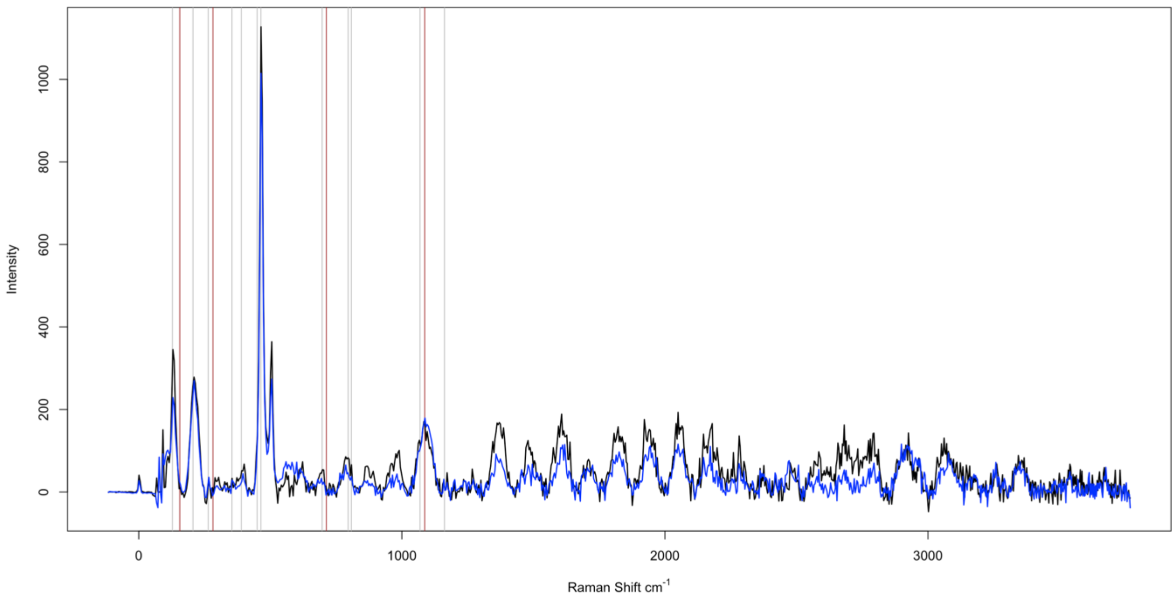


*Figure S.25. Raman spectra of points from the darkly colored/recrystallized regions of Titanosaur eggshell A resin-embedded thin section. These are shown vertically expanded compared to the presentation in Fig. 8. The first dark point is in black and measured from the dark region on the side closer to the outer eggshell surface relative to the central lightly colored, non-recrystallized region. The second dark point is in blue and measured from the dark region on the side closer to the inner eggshell surface relative to the central lightly colored, non-recrystallized region. Brown and grey lines indicate peak positions for the reference calcite and quartz spectra, respectively (from the Handbook of Raman Spectra for Geology, Laboratoire de Géologie de Lyon, Université de Lyon).*

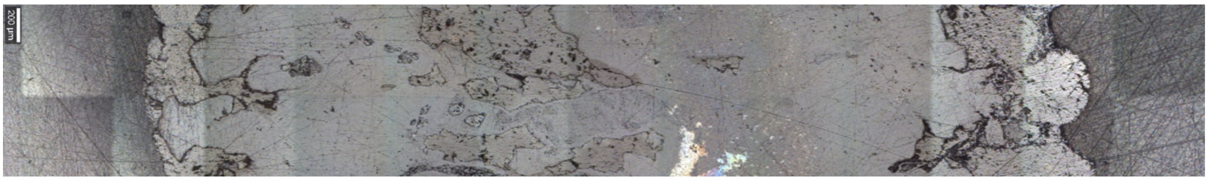

*Figure S.26. Reflected light image of the cross-sectional same area of titanosaur eggshell A resin-embedded thin section as seen in Fig. 8A. Outer surface of eggshell to the right. Inner surface of eggshell to the left. The two phases of the eggshell are discernible.*

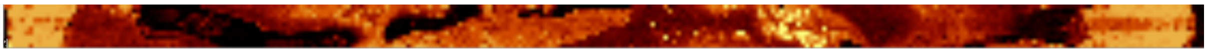

*Figure S.27. Expanded (i.e., excess resin cropped out from left and right edges) whole spectrum Raman spectroscopy map of the resin-embedded thin section of titanosaur eggshell A under slightly higher laser power (0.5 mW). Outer surface of eggshell to the right edge of image (bulb-shaped ornament visible). Inner surface of eggshell to the left edge of image. The two phases of the eggshell are still present, but not as clearly observed as in the map under low laser power in Fig. 8B. The dark phase is best examined under high laser power, but this leads to excessive noise in the light phase, which is best examined under low laser power.*

| Wavenumber | Bond | Compound | Ring? | Bands | Vibration |
|---|---|---|---|---|---|
| 3400-3330 | $NH_2$ | primary amines | | | bonded antisymmetric stretch |
| 3380-3340 | OH | aliphatic alcohols | | | bonded stretch |
| 3355-3325 | $NH_2$ | primary amides | | | bonded antisymmetric stretch |
| 3350-3300 | NH | secondary amines | | | bonded stretch |
| 3310-3290 | NH | secondary amides | | | bonded stretch |
| 3300-3250 | $NH_2$ | primary amines | | | bonded symmetric stretch |
| 3190-3145 | $NH_2$ | primary amides | | | bonded symmetric stretch |
| 3175-3154 | NH | pyrazoles | aromatic | | bonded stretch |
| 3100-3020 | $=CH_2$ | cyclopropane | cyclic | | stretches |
| 3100-3000 | CH | benzene derivatives | aromatic | | aromatic stretch |
| 3095-3070 | $=CH_2$ | $C=CH_2$ derivatives | | | antisymmetric stretch |
| 3062 | CH | benzene | aromatic | | stretch |
| 3040-3000 | CH | C=CHR derivatives | | | stretch |
| 3026 | $=CH_2$ | ethylene (gas) | | | symmetric stretch |
| 2929-2912 | $=CH_2$ | n-alkanes | | | antisymmetric stretch |
| 2850-2700 | CHO group | aliphatic aldehydes | | 2 | |
| 2590-2560 | SH | thiols | | | stretch |
| 2316-2233 | C≡C | $R–C≡C–CH_3$ | | 2 | stretch |
| 2301-2231 | C≡C | R–C≡C–R' | | 2 | stretch |
| 2300-2250 | N=C=O | isocyanates | | | pseudoantisymmetric stretch |
| 2264-2251 | C≡C–C≡C | alkyl diacetylenes | | | symmetric stretch |
| 2259 | C≡N | cyanamide | | | stretch |
| 2251-2232 | C≡N | aliphatic nitriles | | | stretch |
| 2220-2100 | N=C=S | alkyl isothiocyanates | | 2 | pseudoantisymmetric stretch |
| 2220-2000 | C≡N | dialkyl cyanamides | | | stretch |
| 2172 | C≡C–C≡C | diacetylene | | | symmetric stretch |
| 2161-2134 | N+≡C- | aliphatic isonitriles | | | stretch |
| 2160-2100 | C≡C | alkyl acetylenes | | | stretch |
| 2156-2140 | C≡N | alkyl thiocyanates | | | stretch |
| 2049 | C=C=O | ketene | | | pseudoantisymmetric stretch |
| 1820 | C=O | acetic anhydride | | | symmetric stretch |
| 1770-1730 | C=O | halogenated aldehydes | | | stretch |
| 1739-1714 | C=C | $C=CF_2$ derivatives | | | stretch |
| 1725-1700 | C=O | aliphatic ketones | | | stretch |
| 1720-1715 | C=O | *O*-alkyl formates | | | stretch |
| 1712-1694 | C=C | RCF=CFR | | | stretch |
| 1689-1644 | C=C | monofluoroalkanes | | | stretch |
| 1686-1636 | amide I | primary amides (solid) | | | |
| 1670-1630 | amide I | tertiary amides | | | |
| 1663-1636 | C=N | aldazines, ketazines | | | symmetric stretch |
| 1660-1610 | C=N | hydrazones (solid) | | | stretch |
| 1658-1644 | C=C | $R_2C=CH_2$ | | | stretch |
| 1652-1642 | C=N | thiosemicarbazones (solid) | | | stretch |
| 1650-1590 | $NH_2$ | primary amines | | | scissors |
| 1649-1625 | C=C | allyl derivatives | | | stretch |
| 1648-1640 | N=O | alkyl nitrites | | | stretch |
| 1648-1638 | C=C | $H_2C=CHR$ | | | stretch |
| 1647 | C=C | cyclopropene | cyclic | | stretch |
| 1630-1550 | benzene | benzene derivatives | aromatic | doublet | ring stretches |
| 1620-1540 | C=C | polyenes | | | three or more coupled stretches |
| 1616-1571 | C=C | chloroalkenes | | | stretch |
| 1596-1547 | C=C | bromoalkenes | | | stretch |
| 1581-1465 | C=C | iodoalkenes | | | stretch |
| 1515-1490 | 2-furfuryl group | 2-furfuryl group | aromatic | | ring stretch |
| 1480-1460 | 2 furfurylidene or 2-furoyl group | 2 furfurylidene or 2-furoyl group | aromatic | | ring stretch |
| 1473-1446 | $CH_3$, $CH_2$ | n-alkanes | | | deformations |
| 1466-1465 | $CH_3$ | n-alkanes | | | deformation |
| 1450-1400 | N=C=O | isocyanates | | | pseudosymmetric stretch |
| 1443-1398 | 2-substituted thiophene | 2-substituted thiophene | aromatic | | ring stretch |
| 1440-1340 | $CO_2$- | carboxylate ions (aqueous solution) | | | symmetric stretch |
| 1395-1380 | $NO_2$ | primary nitroalkanes | | | symmetric stretch |
| 1390-1370 | naphthalenes | naphthalenes | aromatic | | ring stretch |
| 1385-1368 | $CH_3$ | n-alkanes | | | symmetric deformation |
| 1375-1360 | $NO_2$ | secondary nitroalkanes | | | symmetric stretch |
| 1310-1250 | amide III | secondary amides | | | |
| 1310-1175 | $CH_2$ | n-alkanes | | | twist and rock |
| 1282-1275 | $NO_2$ | alkyl nitrates | | | symmetric stretch |
| 1280-1240 | epoxy derivatives | epoxy derivatives | cyclic | | ring stretch |
| 1276 | N=N=N | $CH_3N_3$ | | | symmetric stretch |
| 1270-1251 | CH | *cis*-dialkyl ethylenes | | | in-plane deformation |
| 1266 | oxirane | ethylene oxide | cyclic | | ring "breathing" |
| 1172-1165 | $SO_2$ | alkyl sulfonates | | | symmetric stretch |
| 1150-950 | CC | n-alkanes | | | stretches |
| 1000-985 | 2- and 4-substituted pyridines | 2- and 4-substituted pyridines | aromatic | | trigonal ring "breathing" |
| 930-830 | COC | aliphatic ethers | | | symmetric stretch |
| 905-837 | CC | n-alkanes | | | skeletal stretch |
| 900-850 | CNC | secondary amines | | | symmetric stretch |

| | | | | | |
|---|---|---|---|---|---|
| 866 | cyclopentane | cyclopentane | cyclic | | ring "breathing" |
| 835-749 | isopropyl group | isopropyl group | | | skeletal stretch |
| 830-720 | benzene | para-disubstituted benzenes | aromatic | | ring vibration |
| 785-700 | alkyl cyclohexanes | alkyl cyclohexanes | cyclic | | ring vibration |
| 760-650 | *tert*-butyl group | *tert*-butyl group | | | symmetric skeletal stretch |
| 740-585 | CS | alkyl sulfides | | 1 or more | stretch |
| 735-690 | C=S | thioamides, thioureas (solid) | | | "stretch" |
| 715-620 | CS | dialkyl disulfides | | 1 or more | stretch |
| 703 | cyclooctane | cyclooctane | cyclic | | ring "breathing" |
| 703 | $CCl_2$ | $CH_2Cl_2$ | | | symmetric stretch |
| 690-650 | N=C=S | alkyl isothiocyanates | | | pseudosymmetric stretch |
| 630-615 | benzene | monosubstituted benzenes | aromatic | | ring deformation |
| 577 | $CBr_2$ | $CH_2Br_2$ | | | symmetric stretch |
| 525-510 | SS | dialkyl disulfides | | | stretch |
| 520-510 | CBr | tertiary bromoalkanes | | | stretch, THHH conformation |
| 510-500 | CI | primary iodoalkanes | | | stretch, PH conformation |
| 510-480 | SS | dialkyl trisulfides | | | stretch |
| 495-485 | CI | secondary iodoalkanes | | | stretch, SHH conformation |
| 495-485 | CI | tertiary iodoalkanes | | | stretch, THHH conformation |
| 484-475 | dialkyl diacetylenes | dialkyl diacetylenes | | | skeletal deformation |
| 425-150 | n-alkanes | n-alkanes | | | "chain expansion" |

*Table S.8 (extends from previous page). Potentially relevant Raman reference vibrations for organic molecules reproduced from Lin-Vien* et al. *(1991) based on potential matches to our titanosaur eggshell A and ostrich resin-embedded thin sections. Color coded as in Figs. 8, S.21. "Cyclic" indicates non-aromatic cyclic compounds. References with "1 or more" bands were coded as single bands (i.e., solid symbols) in Figs. 8, S.23 and Tables S.9–16, as opposed to doublets/2 bands.*

| **Raw Peak (WITec)** | 3 | 102 | 102 | 158 | 284 | 570 | 714 | 1088 | 1449 | 1611 | 1749 | 2045 | 2932 | 3063 |
|---|---|---|---|---|---|---|---|---|---|---|---|---|---|---|
| **Fitted Peak (FitYK)** | **2** | **93** | **109** | **151** | **284** | **583** | **714** | **1088** | **1462** | **1607** | **1749** | **2051** | **2940** | **3062** |
| **Matches** | | | | 425-150 | 425-150 | | 715-620 | 1150-950 | 1473-1446 | 1616-1571 | 1770-1730 | 2049 | | 3062 A |
| C = cyclic | | | | | | | 735-690 | | 1480-1460 A | 1620-1540 | | 2220-2000 | | 3100-3000 A |
| A = aromatic | | | | | | | 740-585 | | | **1630-1550 AD** | | | | 3100-3020 C |
| **D = doublet** | | | | | | | 760-650 | | | 1650-1590 | | | | |
| | | | | | | | 785-700 C | | | | | | | |

*Table S.9. Raw/total Raman peak values obtained from WITec, fitted peaks generated using FitYK, and potential matches to inorganic (from the Handbook of Raman Spectra for Geology, Laboratoire de Géologie de Lyon, Université de Lyon) and organic peaks (Lin-Vien* et al. *1991) for the outer/prismatic external layer of the ostrich resin-embedded thin section. Color coding is the same as in Figs. 8, S.23. Letter coding indicates ring type and whether or not the potential match is listed by Lin-Vien* et al. *(1991) as a doublet. Close matches to calcite or quartz (indicated by brown shading of the fitted peak value) were interpreted as taking priority over any potential organic peaks, and these peaks were subsequently solely labelled as inorganic in the spectra of Figs. 8, S.23. To convert observed raw/total peak values in WITec to deconvoluted/fitted peak values in FitYK, 21 Gaussian curves were generated using the automatic peak addition command once, followed by the fit command, and then repeating this process 21 times.*

| Raw Peak (WITec) | 3 | 96 | 157 | 283 | 714 | 1087 | 1437 | 1750 | 2937 | 3295 |
|---|---|---|---|---|---|---|---|---|---|---|
| **Fitted Peak (FitYK)** | **2** | **97** | **157** | **283** | **714** | **1087** | **1436** | **1750** | **2930** | **3296** |
| **Matches** | | | 425-150 | 425-150 | 715-620 | 1150-950 | 1440-1340 | 1770-1730 | | 3310-3290 |
| C = cyclic | | | | | 735-690 | | 1443-1398 A | | | 3300-3250 |
| A = aromatic | | | | | 740-585 | | 1450-1400 | | | |
| **D = doublet** | | | | | 760-650 | | | | | |
| | | | | | 785-700 C | | | | | |

*Table S.10. Raw/total Raman peak values obtained from WITec, fitted peaks generated using FitYK, and potential matches to inorganic (from the Handbook of Raman Spectra for Geology, Laboratoire de Géologie de Lyon, Université de Lyon) and organic peaks (Lin-Vien* et al. *1991) for the center palisade/column layer of the ostrich resin-embedded thin section. Color coding is the same as in Figs. 8, S.23. Letter coding indicates ring type and whether or not the potential match is listed by Lin-Vien* et al. *(1991) as a doublet. Close matches to calcite or quartz (indicated by brown shading of the fitted peak value) were interpreted as taking priority over any potential organic peaks, and these peaks were subsequently solely labelled as inorganic in the spectra of Figs. 8, S.23. To convert observed raw/total peak values in WITec to deconvoluted/fitted peak values in FitYK, 34 Gaussian curves were generated using the automatic peak addition command once, followed by the fit command, and then repeating this process 34 times.*

| Raw Peak (WITec) | 3 | 96 | 96 | 157 | 283 | 489 | 714 | 1087 | 1438 | 1750 | 2929 | 3065 |
|---|---|---|---|---|---|---|---|---|---|---|---|---|
| **Fitted Peak (FitYK)** | **2** | **90** | **107** | **157** | **283** | **489** | **714** | **1087** | **1435** | **1750** | **2917** | **3063** |
| **Matches** | | | | 425-150 | 425-150 | 495-485 | 715-620 | 1150-950 | 1440-1340 | 1770-1730 | 2929-2912 | 3062 A |
| C = cyclic | | | | | | 495-485 | 735-690 | | 1443-1398 A | | | 3100-3000 A |
| A = aromatic | | | | | | 510-480 | 740-585 | | 1450-1400 | | | 3100-3020 C |
| **D = doublet** | | | | | | | 760-650 | | | | | |
| | | | | | | | 785-700 C | | | | | |

*Table S.11. Raw/total Raman peak values obtained from WITec, fitted peaks generated using FitYK, and potential matches to inorganic (from the Handbook of Raman Spectra for Geology, Laboratoire de Géologie de Lyon, Université de Lyon) and organic peaks (Lin-Vien* et al. *1991) for the inner/mammillary cone layer of the ostrich resin-embedded thin section. Color coding is the same as in Figs. 8, S.23. Letter coding indicates ring type and whether or not the potential match is listed by Lin-Vien* et al. *(1991) as a doublet. Close matches to calcite or quartz (indicated by brown shading of the fitted peak value) were interpreted as taking priority over any potential organic peaks, and these peaks were subsequently solely labelled as inorganic in the spectra of Figs. 8, S.23. To convert observed raw/total peak values in WITec to deconvoluted/fitted peak values in FitYK, 25 Gaussian curves were generated using the automatic peak addition command once, followed by the fit command, and then repeating this process 25 times.*

| Raw Peak | Fitted Peak | Matches | C = cyclic | A = aromatic | D = doublet | | | |
|---|---|---|---|---|---|---|---|---|
| 104 | | | | | | | | |
| 295 | 297 | 425-150 | | | | | | |
| 391 | 391 | 425-150 | | | | | | |
| 505 | 512 | 520-510 | 525-510 | | | | | |
| 619 | 620 | 630-615 A | 715-620 | | | | | |
| 701 | 701 | 703 C | 703 | 715-620 | 735-690 | 740-585 | 760-650 | 785-700 C |
| 793 | 810 | 830-720 A | 835-749 | | | | | |
| 971 | 970 | 1150-950 | | | | | | |
| 1166 | 1168 | 1172-1165 | | | | | | |
| 1264 | 1264 | 1266 C | 1270-1251 | 1280-1240 C | 1310-1175 | 1310-1250 | | |
| 1375 | 1381 | 1385-1368 | 1390-1370 A | 1395-1380 | 1440-1340 | | | |
| 1482 | 1466 | 1466-1465 | 1473-1446 | 1480-1460 A | 1581-1465 | | | |
| | 1494 | 1515-1490 | 1581-1465 | | | | | |
| 1600 | 1594 | 1596-1547 | 1616-1571 | 1620-1540 | **1630-1550 AD** | 1650-1590 | | |
| 1701 | 1703 | 1712-1694 | 1725-1700 | | | | | |
| 1822 | 1819 | 1820 | | | | | | |
| 1925 | 1930 | | | | | | | |
| 2040 | 2039 | 2220-2000 | | | | | | |
| 2145 | 2149 | 2156-2140 | 2160-2100 | 2161-2134 | 2220-2000 | **2220-2100 D** | | |
| 2250 | 2249 | 2251-2232 | **2301-2231 D** | **2316-2233 D** | | | | |
| 2359 | 2354 | | | | | | | |
| 2469 | 2469 | | | | | | | |
| 2586 | 2586 | 2590-2560 | | | | | | |
| 2711 | 2717 | **2850-2700 D** | | | | | | |
| 2804 | 2808 | **2850-2700 D** | | | | | | |
| 2925 | 2916 | 2929-2912 | | | | | | |
| 3037 | 3026 | 3026 | 3040-3000 | 3100-3000 A | 3100-3020 C | | | |
| | 3049 | 3100-3000 A | 3100-3020 C | | | | | |
| 3107 | 3106 | | | | | | | |
| 3340 | 3347 | 3350-3300 | 3355-3325 | 3380-3340 | 3400-3330 | | | |
| 3443 | 3442 | | | | | | | |
| 3470 | 3471 | | | | | | | |
| 3573 | 3569 | | | | | | | |
| 3678 | 3675 | | | | | | | |

*Table S.12. Raw/total Raman peak values obtained from WITec, fitted peaks generated using FitYK, and potential matches to inorganic (from the Handbook of Raman Spectra for Geology, Laboratoire de Géologie de Lyon, Université de Lyon) and organic peaks (Lin-Vien* et al. *1991) in the outer lightly colored region/external ornament in titanosaur eggshell A resin-embedded thin section. Color coding is the same as in Figs. 8, S.23. Letter coding indicates ring type and whether or not the potential match is listed by Lin-Vien* et al. *(1991) as a doublet. Close matches to calcite or quartz (indicated by brown or grey shading of the fitted peak value) were interpreted as taking priority over any potential organic peaks, and these peaks were subsequently solely labelled as inorganic in the spectra of Figs. 8, S.23. To convert observed raw/total peak values in WITec to deconvoluted/fitted peak values in FitYK, 57 Gaussian curves were generated using the automatic peak addition command once, followed by the fit command, and then repeating this process 57 times.*

| Raw Peak | Fitted Peak | Matches | C = cyclic | A = aromatic | D = doublet | |
|---|---|---|---|---|---|---|
| 3 | **2** | | | | | |
| 132 | **131** | | | | | |
| 212 | **211** | 425-150 | | | | |
| 397 | **395** | 425-150 | | | | |
| 466 | **465** | | | | | |
| 504 | **504** | 510-480 | 510-500 | | | |
| 560 | **575** | 577 | | | | |
| 619 | | | | | | |
| 694 | **682** | 690-650 | 715-620 | 740-585 | 760-650 | |
| 791 | **786** | 830-720 A | 835-749 | | | |
| 868 | **865** | 866 C | 900-850 | 905-837 | 930-830 | |
| 983 | **974** | 1150-950 | | | | |
| 1086 | **1090** | 1150-950 | | | | |
| 1368 | **1369** | 1375-1360 | 1385-1368 | 1440-1340 | | |
| 1482 | **1488** | 1581-1465 | | | | |
| 1607 | **1610** | 1616-1571 | 1620-1540 | **1630-1550 AD** | 1650-1590 | 1660-1610 |
| 1710 | **1716** | 1720-1715 | 1725-1700 | 1739-1714 | | |
| 1818 | **1824** | | | | | |
| 1827 | | | | | | |
| 1931 | **1943** | | | | | |
| 2054 | **2050** | 2049 | 2220-2000 | | | |
| 2173 | **2171** | 2172 | 2220-2000 | **2220-2100 D** | | |
| 2284 | **2283** | 2300-2250 | **2301-2231 D** | **2316-2233 D** | | |
| 2363 | | | | | | |
| 2485 | **2478** | | | | | |
| 2584 | **2581** | 2590-2560 | | | | |
| 2792 | **2786** | **2850-2700 D** | | | | |
| 2923 | **2919** | 2929-2912 | | | | |
| 2967 | **2951** | | | | | |
| 3062 | **3079** | 3095-3070 | 3100-3000 A | 3100-3020 C | | |
| 3259 | **3258** | 3300-3250 | | | | |
| 3346 | **3349** | 3350-3300 | 3355-3325 | 3380-3340 | 3400-3330 | |
| 3356 | | | | | | |

*Table S.13. Raw/total Raman peak values obtained from WITec, fitted peaks generated using FitYK, and potential matches to inorganic (from the Handbook of Raman Spectra for Geology, Laboratoire de Géologie de Lyon, Université de Lyon) and organic peaks (Lin-Vien* et al. *1991) for the first darkly colored/recrystallized point in titanosaur eggshell A resin-embedded thin section on the side closer to the external surface of the eggshell relative to the central lightly colored region. Color coding is the same as in Figs. 8, S.23. Letter coding indicates ring type and whether or not the potential match is listed by Lin-Vien* et al. *(1991) as a doublet. Close matches to calcite or quartz (indicated by brown or grey shading of the fitted peak value) were interpreted as taking priority over any potential organic peaks, and these peaks were subsequently solely labelled as inorganic in the spectra of Figs. 8, S.23. To convert observed raw/total peak values in WITec to deconvoluted/fitted peak values in FitYK, 57 Gaussian curves were generated using the automatic peak addition command in succession, followed by the fit command in order to fit all 57 peaks at once, deleting two peaks with negative heights, and then re-fitting the remaining 55 peaks at once using the fit command.*

| **Raw Peak (WITec)** | 792 | 867 | 963 | 1273 | 1491 | 1652 | 1817 | 1850 | 1920 | 2044 | 2166 | 2253 | 2466 | 2692 |
|---|---|---|---|---|---|---|---|---|---|---|---|---|---|---|
| **Fitted Peak (FitYK)** | **793** | **867** | **969** | **1275** | **1487** | **1648** | **1823** | | **1931** | **2046** | **2158** | **2266** | **2465** | **2691** |
| **Matches** | 830-720 A | 866 C | 1150 -950 | 1276 | 1581-1465 | 1647 C | | | | 2220-2000 | 2160-2100 | 2300-2250 | | |
| C = cyclic | 835-749 | 900-850 | | 1280-1240 C | | 1648-1638 | | | | | 2161-2134 | **2301-2231 D** | | |
| A = aromatic | | 905-837 | | 1282-1275 | | 1648-1640 | | | | | 2220-2000 | **2316-2233 D** | | |
| **D = doublet** | | 930-830 | | 1310-1175 | | 1649-1625 | | | | | **2220-2100 D** | | | |
| | | | | 1310-1250 | | 1650-1590 | | | | | | | | |
| | | | | | | 1652-1642 | | | | | | | | |
| | | | | | | 1658-1644 | | | | | | | | |
| | | | | | | 1660-1610 | | | | | | | | |
| | | | | | | 1663-1636 | | | | | | | | |
| | | | | | | 1670-1630 | | | | | | | | |
| | | | | | | 1686-1636 | | | | | | | | |
| | | | | | | 1689-1644 | | | | | | | | |

*Table S.14. Raw/total Raman peak values obtained from WITec, fitted peaks generated using FitYK, and potential matches to inorganic (from the Handbook of Raman Spectra for Geology, Laboratoire de Géologie de Lyon, Université de Lyon) and organic peaks (Lin-Vien* et al. *1991) in the center lightly colored region in titanosaur eggshell A resin-embedded thin section. Color coding is the same as in Figs. 8, S.23. Letter coding indicates ring type and whether or not the potential match is listed by Lin-Vien* et al. *(1991) as a doublet. Close matches to calcite or quartz (indicated by brown or grey shading of the fitted peak value) were interpreted as taking priority over any potential organic peaks, and these peaks were subsequently solely labelled as inorganic in the spectra of Figs. 8, S.23. To convert observed raw/total peak values in WITec to deconvoluted/fitted peak values in FitYK, 57 Gaussian curves were generated using the automatic peak addition command in succession, followed by the fit command in order to fit all 57 peaks at once, deleting one extremely tall outlier peak, and then re-fitting the remaining 56 peaks at once using the fit command.*

| | D = doublet | A = aromatic | C = cyclic | Matches | Fitted Peak | Raw Peak |
|---|---|---|---|---|---|---|
| | | | | | **2** | 2 |
| | | | | | **131** | 132 |
| | | | | 425-150 | **211** | 210 |
| | | | | 425-150 | **395** | 396 |
| | | | | | **465** | 466 |
| | | | 510-500 | 510-480 | **504** | 504 |
| | | | | 577 | **575** | 566 |
| | 760-650 | 740-585 | 715-620 | 690-650 | **682** | 692 |
| | | | 835-749 | 830-720 A | **786** | 788 |
| | 930-830 | 905-837 | 900-850 | 866 C | **865** | 865 |
| | | | 1150-950 | 1000-985 A | **974** | 967 |
| | | | | | | 978 |
| | | | | 1150-950 | **1090** | 1091 |
| | | 1440-1340 | 1385-1368 | 1375-1360 | **1369** | 1368 |
| 1660-1610 | 1650-1590 | **1630-1550 AD** | 1620-1540 | 1616-1571 | **1610** | 1610 |
| | | 1739-1714 | 1725-1700 | 1720-1715 | **1716** | 1719 |
| | | | | | **1824** | 1824 |
| | | | | | **1943** | 1943 |
| | | | 2220-2000 | 2049 | **2050** | 2049 |
| | | **2220-2100 D** | 2220-2000 | 2172 | **2171** | 2170 |
| **2316-2233 D** | **2301-2231 D** | 2300-2250 D | 2264-2251 | 2259 | **2257** | 2258 |
| | | **2316-2233 D** | **2301-2231 D** | 2300-2250 | **2283** | 2285 |
| | | | | | | 2372 |
| | | | | | **2421** | 2423 |
| | | | | | **2478** | 2472 |
| | | | | | **2506** | 2501 |
| | | | | | **2546** | 2547 |
| | | | | 2590-2560 | **2581** | 2582 |
| | | | | | **2684** | 2684 |
| | | | | **2850-2700 D** | **2786** | 2790 |
| | | | | 2929-2912 | **2919** | 2926 |
| | | 3100-3020 C | 3100-3000 A | 3095-3070 | **3079** | 3082 |
| | | | 3190-3145 | 3175-3154 A | **3175** | 3178 |
| | | | | 3300-3250 | **3258** | 3255 |
| | 3400-3330 | 3380-3340 | 3355-3325 | 3350-3300 | **3349** | 3350 |
| | | | | | **3538** | 3538 |
| | | | | | **3675** | 3674 |

*Table S.15. Raw/total Raman peak values obtained from WITec, fitted peaks generated using FitYK, and potential matches to inorganic (from the Handbook of Raman Spectra for Geology, Laboratoire de Géologie de Lyon, Université de Lyon) and organic peaks (Lin-Vien* et al. *1991) for the second darkly colored/recrystallized point in titanosaur eggshell A resin-embedded thin section on the side closer to the internal surface of the eggshell relative to the central lightly colored region. Color coding is the same as in Figs. 8, S.23. Letter coding indicates ring type and whether or not the potential match is listed by Lin-Vien* et al. *(1991) as a doublet. Close matches to calcite or quartz (indicated by brown or grey shading of the fitted peak value) were interpreted as taking priority over any potential organic peaks, and these peaks were subsequently solely labelled as inorganic in the spectra of Figs. 8, S.23. To convert observed raw/total peak values in WITec to deconvoluted/fitted peak values in FitYK, 57 Gaussian curves were generated using the automatic peak addition command in succession, followed by the fit command in order to fit all 57 peaks at once, and then deleting two peaks with negative heights to leave 55 peaks.*

| **D = doublet** | A = aromatic | C = cyclic | **Matches** | **Fitted Peak** | **Raw Peak** |
|---|---|---|---|---|---|
| | | | | **136** | 137 |
| | | | 425-150 | **311** | 301 |
| | | | 425-150 | **390** | 390 |
| | | | 484-475 | **476** | 474 |
| 760-650 | 740-585 | 735-690 | 715-620 | **694** | 692 |
| | | 835-749 | 830-720 A | **790** | 789 |
| | 930-830 | 905-837 | 900-850 | **861** | 864 |
| | | | 1150-950 | **972** | 964 |
| | | | 1150-950 | **1074** | 1093 |
| | | 1440-1340 | 1375-1360 | **1364** | 1369 |
| | | 1581-1465 | 1515-1490 A | **1491** | 1499 |
| 1650-1590 | **1630-1550 AD** | 1620-1540 | 1616-1571 | **1608** | 1601 |
| | | | | | 1700 |
| | | | | **1816** | 1823 |
| | | | | **1950** | 1935 |
| | | 2220-2000 | 2049 | **2049** | 2055 |
| **2220-2100 D** | 2220-2000 | 2161-2134 | 2160-2100 | **2160** | 2166 |
| | | | | | 2284 |
| | | | | **2676** | 2695 |
| | | | **2850-2700 D** | **2782** | 2784 |
| | | | 2929-2965 | **2927** | 2927 |
| | | 3100-3000 C | 3040-3000 | **3014** | 3014 |
| | | 3100-3020 C | 3100-3000 A | **3048** | 3051 |
| 3400-3330 | 3380-3340 | 3355-3325 | 3350-3300 | **3345** | 3344 |

*Table S.16. Raw/total Raman peak values obtained from WITec, fitted peaks generated using FitYK, and potential matches to inorganic (from the Handbook of Raman Spectra for Geology, Laboratoire de Géologie de Lyon, Université de Lyon) and organic peaks (Lin-Vien* et al. *1991) in the inner lightly colored region in titanosaur eggshell A resin-embedded thin section. Color coding is the same as in Figs. 8, S.23. Letter coding indicates ring type and whether or not the potential match is listed by Lin-Vien* et al. *(1991) as a doublet. Close matches to calcite or quartz (indicated by brown or grey shading of the fitted peak value) were interpreted as taking priority over any potential organic peaks, and these peaks were subsequently solely labelled as inorganic in the spectra of Figs. 8, S.23. To convert observed raw/total peak values in WITec to deconvoluted/fitted peak values in FitYK, 57 Gaussian curves were generated using the automatic peak addition command in succession, followed by the fit command in order to fit all 57 peaks at once, deleting one peak with a negative height, and then re-fitting the remaining 56 peaks at once using the fit command.*

**ADDITIONAL EGGSHELL MICROGRAPHS:** Flakes of outer eggshell that separated naturally from titanosaur eggshell A during the sampling process were analysed separately.

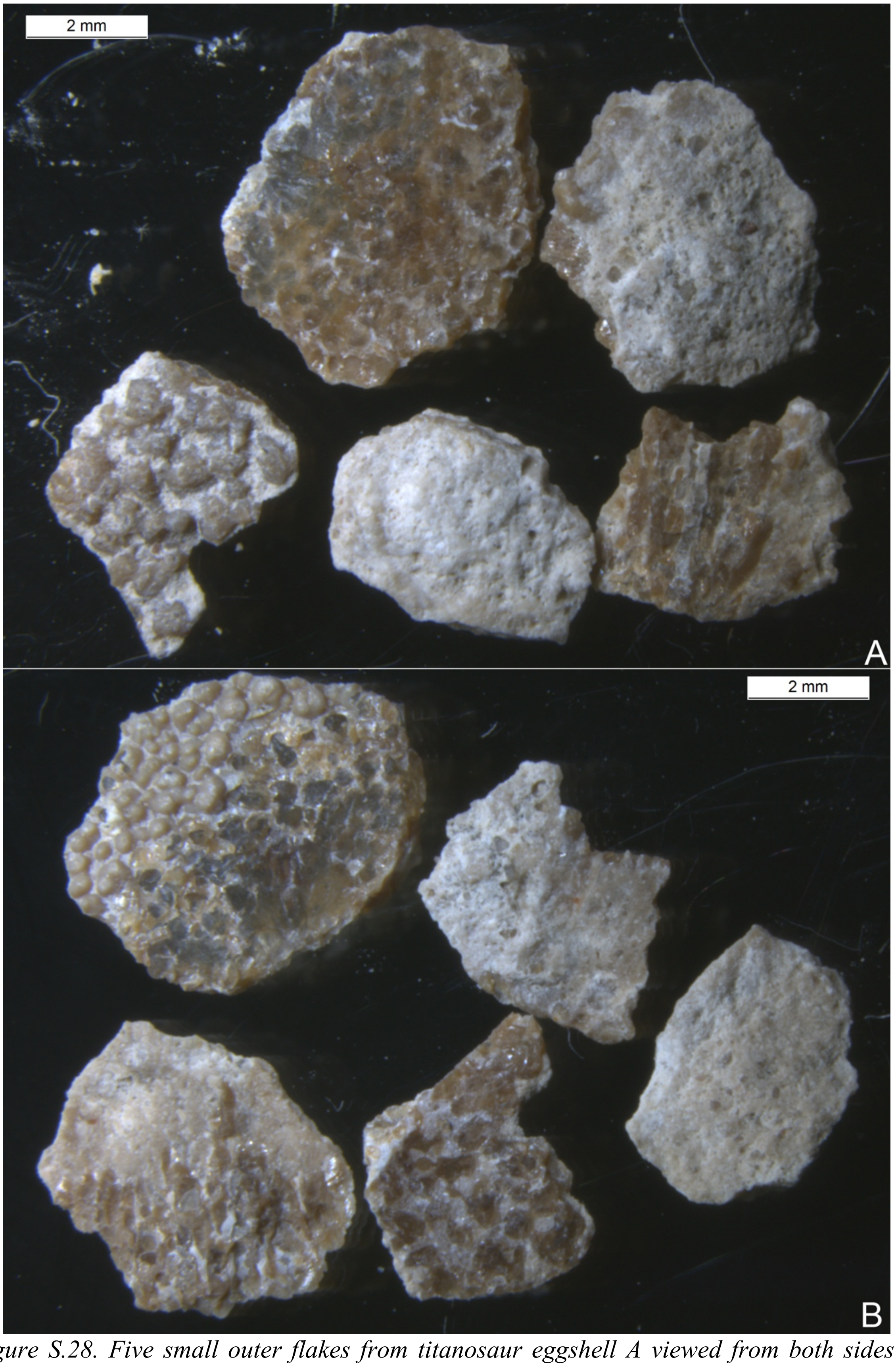


*Figure S.28. Five small outer flakes from titanosaur eggshell A viewed from both sides in panels A and B.*

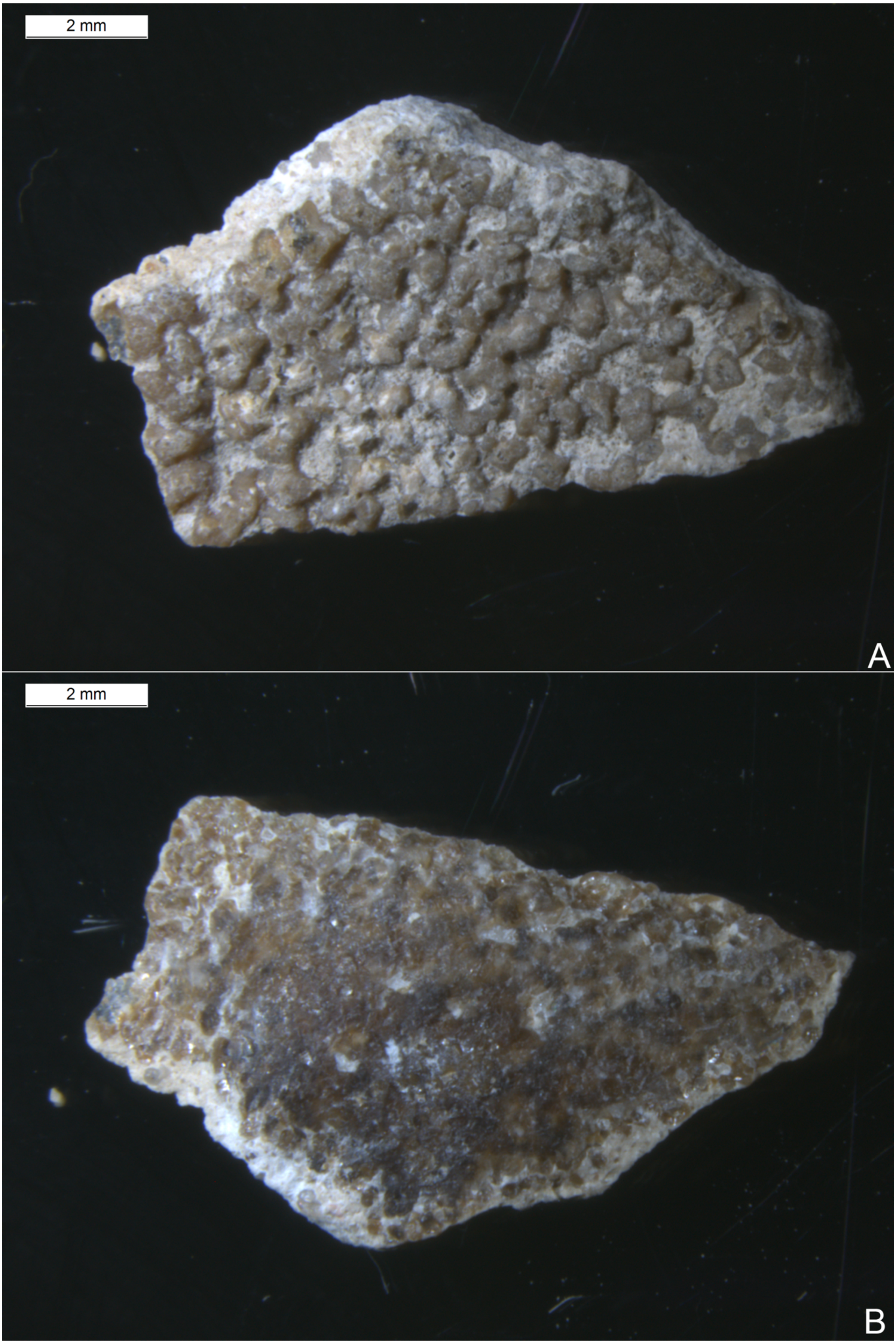

*Figure S.29. Large outer flake from titanosaur eggshell A viewed from both sides in panels A and B.*
